# New genome assemblies reveal patterns of domestication and adaptation across *Brettanomyces* (*Dekkera*) species

**DOI:** 10.1101/805721

**Authors:** Michael J. Roach, Anthony R. Borneman

**Affiliations:** The Australian Wine Research Institute, PO Box 197, Glen Osmond, South Australia, 5046. Australia

**Keywords:** *Brettanomyces*, genome comparison, diploid assembly, wine, yeast

## Abstract

**Background:** Yeast of the genus *Brettanomyces* are of significant interest, both for their capacity to spoil, as well as their potential to positively contribute to different industrial fermentations. However, considerable variance exists in the depth of research and knowledgebase of the five currently known species of *Brettanomyces*. For instance, *Brettanomyces bruxellensis* has been heavily studied and many resources are available for this species, whereas *Brettanomyces nanus* is rarely studied and lacks a publicly available genome assembly altogether. The purpose of this study is to fill this knowledge gap and explore the genomic adaptations that have shaped the evolution of this genus.

**Results:** Strains for each of the five widely accepted species of *Brettanomyces* (*Brettanomyces anomalus*, *B. bruxellensis*, *Brettanomyces custersianus*, *Brettanomyces naardenensis*, and *B. nanus*) were sequenced using a combination of long- and short-read sequencing technologies. Highly contiguous assemblies were produced for each species. Sweeping and extensive structural variation between the species’ genomes were observed and gene expansions in fermentation-relevant genes (particularly in *B. bruxellensis* and *B. nanus*) were identified. Numerous horizontal gene transfer (HGT) events in all *Brettanomyces* species’, including an HGT event that is probably responsible for allowing *B. bruxellensis* and *B. anomalus* to utilize sucrose were also observed.

**Conclusions:** Genomic adaptations and some evidence of domestication that have taken place in *Brettanomyces*are outlined. These new genome assemblies form a valuable resource for future research in *Brettanomyces*.

## Background

Most commercial alcoholic fermentations are currently performed by yeast from the genus *Saccharomyces* with the most common species being *Saccharomyces cerevisiae*. The domestication of *S. cerevisiae* is thought to have begun as early as prehistoric times [1].To date, many commercially available strains are available which have been highly selected for fermentation in harsh conditions, such as those encountered during wine, beer, and industrial bioethanol fermentations [2–4]. In parallel with *Saccharomyces*, a distantly related genus of budding yeasts, *Brettanomyces* (telemorph *Dekkera*), has also convergently evolved to occupy this same niche [5].

There are currently five widely accepted species of *Brettanomyces*: *B. anomalus*, *B. bruxellensis*, *B. custersianus*, *B. naardenensis*, and *B. nanus* [6]. A sixth species that was not included in this study, *Brettanomyces acidodurans*, was recently described and was tentatively assigned to this genus in G Péter, D Dlauchy, A Tóbiás, L Fülöp, M Podgoršek and N Čadež [7]. However, the authors note that it is genetically highly diverged from the other five species and could be considered a new genus. *Brettanomyces* are most commonly associated with spoilage in beer, wine, and soft drink due to the production of many off-flavour metabolites including acetic acid, and vinyl- and ethyl-phenols [5, 8, 9]. However, *Brettanomyces* can also represent an important and favorable component of traditional Belgian Lambic beers [10, 11], and their use has surged in recent years in the craft brewing industry [12]. Furthermore, *B. bruxellensis* has shown potential in bioethanol production by outcompeting *S. cerevisiae* and for its ability to utilize novel substrates [13, 14].

*B. bruxellensis* and to a lesser extent *B. anomalus*, are the main species encountered during wine and beer fermentation which has led to the majority of *Brettanomyces* research focusing only on these two species. The initial assembly of the triploid *B. Brettanomyces* strain AWRI1499 [15] has enabled genomics to facilitate research on this organism [16–20]. Subsequent efforts have seen the *B. bruxellensis* genome resolved to chromosome-level scaffolds [21]. In contrast, the assemblies that are available for *B. anomalus* [22], *B. custersianus*, and *B. naardenensis*, are less contiguous, and are mostly un-annotated, while no genome assembly is currently available for *B. nanus*.

Recent advancements in third-generation long-read sequencing have enabled the rapid production of highly accurate and contiguous genome assemblies, particularly for microorganisms (reviewed in S Koren and AM Phillippy [23]). This study sought to fill knowledge gaps for various *Brettanomyces* species by sequencing and assembling genomes using current-generation long-read sequencing technologies [24], and then to use these new assemblies to explore the genomic adaptations that have taken place across the *Brettanomyces* genus.

## Results and Discussion

### Significant improvements over current *Brettanomyces* genome assemblies

Information about the strains that were used in this study are shown in Table 1. In the interest of obtaining high-quality and contiguous assemblies, haploid or homozygous strains were favored (the *B. anomalus* strain was the exception), with strains that featured in past studies prioritized. All strains were isolated from commercial beverage products, with three from commercial fermentations. New genome assemblies for the five *Brettanomyces* species are described, generally exhibiting significant improvements over previous assemblies. Genome assembly summary statistics are shown in Table 2 and MinION sequencing statistics are available in Table S1. Genome sizes for the haploid *Brettanomyces* species ranged from 10.2 Mb (*B. nanus*) to 13.8 Mb (*B. anomalus*). The assemblies were similar in size to currently available assemblies [15, 21, 22], with overall assembly contiguity varying due to differences in heterozygosity and sequencing read lengths. The *B. anomalus* strain is a heterozygous diploid and while read coverage was high, the median read length was relatively low at 4.7 kb. This resulted in the lowest contiguity in the study consisting of 48 contigs with an N50 of 640 kb. The haploid *B. nanus* strain had a much higher median read length of 14.9 kb. As such, this assembly had the best contiguity consisting of only 5 contigs with an N50 of 3.3 Mb. To the best of our knowledge, this makes the *B. nanus* assembly the most contiguous *Brettanomyces* assembly to date. Furthermore, the *B. anomalus, B. custersianus,* and *B. naardenensis* assemblies represent 4.7-, 9.4-, and 6.5-fold improvements in contiguity over the currently available assemblies.

**Table 1:** Strain details and growth conditions

| ID | Species | Other IDs | Sample origin | Source; Reference |
| --- | --- | --- | --- | --- |
| AWRI950 | *B. custersianus* | CBS 4805 / IFO 1585 | Beer | CBS; [28] |
| AWRI951 | *B. naardenensis* | CBS 6042 / IFO 1588 | Soft drink | CBS; [29] |
| AWRI953 | *B. anomalus* | CBS 8139 | Soft drink | CBS; [30] |
| AWRI2804 | *B. bruxellensis* | UCD 2041 | Fruit wine | UC Davis Collection |
| AWRI2847 | *B. nanus* | CBS 1945 | Beer | CBS; [31] |

**Table 2:** Assembly and BUSCO summary statistics for the haploid assemblies

|  | *B. anomalus* | *B. anomalus*<br>(diploid) | *B. bruxellensis* | *B. custersianus* | *B. naardenensis* | *B. nanus* |
| --- | --- | --- | --- | --- | --- | --- |
| **Contigs** | 48 | 93 | 12 | 24 | 16 | 5 |
| **Length (Mb)** | 13.77 | 27.07 | 13.20 | 10.73 | 11.16 | 10.19 |
| **N50 (Mb)** | 0.640 | 0.730 | 2.936 | 0.847 | 1.231 | 3.303 |
| **GC (%)** | 39.81 | 39.84 | 39.88 | 40.24 | 44.60 | 41.51 |
| **BUSCOs (%)** |  |  |  |  |  |  |
| **Complete** | 83.0 | 84.2 | 88.6 | 88.2 | 90.6 | 90.6 |
| **-Single-copy** | 81.8 | 48.3 | 88 | 87.3 | 90 | 90.1 |
| **-Duplicate** | 1.2 | 35.9 | 0.6 | 0.9 | 0.6 | 0.5 |
| **Fragmented** | 9.8 | 8.8 | 6.1 | 6.4 | 5.6 | 5.2 |
| **Missing** | 7.2 | 7.0 | 5.3 | 5.4 | 3.8 | 4.2 |

The BUSCO results are shown in Table 2. Predicted genome completeness was high for the haploid assemblies, with between 3.8 % (*B. naardenensis*) and 7.2 % (*B. anomalus*) missing BUSCOs. The assemblies were processed with Purge Haplotigs [25] to remove duplicated and artifactual contigs. Duplication was low for not only the homozygous strains but also for the heterozygous *B. anomalus* assembly with between 0.5 % (*B. nanus*) and 1.2 % (*B. anomalus*) duplicate BUSCOs.

Given its heterozygous genome, a diploid assembly was also generated for the *B. anomalus* strain. The resultant diploid assembly was approximately twice the size of the haploid assembly and had a slightly higher N50 of 730 kb. While the genome size doubled, the duplicated BUSCOs only increased from 1.2 % for the haploid assembly to 35.9 % for the diploid assembly. This was mainly a result of BUSCOs having a fragmented gene model on only one of the two haplomes. It’s possible that this functional hemizygosity of core genes stabilizes the heterozygous nature of the genome. It should be noted that while the diploid *B. anomalus* assembly is split into Haplome 1 (H1) and Haplome 2 (H2), these haplomes consist of mosaics of both parental haplomes as haplotype switching can randomly occur between pairs of separated phase blocks.

The *B. nanus* strain that was used in this study exhibited a more reduced genome (loss of genes and reduction of intergenic sequence) when compared to the other species. Loss of genes can occur when a new environment (such as a nutrient-rich medium) results in genes that were previously indispensable are no longer required for survival, or gene loss can even confer an adaptive advantage (reviewed in R Albalat and C Cañestro [26]). Genome compaction can occur via shortening of intergenic regions and removal of pseudogenes [27]; this is more common in bacteria. The number of predicted genes and the gene densities (as percent of genome that is genic) for the *Brettanomyces* genomes as well as the reference genome for *S. cerevisiae* strain S288C are shown in Table S2. *B. nanus* had the highest compaction with both the fewest genes (5,083) and highest gene density (78.1 %). *B. naardenensis* and *B. custersianus* both had higher gene densities (75.2 % and 75.4 % respectively) than *B. bruxellensis*, *B. anomalus*, and *S. cerevisiae* S288C (64.2 %, 62.2 %, and 74.1 % respectively).

### Revisiting the taxonomy of *Brettanomyces*

Availability these new *Brettanomyces* genomes allowed for a comprehensive phylogeny to be generated utilizing the entire genome as opposed to extrapolating from ribosomal segments. Codon-based alignments were produced for 3482 single-copy orthologues (SCOs) that were found across the five *Brettanomyces* species, in addition to *Ogataea polymorpha* (*Brettanomyces*’ closest relative) as an outgroup. This data was used to calculate a maximum-likelihood tree (Figure 1a) and to estimate average nucleotide identity (ANI) between pairs of genomes (Table 3). This method was also applied to the *Saccharomyces sensu stricto* clade (with *Naumovozyma castellii*, *Saccharomyces*’ closest relative, as the outgroup) to serve as a comparison (Figure 1b and Table 4). The *Brettanomyces* whole-genome phylogeny was first compared to trees of *Brettanomyces* produced in previous studies. The whole-genome phylogeny generally agreed with the trees derived from rRNA sequences as described in Y Yamada, M Matsuda, K Maeda and K Mikata [32], Y Yamada, M Matsuda and K Mikata [33], and C Röder, H König and J Fröhlich [34]. However, these earlier studies were not able to consistently resolve the placement of *B. nanus*, with conflicting results between phylogenies based on 18S and 26S ribosomal RNA sequences. By utilizing the entire genome, it is now possible to confirm that *Brettanomyces* forms two clades, with *B. nanus* and *B. naardenensis* forming a clade separate from the other species (consistent with the 18S trees in these earlier studies).

**Figure 1:**
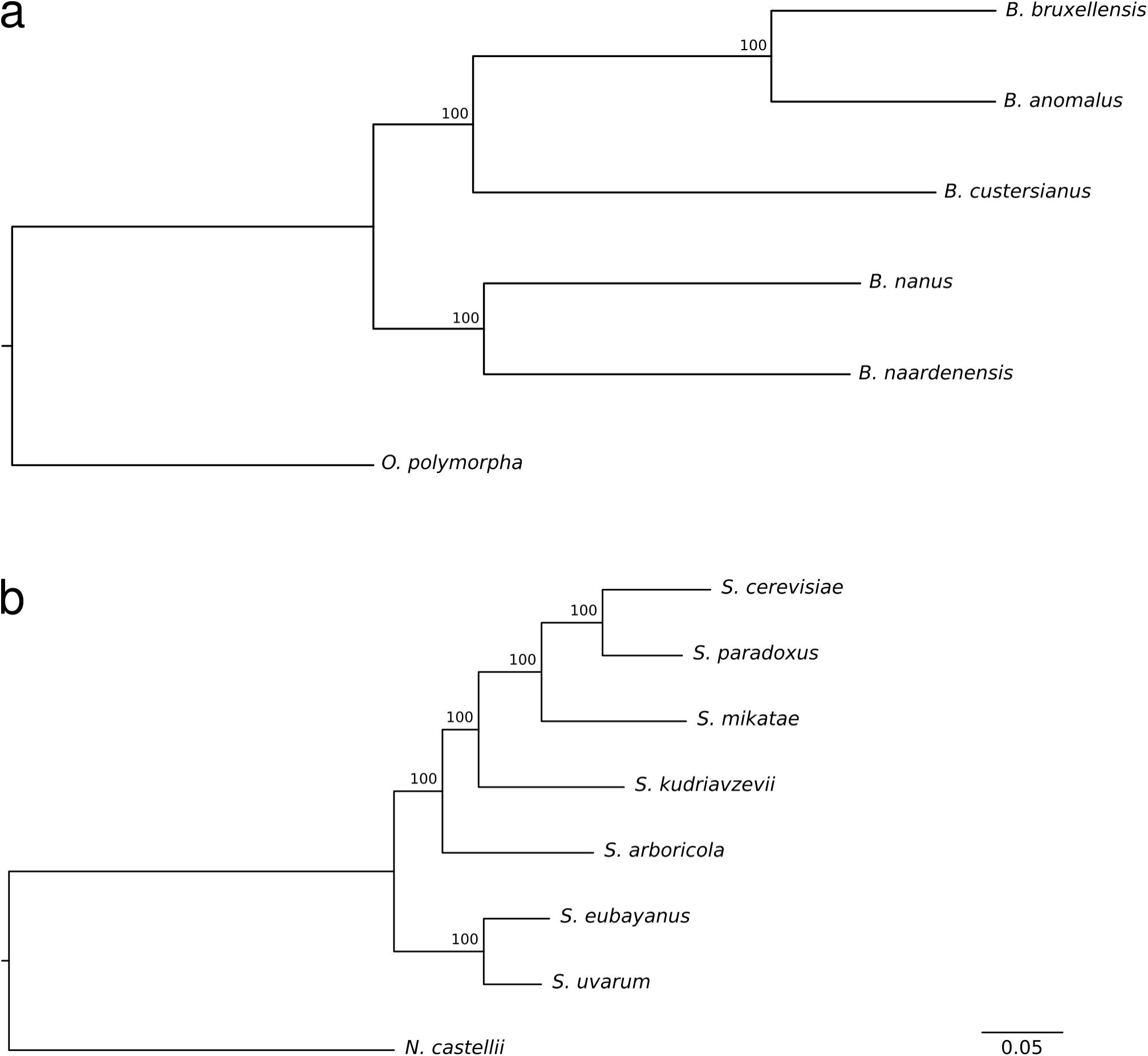
Phylogenies of *Brettanomyces* and *Saccharomyces* species. Rooted, maximum likelihood trees were calculated for *Brettanomyces* species with *Ogataea polymorpha* as an outgroup (*a*) and *Saccharomyces* species with *Naumovozyma castellii* as an outgroup (*b*). The phylogenies were calculated from concatenated codon alignments of single copy orthologs. Bootstrap values are calculated from 100 replications are shown at branch nodes. The two phylogenies are transformed to the same scale (substitutions per site), indicated bottom right.

**Table 3:** Average Nucleotide Identities (percent) between *Brettanomyces* species and *Ogataea polymorpha* concatenated single copy ortholog codon alignments.

|  | *B. naardenensis* | *B. bruxellensis* | *B. custersianus* | *O. polymorpha* | *B. anomalus* |
| --- | --- | --- | --- | --- | --- |
| *B. nanus* | 66.4 | 60.6 | 61.0 | 56.3 | 60.7 |
| *B. naardenensis* |  | 60.8 | 61.3 | 56.6 | 60.9 |
| *B. bruxellensis* |  |  | 60.7 | 55.1 | 77.1 |
| *B. custersianus* |  |  |  | 54.8 | 60.8 |
| *O. polymorpha* |  |  |  |  | 55.2 |

**Table 4:** Average Nucleotide Identities (percent) between *Saccharomyces* species and *Naumovozyma castellii* concatenated single copy ortholog codon alignments.

|  | *S. eubayanus* | *S. uvarum* | *S. cerevisiae* | *N. castellii* | *S. paradoxus* | *S. mikatae* | *S. kudriavzevii* |
| --- | --- | --- | --- | --- | --- | --- | --- |
| *S. arboricola* | 82.1 | 82.4 | 81.1 | 61.6 | 81.9 | 81.3 | 83.2 |
| *S. eubayanus* |  | 92.8 | 79.9 | 61.6 | 80.6 | 80.1 | 81.8 |
| *S. uvarum* |  |  | 80.1 | 61.5 | 80.9 | 80.3 | 82.2 |
| *S. cerevisiae* |  |  |  | 61.6 | 89.3 | 84.0 | 81.9 |
| *N. castellii* |  |  |  |  | 61.6 | 61.6 | 61.4 |
| *S. paradoxus* |  |  |  |  |  | 85.2 | 82.8 |
| *S. mikatae* |  |  |  |  |  |  | 82.2 |

There is generally a very large genetic distance separating the *Brettanomyces* species, and this is particularly striking when comparing the phylogenies for *Brettanomyces* and *Saccharomyces*. There is a greater genetic distance between most of the *Brettanomyces* species than there is between any of the *Saccharomyces* species and the *N. castellii* outgroup. The largest separation is between *B. nanus* and *B. bruxellensis* with an ANI of only 60.6 %. The closest relation between *Brettanomyces* species is between *B. bruxellensis* and *B. anomalus* with an ANI of 77.1 %, followed by *B. nanus* and *B. naardenensis* with an ANI of 66.4 %. The remainder are between 60.6 % and 61.3 % ANI. As a comparison, the ANIs between the *Saccharomyces* species and the outgroup (*N. castellii*) ranged from 61.4 % (*Saccharomyces kudriavzeviiI*) to 61.6 % (*Saccharomyces cerevisiae*). Furthermore, the genetic distance between the most distantly related *Saccharomyces* species (*S. cerevisiae* and *Saccharomyces eubayanus*, ANI of 79.9 %) is less than the genetic distance between the most closely related *Brettanomyces* species. Given the distinct clades for *Brettanomyces*, together with the relatively very large genetic distances separating them, it may be appropriate for the *Brettanomyces* genus to be divided and a new genus proposed for *B. nanus* and *B. naardenesis*.

### Extensive rearrangements are present throughout *Brettanomyces* genomes

Genomic rearrangements featured extensively between all *Brettanomyces* genomes in this study (Figure 2). There were numerous small and several large translocations visible between the *B. bruxellensis* and the *B. anomalus* assemblies (Figure 2a), and to a lesser extent the *B. bruxellensis* and *B. custersianus* assemblies (Figure S1). Comparing *B. bruxellensis* to the more distantly related species *B. naardenensis* (Figure 2b) and *B. nanus* (Figure 2c), these large and small breaks in synteny appear even more extensively. The chromosomal rearrangements were not limited to a single species or clade; when comparing *B. nanus* to *B. naardenensis* (Figure 2d) there is a similar level of rearrangements to that occurring between *B. bruxellensis* and *B. anomalus*.

**Figure 2:**
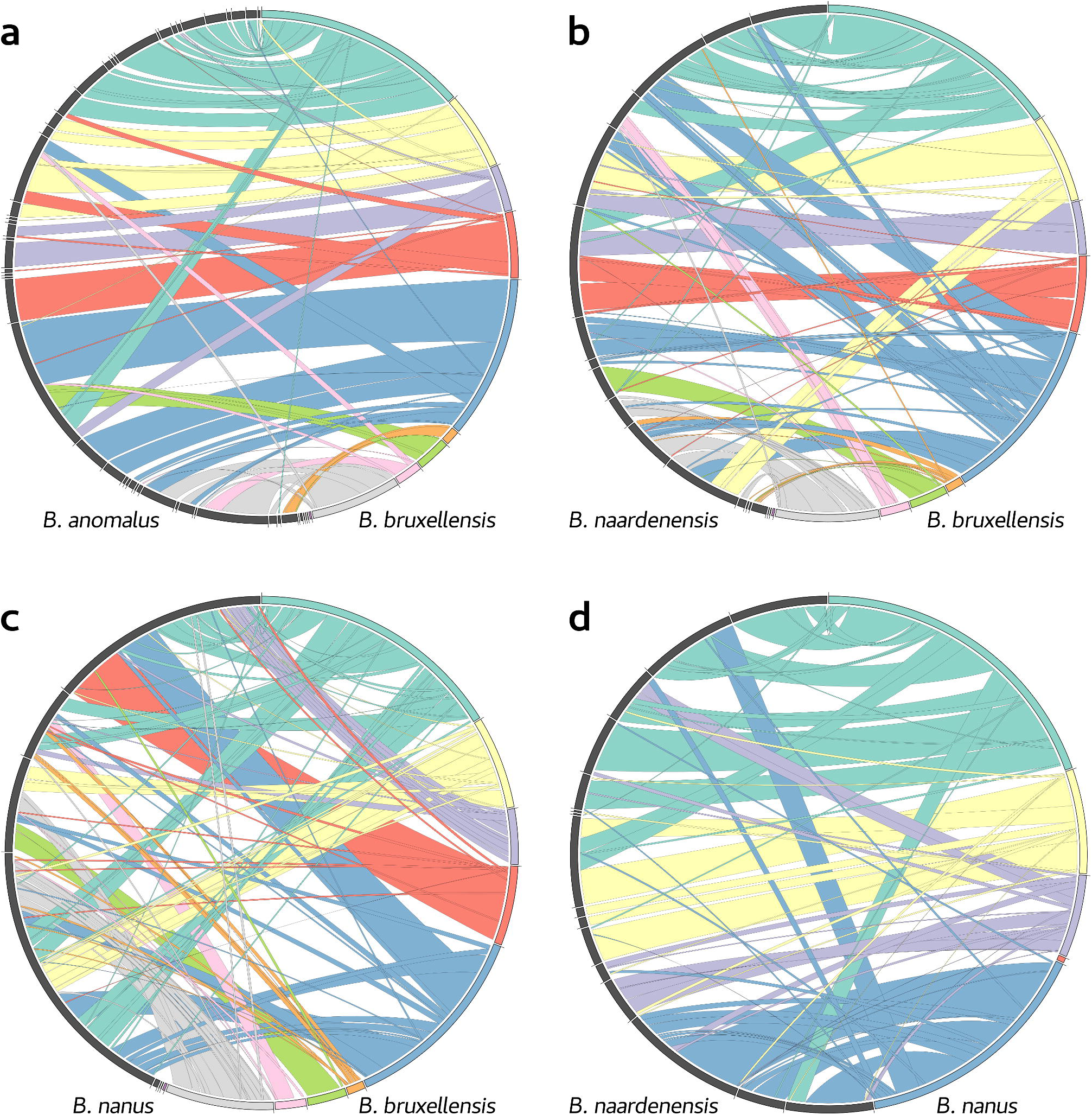
Synteny between haploid assemblies of *Brettanomyces*, visualized as Circos plots. Reference assembly Contigs are coloured sequentially. Alignments are coloured according to the reference assembly contigs and are layered by alignment length. The query assembly contigs are coloured grey. Alignments are depicted between *B. bruxellensis* and *B. anomalus* (*a*), *B. bruxellensis* and *B. naardenensis* (*b*), *B. bruxellensis* and *B. nanus* (*c*), and *B. nanus* and *B. naardenensis* (*d*).

Chromosomal rearrangements, and karyotype and ploidy variability have been reported previously in *Brettanomyces* [17, 35–39]. This genome plasticity is thought to be a mechanism in yeast for adaptation to new environments and niches, and in response to new stressors (see S Marsit, J-B Leducq, É Durand, A Marchant, M Filteau and CR Landry [40] for a review). Another one of these mechanisms—loss-of-heterozygosity (LOH)—is present in the heterozygous *B. anomalus* genome. Three large contigs, comprising 2.14 Mb (15 %) of the *B. anomalus* genome, were predicted to be homozygous (0.0353 SNPs/kb) while the rest of the genome is heterozygous (3.21 SNPs/kb) (Figure S2). The strains used in this study as reference for *B. bruxellensis, B. custersianus*, *B. naardenensis*, and *B. nanus* appeared homozygous as expected, with heterozygous SNP densities ranging from 0.01 (*B. naardenensis)* to 0.05 (*B. bruxellensis)* SNPs/kb.

### *Brettanomyces* species harbor enrichments of fermentation-relevant genes

Species-specific expansion of specific gene families was investigated across the *Brettanomyces* genomes with enriched gene ontologies identified for each of the species (Table 4). Both *B. bruxellensis* and *B. nanus* are predicted to have undergone copy number expansion of ORFs predicted to encode oligo-1,6-glucosidase (EC 3.2.1.10) enzymes (Figure 3a), which are commonly associated with starch and galactose metabolism. *B. nanus* is also predicted to possess an expanded set of genes encoding β-glucosidase (EC 3.2.1.21) (Figure 3b) and β-galactosidase (EC 3.2.1.23) activities (Figure 3c). These specializations for scavenging sugars from complex polysaccharides are a hallmark of the domestication of beer and wine strains of *S. cerevisiae* and suggest that the same may be occurring in *B. nanus* [41–43]. The three known *B. nanus* strains that have been isolated to date were all sourced from beer samples obtained from Swedish breweries in 1952. The *B. nanus* strain AWRI2847 (CBS 1945) was evaluated in V Harris, CM Ford, V Jiranek and PR Grbin [44] and was found to have far less spoilage potential than either *B. bruxellensis* or *B. anomalus*. At the time of this strain’s original isolation, spoilage was determined sensorially and sharing yeast samples between breweries was common practice [45]. Yeast from a completed beer fermentation is commonly used to inoculate (re-pitch) the next batch of wort. Taken together, it may be possible that *B. nanus* represented a long-term undetected contaminant, surviving successive serial re-pitchings and spreading to multiple breweries, thus allowing these genomic adaptations to manifest.

**Figure 3:**
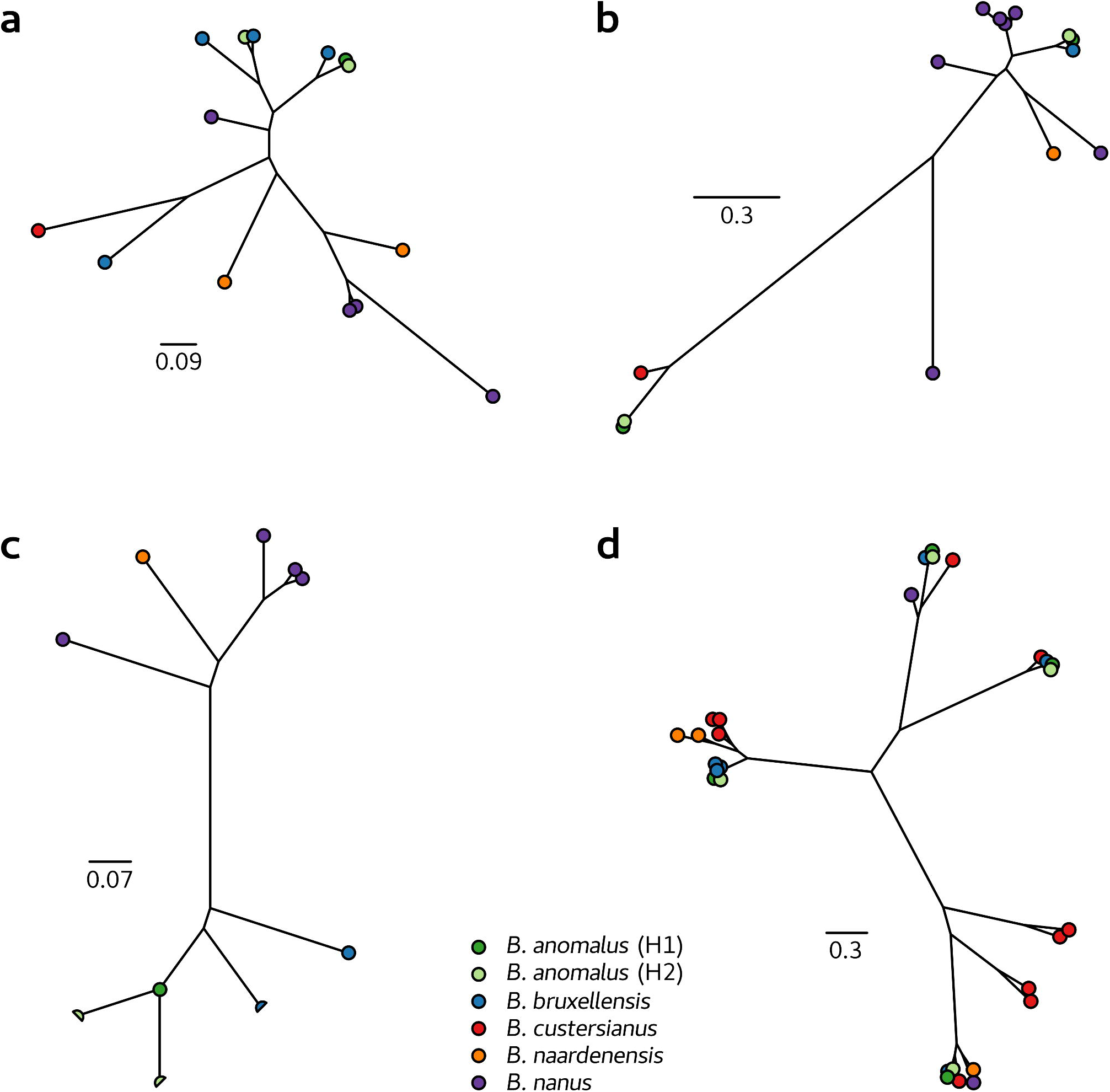
Phylogenies of several enriched orthogroups in *Brettanomyces*. Broken gene models or pseudo-genes are indicated as half circles. The enriched gene orthogroups are: oligo-1,6-glucosidase (EC 3.2.1.10) (*a*), β-glucosidase (EC 3.2.1.21) (*b*), β-galactosidase (EC 3.2.1.23) (*c*), and sarcosine oxidase (EC 1.5.3.1/1.5.3.7) (*d*).

Both *B. custersianus* and *B. bruxellensis* presented large expansions (10 and 6 copies respectively) of genes encoding sarcosine oxidase / L-pipecolate oxidase (PIPOX) (EC 1.5.3.1/1.5.3.7) and the remaining *Brettanomyces* species also contain multiple copies of this gene. The evolutionary expansion of this gene family is complex, but it appears as though multiple independent duplications occurred (Figure 3d). PIPOX exhibits broad substrate specificity but primarily catalyses the breakdown of sarcosine to glycine and formaldehyde, as well as the oxidation of L-pipecolate [46]. However, it has been shown to also act on numerous other *N*-methyl amino acids such as *N*-methyl-L-alanine, *N*-ethylglycine, and both L- and D-proline [46–49]. *Brettanomyces* are adapted to grow in nutrient-depleted conditions and this has largely been attributed to the utilization of alternative nitrogen sources such as free nitrates and amino acids [50–52]. Interestingly, proline—a substrate of PIPOX—is one of the more common amino acids in fermented wine and beer. Proline is poorly utilized by *S. cerevisiae* (and is actually produced during fermentation as a means of maintaining redox homeostasis), but is readily metabolized in *B. bruxellensis* [53–56]. PIPOX converts proline to 1-pyrroline-2-carboxylate, which can ultimately be converted to D-Ornithine by the action of a general aminotransferase. As opposed to the redox cofactor-dependent enzymes proline oxidase (EC 1.5.1.2) and proline dehydrogenase (EC 1.5.5.2), PIPOX could therefore represent an avenue for proline utilisation that does not directly impact redox homeostasis.

Beyond PIPOX, *B. bruxellensis* and *B. anomalus* share an expansion of S-formylglutathione hydrolase (EC 3.1.2.12), and *B. anomalus* contains an expansion of formate dehydrogenase (EC 1.17.1.9). While these genes are part of methanol metabolism in other species, a capability that is lost in *Brettanomyces*, both genes are involved with the metabolism of formaldehyde (a common metabolic byproduct during fermentation). Lastly, *B. naardenensis* contains an expansion of a gene encoding sulfonate dioxygenase (EC 1.14.11.-) activity, which is associated with the utilisation of alternative sulphur sources, in addition to an expansion of acetylornithine deacetylase (EC 3.5.1.16), a component of the arginine biosynthetic pathway.

In order to identify genes that are important in the evolution of the *Brettanomyces* genus, the *Brettanomyces* genomes were examined to identify single copy orthologs with nonsynonymous mutations under strong selective pressure (residues under site-selection). There were 279 SCOs identified, 182 of which had KEGG annotations (Table S3). These include many ribosomal proteins and transcription factors that are commonly under site-based selection, as well as numerous notable metabolic enzymes. Four enzymes identified are known to be involved in balancing the metabolic needs of the cell during cell proliferation: Pyruvate kinase (EC 2.7.1.40), Ribose-phosphate pyrophosphokinase (EC 2.7.6.1), Pyruvate carboxylase (EC 6.4.1.1), and Thymidylate synthase (EC 2.1.1.45). Further enzymes are involved with the biosynthesis of B-vitamin cofactors related to carbohydrate metabolism: Biotin synthase (EC 2.8.1.6), Pantothenate kinase (EC 2.7.1.33), Phosphopantothenoylcysteine decarboxylase (EC 4.1.1.36), and Thiamine pyrophosphokinase (EC 2.7.6.2). Finally, seven genes in the MAPK signaling pathway were identified, relating to osmotic stress response and pheromone response.

**Table 4a:**
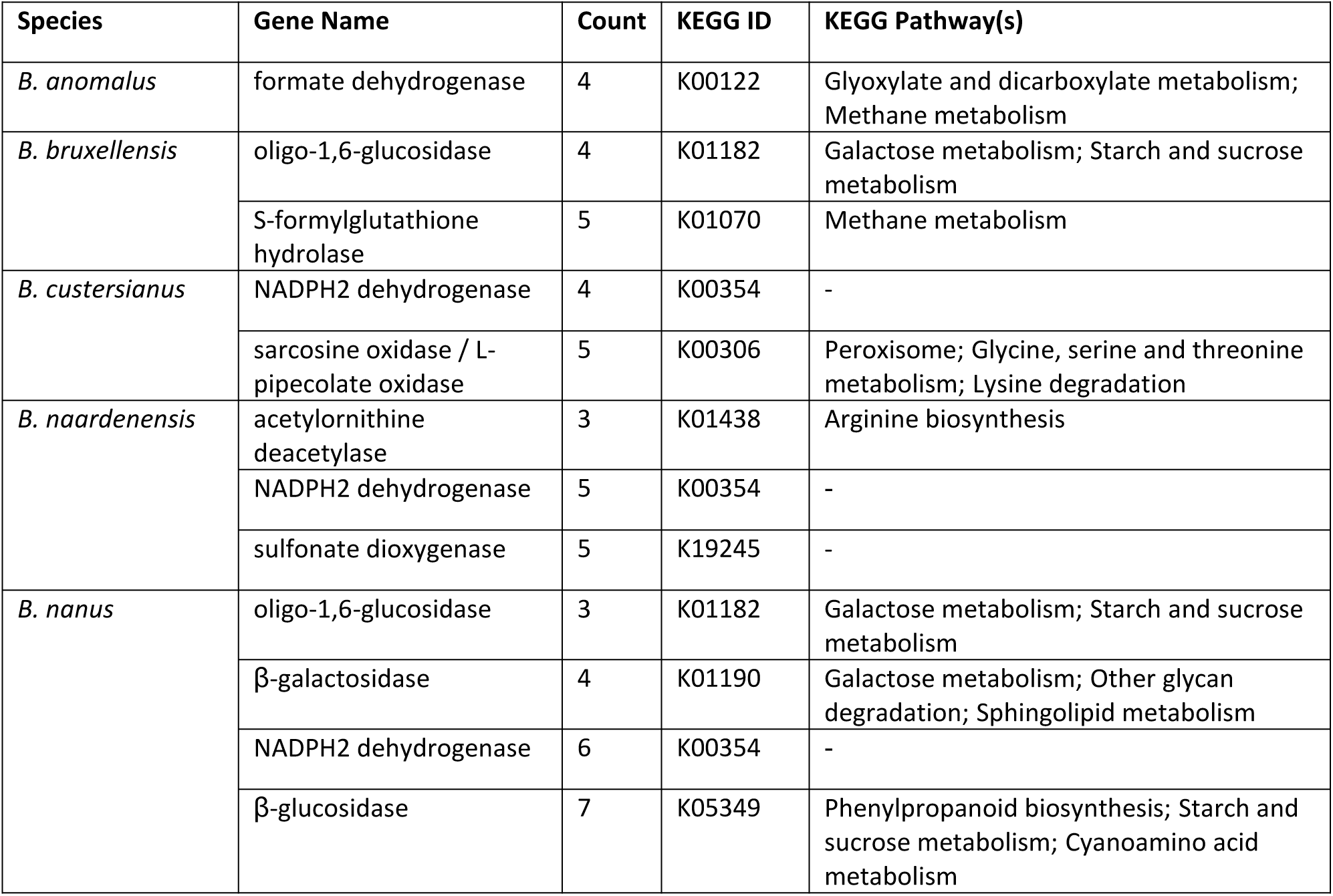
Expanded gene families in *Brettanomyces*

### Horizontal gene transfer enables sucrose utilization in *B. bruxellensis* and *B. anomalus*

Horizontal Gene Transfer (HGT) has been reported as a mechanism of adaptative evolution in fungal species and to have contributed to the domestication of *S. cerevisiae* [57–59]. Consequently, potential HGT events that may have contributed to the evolution of *Brettanomyces* were investigated. Twelve *Brettanomyces* orthogroups could be identified with genes that were predicted to be the result of HGT from bacteria according to protein homology with RefSeqKB database protein sequences (Table 5). Of these bacterially derived gene families, β-fructofuranosidase (also known as Invertase) (EC 3.2.1.26), stands out as having a key phenotypic impact. Invertases are responsible for the conversion of sucrose to fructose and glucose and this enzyme activity is required for the utilization of sucrose as a carbon source.

**Table 5:** Genes predicted to occur in *Brettanomyces* via Horizontal Gene Transfer

| Orthogroup<br>(Saccharomycetaceae) | Gene name | KEGG | Species (gene IDs) | Closest genus<br>BLAST hit(s) |
| --- | --- | --- | --- | --- |
| OG0000021 | Glucose 1-dehydrogenase | - | *B. naardenensis* (g20, g1494, g1495, g1768, g1785, g1792, g4886, g5231)<br>*B. nanus* (g5077) | *Labilithrix*,<br>*Methylovulum* |
| OG0000714 | glutamine amidotransferase | - | *B. nanus* (g3549),<br>*B. naardenensis* (g138) | *Cyanobacterium* |
| OG0001026 | $\alpha/\beta$ hydrolase | - | *B. naardenensis* (g5185) | *Klebsiella* |
| OG0003977 | nitronate monooxygenase | - | *B. naardenensis* (g5218) | *Bordetella* |
| OG0003988 | β-fructofuranosidase (invertase) | K01193 | *B. anomalus* H1 (g3595),<br>*B. anomalus* H2 (g273),<br>*B. bruxellensis* (g1543) | *Bacillus*,<br>*Saccharothrix* |
| OG0005068 | NADP oxidoreductase | - | *B. naardenensis* (g4928, g5192, g5211) | *Halomonas* |
| OG0005439 | flavodoxin family protein | K08071 | *B. naardenensis* (g1784) | *Gluconobacter* |
| OG0005699 | cysteine hydrolase | - | *B. naardenensis* (g5191) | *Pseudomonas* |
| OG0005912 | capsule biosynthesis protein<br>CapA | - | *B. anomalus* H1 (g1924),<br>*B. anomalus* H2 (g5929),<br>*B. bruxellensis* (g4262),<br>*B. custersianus* (g2790),<br>*B. nanus* (g654) | *Izhakiella* |
| OG0006075 | YbjQ family protein | - | *B. bruxellensis* (g4270),<br>*B. naardenensis* (g4529, g4528, g1637),<br>*B. nanus* (g1955) | *Streptomyces* |
| OG0006081 | S-antigen protein | - | *B. naardenensis* (g3365) | *Symbiodinium* |
| OG0006556 | NAD(P)-dependent<br>oxidoreductase | - | *B. anomalus* H1 (g109),<br>*B. anomalus* H2 (g1567),<br>*B. bruxellensis* (g2456) | *Pleomorphomonas* |

To confirm the bacterial origins of the *Brettanomyces* invertases, a protein-based phylogeny was created from the known fungal and bacterial invertases in the RefSeqKB database, as well as from the three *Brettanomyces* invertases (Figure 4a). The fungal invertases form one distinct clade, while the bacterial proteins are spread across several distinct groups. Consistent with a bacterial-derived HGT event, the *Brettanomyces* invertase proteins reside within a bacterial clade and are evolutionarily distinct from the fungal group. The invertases present in *B. bruxellensis* and *B. anomalus* appear to reside within subtelomeres (Figure 4b); these are genomic regions that are known to be hotspots for structural rearrangements and HGT events in *Saccharomyces* [60–64]. In *Brettanomyces*, there is significant structural variation and a general loss of synteny that is typical of subtelomeric regions in other species (Figure 4b). For example, in *B. nanus* the NAG gene cluster resides within a different subtelomere relative to *B. bruxellensis* and *B. anomalus*. The NAG genes are also present in *B. naardenensis* but they are not co-located, and they appear to be missing entirely in *B. custersianus*. Likewise, homologues of *MPH3* and *TIP1* genes are present in all *Brettanomyces* species, but are only found in this specific subtelomeric region in *B. bruxellensis* and *B. anomalus*.

**Figure 4:**
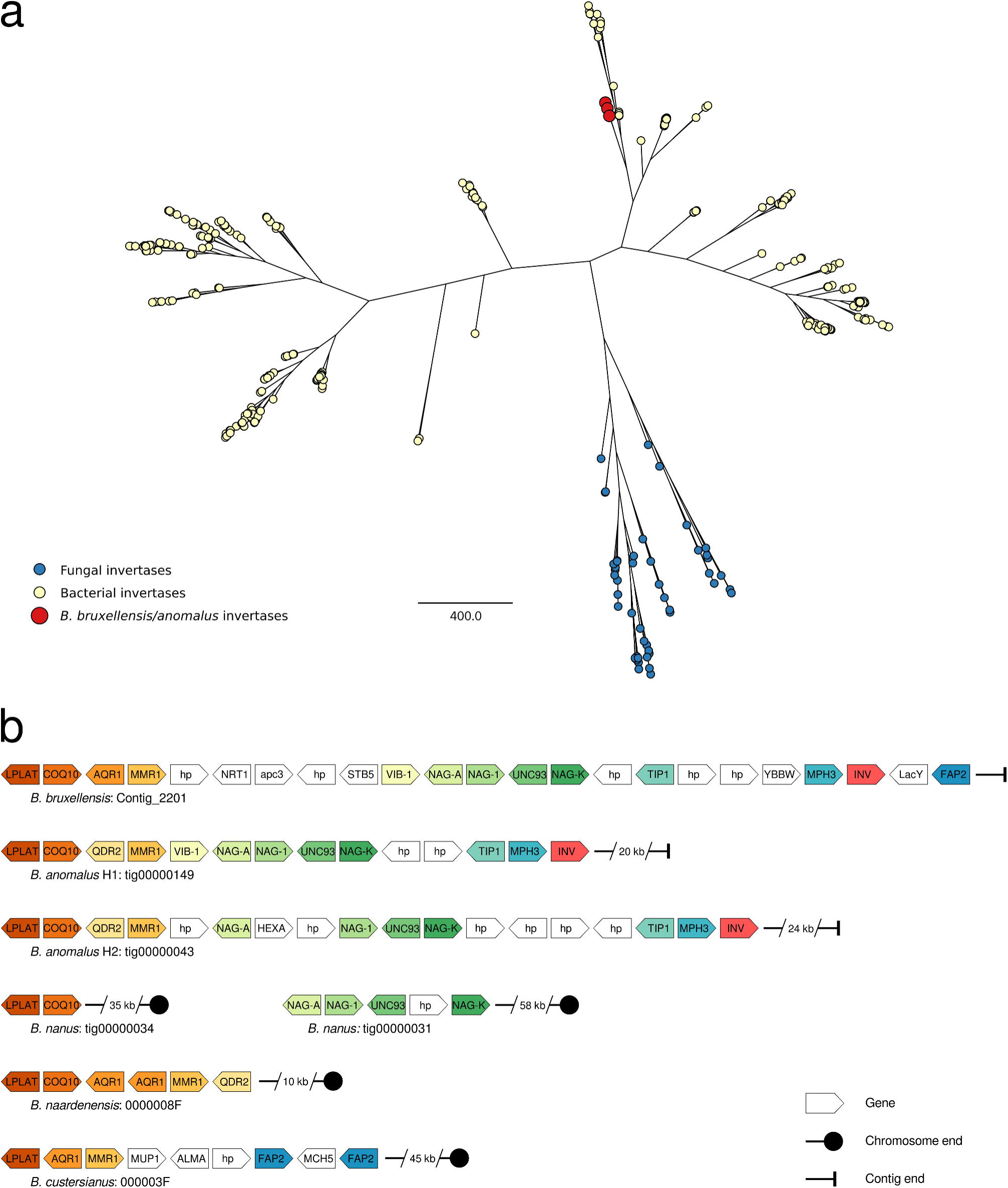
β-fructofuranosidases (invertases) from *Brettanomyces*. Phylogeny of invertases from *Brettanomyces* and the RefSeqKB fungal and bacterial databases, with *Brettanomyces* nodes enlarged for clarity (*a*). Genomic context of invertases in *Brettanomyces*, showing cluster of conserved genes, orange; NAG gene cluster, green; cluster of metabolic genes, blue; Invertase, red (*b*).

Previous phenotypic testing has shown *B. bruxellensis* and *B. anomalus* to be the only *Brettanomyces* species capable of utilizing sucrose [6] and this phenotype correlates with the presence of the HGT-derived invertases, which are only observed in the *B. bruxellensis* and *B. anomalus* genomes (there are no other invertase encoding ORFs predicted in *Brettanomyces*). Sucrose utilization likely conferred a significant advantage in fruit fermentations, possibly shaping the evolution of the common ancestor of *B. bruxellensis* and *B. anomalus* towards fermentation specialization.

## Conclusions

High quality genome assemblies for all five currently accepted *Brettanomyces* species are described, including the first assembly for *B. nanus* and the most contiguous assemblies available to date for *B. anomalus*, *B. custersianus*, and *B. naardenensis*. Comparative genome analysis established that the species are genetically distant and polyphyletic. Numerous indicators of domestication and adaptation in *Brettanomyces* were identified with some notable parallels to the evolution of *Saccharomyces*. Extensive structural differences between the genomes of the *Brettanomyces* species and apparent loss of heterozygosity in *B. anomalus* were observed. Enrichments of fermentation-relevant genes were identified in *B. anomalus*, *B. bruxellensis* and *B. nanus*, as well as multiple horizontal gene transfer events in all *Brettanomyces* genomes, including a gene in the *B. anomalus* and *B. bruxellensis* genomes that is probably responsible for these species’ ability to utilize sucrose.

## Methods

Detailed workflows, custom scripts for computational analyses and annotations are available in Additional File 1. All sequencing reads and genome assemblies have been deposited at the National Center for Biotechnology Information (NCBI) Sequence Read Archive (SRA) under the BioProject: PRJNA554210. Raw FAST5-format files for all Oxford Nanopore sequencing are available from the European Bioinformatics Institute (EMBL-EBI) European Nucleotide Archive (ENA) under the study: ERP116386.

### Strains and media

The five *Brettanomyces* strains selected for sequencing were supplied by the Australian Wine Research Institute’s wine microorganism culture collection. Strains were grown in either MYPG medium (0.3% malt extract, 0.3% yeast extract, 0.2% peptone, 1% glucose) at 27°C or in GPYA+CaCO3 medium (4% glucose, 0.5% peptone, 0.5% yeast extract, 1% calcium carbonate) at 25°C.

### Library preparation and sequencing

Genomic DNA was extracted from liquid cultures using a QIAGEN Gentra Puregene Yeast/Bact Kit. *B. bruxellensis* was sequenced using PacBio RS-II SMRT sequencing. The sequencing library for *B. nanus* was multiplexed with other samples (not reported here) using the SQK-LSK109 and EXP-NBD103 kits following the Oxford Nanopore protocol NBE_9065_v109_revA 23MAY2018. For the remaining species, libraries were prepared using the SQK-LSK108 kit following the protocol

GDE_9002_v108_revT_18OCT2016. Sequencing was performed on a MinION using FLO-MIN106 flow-cells. Demultiplexing and base-calling were performed using Albacore v2.3.1.

Illumina sequencing was performed on each strain using a combination of short-insert (TruSeq PCR-free) and mate-pair (2-5kb insert and 6-10 kb insert) libraries. All libraries were barcoded and pooled in a single Miseq sequencing run using 2x300bp chemistry.

### Assembly

The *B. bruxellensis* genome in this study was assembled with Mira v4.9.3 [65] using a hybrid assembly approach consisting of illumina paired-end and mate-pair reads, and PacBio long-reads. This assembly was manually finished in DNASTAR SeqMan Pro. Haploid assemblies for all other *Brettanomyces* species were generated from FASTQ-format Nanopore reads using Canu v1.7 [66]. The Nanopore reads were mapped to the assemblies using minimap2 [67] and initial base-call polishing was performed with Nanopolish v0.9.2 [68], utilizing the FAST5 signal-level sequencing data. Further base-call polishing was performed with Illumina paired-end, and 2–4 kb and 6–10kb mate-pair reads. Paired-end and mate-pair reads were mapped with BWA-MEM v0.7.12-r1039 [69] and Bowtie2 v2.2.9 [70] respectively; base-call polishing was then performed with Pilon v1.22 [71]. Finally, raw Nanopore reads were mapped to the base-call-polished assemblies and Purge Haplotigs v1.0.1 [25] was used to remove any duplicate or artefactual contigs.

A diploid assembly for AWRI953 (*B. anomalus*) was also generated. Paired-end reads were mapped to the haploid assembly with BWA-MEM, and high-confidence SNPs were called using VarScan v2.3.9 [72]. Nanopore reads were mapped to the assembly using BWA-MEM. Heterozygous SNPs were phased using the mapped Nanopore reads with HapCut2 commit: c2e6608 [73] and converted to VCF format with WhatsHap v0.16 [74]. New consensus sequences were called for each haplotype from the phased SNPs and the nanopore reads were binned according to which haplotype they mapped best. The two *B. anomalus* haplotypes were then independently reassembled from the haplotype-binned nanopore reads using the method described for the other species.

All other *Brettanomyces* assemblies were aligned to the *B. bruxellensis* assembly using NUCmer (MUMmer) v4.0.0beta2 [75]. Dotplots were visualized and contigs with split alignments were manually inspected for indications of mis-assemblies using mapped alignments of Nanopore reads and Illumina mate-pair reads. Genome metrics were calculated with Quast [76] and completeness, duplication, and fragmentation were estimated using BUSCO v3.0.2 [77] with the odb9 Saccharomyceta dataset.

### Annotation

Gene models were predicted with Augustus v3.2.3 [78] using the *S. cerevisiae* S288C configuration. Gene models were submitted for KEGG annotation using BlastKOALA [79], and GO-terms were annotated using InterProScan v5.32-71.0 [80]. Orthogroups were assigned with OrthoFinder v2.2.6 [81] using representative species from Saccharomycetaceae (Table S5) and also using only the haploid *Brettanomyces* assemblies.

### Phylogeny

Orthofinder (*Brettanomyces* + *O. polymorpha*) was used to find SCOs over these genomes. Protein sequences were aligned with Muscle v3.8.31 [82] and then converted to codon-spaced alignments using PAL2NAL [83]. Average nucleotide identities were estimated using panito commit: f65ba29 (github.com/sanger-pathogens/panito). A rooted maximum likelihood phylogeny was generated in R using ape [84] and phangorn [85]. A phylogeny was created using the same method for the *Saccharomyces sensu stricto* species + *N. castellii* (outgroup) to serve as a comparison.

### Whole genome synteny visualization

Pairwise synteny blocks were generated between the reference *B. bruxellensis* assembly and the other haploid assemblies, as well as between the *B. naardenensis* and *B. nanus* assemblies. Contigs were placed in chromosome order using Purge Haplotigs [25] to generate placement files that were then used to rearrange contigs. Alignments between the assemblies were calculated using NUCmer with sensitive parameters (-b 500 -c 40 -d 0.5 -g 200 -l 12). Genome windows (20 kb windows, 10 kb steps) were generated for the assemblies and a custom script was used to pair syntenic genome windows based on the NUCmer alignments. Concordant overlapping and adjacent windows were merged, and overlapping discordant windows were trimmed. The synteny blocks were then visualized using Circos v0.69.6 [86].

### Gene enrichment and selection

OrthoFinder (Saccharomycetaceae) annotations were used to identify gene-count differences between the *Brettanomyces* species. The ratio of the gene-count to the average gene-count was calculated for the *Brettanomyces* species over all OrthoFinder orthogroups. All orthogroups with a ratio ≥ 2 for any *Brettanomyces* species were subject to GO-enrichment analysis using BiNGO v3.0.3 [87] using the hypergeometric test with Bonferroni FWER correction. Genes for overrepresented categories (p-value ≤ 0.05) were returned. Multiple sequence alignments were generated for GO-enriched orthogroups using Muscle and phylogeny trees generated using PhyML within SeaView v4.7 [88] using default parameters.

Adaptive selection was predicted for SCOs on an OrthoFinder run consisting only of the haploid *Brettanomyces* assemblies. Protein sequences were initially aligned with Muscle and then converted to codon-spaced alignments using PAL2NAL. Protein alignments were concatenated and used to produce an unrooted species tree using PhyML within SeaView. Transcript alignments were assessed with codeml from PAML v4.9 [89] against the site models M1a (nearly neutral selection), M2a (adaptive selection), M7 (no adaptive selection), and M8 (adaptive selection). Log ratio tests of maximum likelihoods were used to filter for M2a vs. M1a and M8 vs. M7 models and orthogroups with sites under selection (according to either Naïve Empirical Bayes analysis or Bayes Empirical Bayes analysis, P > 95 %) were returned.

### Horizontal gene transfer

HGT events were predicted for the *Brettanomyces* species. Protein sequences for the assemblies were used in BLAST-P searches against the RefSeqKB non-redundant Fungi and Bacteria datasets [90]. All *Brettanomyces* proteins with a higher scoring hit to a Bacterial protein than a Fungal protein were investigated further. The multiple sequence alignments and trees were retrieved for the HGT candidates’ orthogroups and several candidates were removed following manual inspection. A phylogeny was generated for one HGT prediction of interest. The *Brettanomyces* genes, and the orthologs from the ResSeq Fungal and bacterial datasets were aligned with Muscle, and the phylogeny was generated in R using Ape and Phangorn.

## List of Abbreviations

BUSCO: The name of the pipeline for detecting BUSCOs
BUSCO(s): Benchmarking universal single-copy ortholog(s)
H1/H2: Haplome 1/Haplome 2 (*B. anomalus*)
HGT: Horizontal gene transfer
LOH: Loss of heterozygosity
SCO: Single copy ortholog

## Declarations

### Ethics approval and consent to participate

Not applicable.

### Consent for publication

Not applicable.

### Availability of data and materials

All sequencing reads and genome assemblies have been deposited at the National Center for Biotechnology Information (NCBI) Sequence Read Archive (SRA) under the BioProject: PRJNA554210. Raw FAST5-format files for all Oxford Nanopore sequencing are available from the European Bioinformatics Institute (EMBL-EBI) European Nucleotide Archive (ENA) under the study: ERP116386.

### Authors’ contributions

ARB conceived and designed the work and assisted with data collection, analysis and drafting the manuscript. MJR designed and performed in silico analysis and drafted the manuscript. All authors read and approved the final manuscript.

## Acknowledgements

The authors thank Cristobal Onetto (Australian Wine Research Institute) for helping with data collection, and Simon Schmidt and Markus Herderich (Australian Wine Research Institute) for critically reviewing the manuscript.

## Supplementary Tables and Figures

### Additional File 1: Archive.zip

**Workflows, scripts, and annotations for the *Brettanomyces* genome assemblies.** Command lines and descriptions for performing analyses are available in *Workflows.pdf*. Custom scripts are in the *scripts/* directory. Genome annotations are available in the *annotations/* directory.

### Additional File 2: SupplementaryTables.pdf

**Supplementary Tables**. Table S1: MinION sequencing metrics for Brettanomyces sequencing, Table S2: Predicted genes and gene density for the *Brettanomyces* genomes, Table S3: KEGG-annotated genes under site selection across *Brettanomyces*, Table S4: Saccharomycetaceae species used with *Brettanomyces* species in OrthoFinder

### Additional File 3: Figure_S1.tiff

**Figure S1: Synteny between haploid assemblies of *B. bruxellensis* and *B. custersianus*, visualized as a Circos plot.** Reference assembly Contigs are coloured sequentially. Alignments are coloured according to the reference assembly contigs and are layered by alignment length. The query assembly contigs are coloured grey.

### Additional File 4: Figure_S2.tiff

**Figure S2: Read-depth and SNP density over haploid assembly of *B. anomalus*, visualized as a Circos plot.** Contigs arranged by length (*i*), read-coverage histogram (blue, median coverage; red, low/high coverage) (*ii*), SNP-density (red, low; blue, high) (*iii*).

## References

1. Michel RH, McGovern PE, Badler VR: Chemical evidence for ancient beer. Nature 1992, 360(6399):24–24.

2. Fay JC, Benavides JA: Evidence for Domesticated and Wild Populations of Saccharomyces cerevisiae. PLOS Genetics 2005, 1(1):e5.

3. Edgardo A, Carolina P, Manuel R, Juanita F, Baeza J: Selection of thermotolerant yeast strains Saccharomyces cerevisiae for bioethanol production. Enzyme and Microbial Technology 2008, 43(2):120–123.

4. Marsit S, Dequin S: Diversity and adaptive evolution of Saccharomyces wine yeast: a review. FEMS Yeast Research 2015, 15(7).

5. Rozpędowska E, Hellborg L, Ishchuk OP, Orhan F, Galafassi S, Merico A, Woolfit M, Compagno C, Piškur J: Parallel evolution of the make–accumulate–consume strategy in Saccharomyces and Dekkera yeasts. Nature communications 2011, 2:302.

6. Smith MT: Chapter 89 - Brettanomyces Kufferath & van Laer (1921). In: The Yeasts (Fifth Edition). Edited by Kurtzman CP, Fell JW, Boekhout T. London: Elsevier; 2011: 983–986.

7. Péter G, Dlauchy D, Tóbiás A, Fülöp L, Podgoršek M, Čadež N: Brettanomyces acidodurans sp. nov., a new acetic acid producing yeast species from olive oil. Antonie van Leeuwenhoek 2017, 110(5):657–664.

8. Chatonnet P, Dubourdie D, Boidron J-n, Pons M: The origin of ethylphenols in wines. Journal of the Science of Food and Agriculture 1992, 60(2):165–178.

9. Chatonnet P, Dubourdieu D, Boidron JN: The Influence of *Brettanomyces/Dekkera* sp. Yeasts and Lactic Acid Bacteria on the Ethylphenol Content of Red Wines. American Journal of Enology and Viticulture 1995, 46(4):463.

10. Van Oevelen D, Spaepen M, Timmermans P, Verachtert H: MICROBIOLOGICAL ASPECTS OF SPONTANEOUS WORT FERMENTATION IN THE PRODUCTION OF LAMBIC AND GUEUZE. Journal of the Institute of Brewing 1977, 83(6):356–360.

11. Spaepen M, Van Oevelen D, Verachtert H: FATTY ACIDS AND ESTERS PRODUCED DURING THE SPONTANEOUS FERMENTATION OF LAMBIC AND GUEUZE. Journal of the Institute of Brewing 1978, 84(5):278–282.

12. Basso RF, Alcarde AR, Portugal CB: Could non-Saccharomyces yeasts contribute on innovative brewing fermentations? Food Research International 2016, 86:112–120.

13. De Souza Liberal AT, Basílio ACM, Do Monte Resende A, Brasileiro BTV, Da Silva-Filho EA, De Morais JOF, Simões DA, De Morais Jr MA: Identification of Dekkera bruxellensis as a major contaminant yeast in continuous fuel ethanol fermentation. Journal of Applied Microbiology 2007, 102(2):538–547.

14. Reis ALS, de Fátima Rodrigues de Souza R, Baptista Torres RRN, Leite FCB, Paiva PMG, Vidal EE, de Morais MA: Oxygen-limited cellobiose fermentation and the characterization of the cellobiase of an industrial Dekkera/Brettanomyces bruxellensis strain. SpringerPlus 2014, 3(1):38.

15. Curtin CD, Borneman AR, Chambers PJ, Pretorius IS: De-novo assembly and analysis of the heterozygous triploid genome of the wine spoilage yeast Dekkera bruxellensis AWRI1499. PLoS One 2012, 7(3):e33840.

16. Albertin W, Panfili A, Miot-Sertier C, Goulielmakis A, Delcamp A, Salin F, Lonvaud-Funel A, Curtin C, Masneuf-Pomarede I: Development of microsatellite markers for the rapid and reliable genotyping of Brettanomyces bruxellensis at strain level. Food Microbiology 2014, 42:188–195.

17. Borneman AR, Zeppel R, Chambers PJ, Curtin CD: Insights into the Dekkera bruxellensis Genomic Landscape: Comparative Genomics Reveals Variations in Ploidy and Nutrient Utilisation Potential amongst Wine Isolates. PLOS Genetics 2014, 10(2):e1004161.

18. Crauwels S, Zhu B, Steensels J, Busschaert P, De Samblanx G, Marchal K, Willems KA, Verstrepen KJ, Lievens B: Assessing Genetic Diversity among &lt;span class=&quot;named-content genus-species&quot; id=&quot;named-content-1&quot;&gt;Brettanomyces&lt;/span&gt; Yeasts by DNA Fingerprinting and Whole-Genome Sequencing. Applied and Environmental Microbiology 2014, 80(14):4398.

19. Varela C, Lleixà J, Curtin C, Borneman A: Development of a genetic transformation toolkit for Brettanomyces bruxellensis. FEMS Yeast Research 2018, 18(7).

20. Varela C, Bartel C, Roach M, Borneman A, Curtin C: Brettanomyces bruxellensis SSU1 Haplotypes Confer Different Levels of Sulfite Tolerance When Expressed in a Saccharomyces cerevisiae SSU1 Null Mutant. Applied and Environmental Microbiology 2019, 85(4):e02429–02418.

21. Fournier T, Gounot J-S, Freel K, Cruaud C, Lemainque A, Aury J-M, Wincker P, Schacherer J, Friedrich A: High-Quality de Novo Genome Assembly of the Dekkera bruxellensis Yeast Using Nanopore MinION Sequencing. G3: Genes|Genomes|Genetics 2017, 7(10):3243–3250.

22. Vervoort Y, Herrera-Malaver B, Mertens S, Guadalupe Medina V, Duitama J, Michiels L, Derdelinckx G, Voordeckers K, Verstrepen KJ: Characterization of the recombinant Brettanomyces anomalus β-glucosidase and its potential for bioflavouring. Journal of Applied Microbiology 2016, 121(3):721–733.

23. Koren S, Phillippy AM: One chromosome, one contig: complete microbial genomes from long-read sequencing and assembly. Current Opinion in Microbiology 2015, 23:110–120.

24. Jain M, Koren S, Miga KH, Quick J, Rand AC, Sasani TA, Tyson JR, Beggs AD, Dilthey AT, Fiddes IT, et al: Nanopore sequencing and assembly of a human genome with ultra-long reads. Nature Biotechnology 2018, 36:338.

25. Roach MJ, Schmidt SA, Borneman AR: Purge Haplotigs: allelic contig reassignment for third-gen diploid genome assemblies. BMC Bioinformatics 2018, 19(1):460.

26. Albalat R, Cañestro C: Evolution by gene loss. Nature Reviews Genetics 2016, 17:379.

27. Keeling PJ, Slamovits CH: Causes and effects of nuclear genome reduction. Current Opinion in Genetics & Development 2005, 15(6):601–608.

28. van der Walt J: Brettanomyces custersianus Nov. spec. Antonie Van Leeuwenhoek 1961, 27:332–336.

29. Kolfschoten GA, Yarrow D: Brettanomyces naardenensis, a new yeast from soft drinks. Antonie van Leeuwenhoek 1970, 36(1):458–460.

30. Smith MT, van Grinsven AM: Dekkera anomala sp. nov., the teleomorph of Brettanomyces anomalus, recovered from spoiled soft drinks. Antonie Van Leeuwenhoek 1984, 50(2):143–148.

31. Boekhout T, Kurtzman CP, O’Donnell K, Smith MT: Phylogeny of the yeast genera Hanseniaspora (anamorph Kloeckera), Dekkera (anamorph Brettanomyces), and Eeniella as inferred from partial 26S ribosomal DNA nucleotide sequences. International journal of systematic bacteriology 1994, 44(4):781–786.

32. Yamada Y, Matsuda M, Maeda K, Mikata K: The Phylogenetic Relationships of Species of the Genus Dekkera van der Walt Based on the Partial Sequences of 18S and 26S Ribosomal RNAs (Saccharomycetaceae). Bioscience, Biotechnology, and Biochemistry 1994, 58(10):1803–1808.

33. Yamada Y, Matsuda M, Mikata K: The phylogenetic relationships ofEeniella nana Smith, Batenburg-van der Vegte et Scheffers based on the partial sequences of 18S and 26S ribosomal RNAs (Candidaceae). Journal of Industrial Microbiology 1995, 14(6):456–460.

34. Röder C, König H, Fröhlich J: Species-specific identification of Dekkera/Brettanomyces yeasts by fluorescently labeled DNA probes targeting the 26S rRNA. FEMS Yeast Research 2007, 7(6):1013–1026.

35. Landry CR, Oh J, Hartl DL, Cavalieri D: Genome-wide scan reveals that genetic variation for transcriptional plasticity in yeast is biased towards multi-copy and dispensable genes. Gene 2006, 366(2):343–351.

36. Hellborg L, Piskur J: Complex nature of the genome in a wine spoilage yeast, Dekkera bruxellensis. Eukaryotic cell 2009, 8(11):1739–1749.

37. Vigentini I, De Lorenzis G, Picozzi C, Imazio S, Merico A, Galafassi S, Piskur J, Foschino R: Intraspecific variations of Dekkera/Brettanomyces bruxellensis genome studied by capillary electrophoresis separation of the intron splice site profiles. International journal of food microbiology 2012, 157(1):6–15.

38. Ishchuk OP, Vojvoda Zeljko T, Schifferdecker AJ, Mebrahtu Wisen S, Hagstrom AK, Rozpedowska E, Rordam Andersen M, Hellborg L, Ling Z, Sibirny AA et al: Novel Centromeric Loci of the Wine and Beer Yeast Dekkera bruxellensis CEN1 and CEN2. PLoS One 2016, 11(8):e0161741.

39. Avramova M, Cibrario A, Peltier E, Coton M, Coton E, Schacherer J, Spano G, Capozzi V, Blaiotta G, Salin F et al: Brettanomyces bruxellensis population survey reveals a diploid-triploid complex structured according to substrate of isolation and geographical distribution. Scientific Reports 2018, 8(1):4136.

40. Marsit S, Leducq J-B, Durand É, Marchant A, Filteau M, Landry CR: Evolutionary biology through the lens of budding yeast comparative genomics. Nature Reviews Genetics 2017, 18:581.

41. Dunn B, Richter C, Kvitek DJ, Pugh T, Sherlock G: Analysis of the Saccharomyces cerevisiae pan-genome reveals a pool of copy number variants distributed in diverse yeast strains from differing industrial environments. Genome Research 2012, 22(5):908–924.

42. Gallone B, Steensels J, Prahl T, Soriaga L, Saels V, Herrera-Malaver B, Merlevede A, Roncoroni M, Voordeckers K, Miraglia L et al: Domestication and Divergence of Saccharomyces cerevisiae Beer Yeasts. Cell 2016, 166(6):1397–1410.e1316.

43. Gonçalves M, Pontes A, Almeida P, Barbosa R, Serra M, Libkind D, Hutzler M, Gonçalves P, Sampaio José P: Distinct Domestication Trajectories in Top-Fermenting Beer Yeasts and Wine Yeasts. Current Biology 2016, 26(20):2750–2761.

44. Harris V, Ford CM, Jiranek V, Grbin PR: Survey of enzyme activity responsible for phenolic off-flavour production by Dekkera and Brettanomyces yeast. Appl Microbiol Biotechnol 2009, 81(6):1117–1127.

45. Ault RG, Newton R: Spoilage Organisms in Brewing. In: Modern Brewing Technology. Edited by Findlay WPK. London: The Macmillan Press; 1971: 164-197.

46. Reuber BE, Karl C, Reimann SA, Mihalik SJ, Dodt G: Cloning and Functional Expression of a Mammalian Gene for a Peroxisomal Sarcosine Oxidase. Journal of Biological Chemistry 1997, 272(10):6766–6776.

47. Wagner MA, Jorns MS: Monomeric Sarcosine Oxidase: 2. Kinetic Studies with Sarcosine, Alternate Substrates, and a Substrate Analogue. Biochemistry 2000, 39(30):8825–8829.

48. Yoshida N, Akazawa S-I, Katsuragi T, Tani Y: Characterization of two fructosyl-amino acid oxidase homologs of Schizosaccharomyces pombe. Journal of Bioscience and Bioengineering 2004, 97(4):278–280.

49. Nishiya Y, Nakano S, Kawamura K: Monomeric sarcosine oxidase acts on both L- and D-substrates. 生物試料分析 = Journal of analytical bio-science 2012, 35(5):426–430.

50. Conterno L, Joseph CML, Arvik TJ, Henick-Kling T, Bisson LF: Genetic and Physiological Characterization of Brettanomyces bruxellensis Strains Isolated from Wines. American Journal of Enology and Viticulture 2006, 57(2):139.

51. Woolfit M, Rozpędowska E, Piškur J, Wolfe KH: Genome Survey Sequencing of the Wine Spoilage Yeast &lt;em&gt;Dekkera (Brettanomyces) bruxellensis&lt;/em&gt. Eukaryotic cell 2007, 6(4):721.

52. Parente DC, Cajueiro DBB, Moreno ICP, Leite FCB, De Barros Pita W, De Morais Jr MA: On the catabolism of amino acids in the yeast Dekkera bruxellensis and the implications for industrial fermentation processes. Yeast 2018, 35(3):299–309.

53. Ough CS, Stashak RM: Further Studies on Proline Concentration in Grapes and Wines. American Journal of Enology and Viticulture 1974, 25(1):7.

54. Jin H, Ferguson K, Bond M, Kavanagh T, Hawthorne D: Malt nitrogen parameters and yeast fermentation behaviour. In: PROCEEDINGS OF THE CONVENTION-INSTITUTE OF BREWING ASIA PACIFIC SECTION: 1996; 1996: 44-50.

55. Gorinstein S, Zemser M, Vargas-Albores F, Ochoa JL, Paredes-Lopez O, Scheler C, Salnikow J, Martin-Belloso O, Trakhtenberg S: Proteins and amino acids in beers, their contents and relationships with other analytical data. Food Chemistry 1999, 67(1):71–78.

56. Crauwels S, Van Assche A, de Jonge R, Borneman AR, Verreth C, Troels P, De Samblanx G, Marchal K, Van de Peer Y, Willems KA et al: Comparative phenomics and targeted use of genomics reveals variation in carbon and nitrogen assimilation among different Brettanomyces bruxellensis strains. Appl Microbiol Biotechnol 2015, 99(21):9123–9134.

57. Marsit S, Sanchez I, Galeote V, Dequin S: Horizontally acquired oligopeptide transporters favour adaptation of Saccharomyces cerevisiae wine yeast to oenological environment. Environmental Microbiology 2016, 18(4):1148–1161.

58. Camarasa C, Bigey F, Marsit S, Dequin S, Nidelet T, Galeote V, Legras J-L, Sanchez I, Couloux A, Guy J et al: Adaptation of S. cerevisiae to Fermented Food Environments Reveals Remarkable Genome Plasticity and the Footprints of Domestication. Molecular Biology and Evolution 2018, 35(7):1712–1727.

59. Feurtey A, Stukenbrock EH: Interspecific Gene Exchange as a Driver of Adaptive Evolution in Fungi. Annual Review of Microbiology 2018, 72(1):377–398.

60. Hall C, Brachat S, Dietrich FS: Contribution of Horizontal Gene Transfer to the Evolution of &lt;em&gt;Saccharomyces cerevisiae&lt;/em&gt. Eukaryotic cell 2005, 4(6):1102.

61. Lin Z, Li W-H: Expansion of Hexose Transporter Genes Was Associated with the Evolution of Aerobic Fermentation in Yeasts. Molecular Biology and Evolution 2010, 28(1):131–142.

62. Monerawela C, James TC, Bond U, Wolfe KH: Loss of lager specific genes and subtelomeric regions define two different Saccharomyces cerevisiae lineages for Saccharomyces pastorianus Group I and II strains. FEMS Yeast Research 2015, 15(2).

63. Steenwyk J, Rokas A: Extensive Copy Number Variation in Fermentation-Related Genes Among &lt;em&gt;Saccharomyces cerevisiae&lt;/em&gt; Wine Strains. *G3*: Genes|Genomes|Genetics 2017, 7(5):1475.

64. Yue J-X, Li J, Aigrain L, Hallin J, Persson K, Oliver K, Bergström A, Coupland P, Warringer J, Lagomarsino MC et al: Contrasting evolutionary genome dynamics between domesticated and wild yeasts. Nature genetics 2017, 49(6):913–924.

65. Chevreux B, Wetter T, Suhai S: Genome Sequence Assembly Using Trace Signals and Additional Sequence Information. In: Computer Science and Biology: Proceedings of the German Conference on Bioinformatics (GCB) 99: 1999; 1999: 45–56.

66. Koren S, Walenz BP, Berlin K, Miller JR, Bergman NH, Phillippy AM: Canu: scalable and accurate long-read assembly via adaptive k-mer weighting and repeat separation. Genome Research 2017, 27(5):722–736.

67. Li H: Minimap2: pairwise alignment for nucleotide sequences. Bioinformatics (Oxford, England) 2018, 34(18):3094–3100.

68. Loman NJ, Quick J, Simpson JT: A complete bacterial genome assembled de novo using only nanopore sequencing data. Nature Methods 2015, 12:733.

69. Li H: Aligning sequence reads, clone sequences and assembly contigs with BWA-MEM. arXiv:13033997v1 2013.

70. Langmead B, Salzberg SL: Fast gapped-read alignment with Bowtie 2. Nature methods 2012, 9(4):357–359.

71. Walker BJ, Abeel T, Shea T, Priest M, Abouelliel A, Sakthikumar S, Cuomo CA, Zeng Q, Wortman J, Young SK et al: Pilon: An Integrated Tool for Comprehensive Microbial Variant Detection and Genome Assembly Improvement. PLOS ONE 2014, 9(11):e112963.

72. Koboldt DC, Chen K, Wylie T, Larson DE, McLellan MD, Mardis ER, Weinstock GM, Wilson RK, Ding L: VarScan: variant detection in massively parallel sequencing of individual and pooled samples. *Bioinformatics (Oxford*, England) 2009, 25(17):2283–2285.

73. Edge P, Bafna V, Bansal V: HapCUT2: robust and accurate haplotype assembly for diverse sequencing technologies. Genome research 2017, 27(5):801–812.

74. Patterson M, Marschall T, Pisanti N, van Iersel L, Stougie L, Klau GW, Schönhuth A: WhatsHap: Weighted Haplotype Assembly for Future-Generation Sequencing Reads. Journal of Computational Biology 2015, 22(6):498–509.

75. Kurtz S, Phillippy A, Delcher AL, Smoot M, Shumway M, Antonescu C, Salzberg SL: Versatile and open software for comparing large genomes. Genome biology 2004, 5(2):R12–R12.

76. Gurevich A, Tesler G, Vyahhi N, Saveliev V: QUAST: quality assessment tool for genome assemblies. *Bioinformatics (Oxford*, England) 2013, 29(8):1072–1075.

77. Kriventseva EV, Zdobnov EM, Simão FA, Ioannidis P, Waterhouse RM: BUSCO: assessing genome assembly and annotation completeness with single-copy orthologs. *Bioinformatics (Oxford*, England) 2015, 31(19):3210–3212.

78. Stanke M, Keller O, Gunduz I, Hayes A, Waack S, Morgenstern B: AUGUSTUS: ab initio prediction of alternative transcripts. Nucleic acids research 2006, 34(Web Server issue):W435-W439.

79. Kanehisa M, Sato Y, Morishima K: BlastKOALA and GhostKOALA: KEGG Tools for Functional Characterization of Genome and Metagenome Sequences. Journal of molecular biology 2016, 428(4):726–731.

80. Jones P, Binns D, Chang H-Y, Fraser M, Li W, McAnulla C, McWilliam H, Maslen J, Mitchell A, Nuka G et al: InterProScan 5: genome-scale protein function classification. *Bioinformatics (Oxford*, England) 2014, 30(9):1236–1240.

81. Emms DM, Kelly S: OrthoFinder: solving fundamental biases in whole genome comparisons dramatically improves orthogroup inference accuracy. Genome Biology 2015, 16(1):157.

82. Edgar RC: MUSCLE: multiple sequence alignment with high accuracy and high throughput. Nucleic acids research 2004, 32(5):1792–1797.

83. Suyama M, Torrents D, Bork P: PAL2NAL: robust conversion of protein sequence alignments into the corresponding codon alignments. Nucleic acids research 2006, 34(Web Server issue):W609-W612.

84. Paradis E, Claude J, Strimmer K: APE: Analyses of Phylogenetics and Evolution in R language. Bioinformatics (Oxford, England) 2004, 20(2):289–290.

85. Schliep KP: phangorn: phylogenetic analysis in R. Bioinformatics (Oxford, England) 2011, 27(4):592–593.

86. Krzywinski M, Schein J, Birol İ, Connors J, Gascoyne R, Horsman D, Jones SJ, Marra MA: Circos: An information aesthetic for comparative genomics. Genome Research 2009, 19(9):1639–1645.

87. Heymans K, Kuiper M, Maere S: BiNGO: a Cytoscape plugin to assess overrepresentation of Gene Ontology categories in Biological Networks. Bioinformatics (Oxford, England) 2005, 21(16):3448–3449.

88. Gouy M, Gascuel O, Guindon S: SeaView Version 4: A Multiplatform Graphical User Interface for Sequence Alignment and Phylogenetic Tree Building. Molecular Biology and Evolution 2009, 27(2):221–224.

89. Yang Z: PAML 4: Phylogenetic Analysis by Maximum Likelihood. Molecular Biology and Evolution 2007, 24(8):1586–1591.

90. Pruitt KD, Tatusova T, Klimke W, Maglott DR: NCBI Reference Sequences: current status, policy and new initiatives. Nucleic acids research 2009, 37(Database issue):D32-D36.

