## Additional File 1 for "New genome assemblies reveal patterns of domestication and adaptation across *Brettanomyces* (*Dekkera*) species": BUSCO_short_summary_B.custersianus.txt

# BUSCO version is: 3.0.2 
# The lineage dataset is: saccharomyceta_odb9 (Creation date: 2016-02-13, number of species: 70, number of BUSCOs: 1759)
# To reproduce this run: python /home/mike/anaconda3/envs/busco/bin/run_busco -i ../assembly-gene-annotation/B.custersianus.prot.fasta -o custersianus -l /home/mike/BUSCO/datasets/saccharomyceta_odb9/ -m proteins -c 16
#
# Summarized benchmarking in BUSCO notation for file ../assembly-gene-annotation/B.custersianus.prot.fasta
# BUSCO was run in mode: proteins

	C:88.2%[S:87.3%,D:0.9%],F:6.4%,M:5.4%,n:1759

	1551	Complete BUSCOs (C)
	1535	Complete and single-copy BUSCOs (S)
	16	Complete and duplicated BUSCOs (D)
	113	Fragmented BUSCOs (F)
	95	Missing BUSCOs (M)
	1759	Total BUSCO groups searched
