## Additional File 1 for "New genome assemblies reveal patterns of domestication and adaptation across *Brettanomyces* (*Dekkera*) species": add_annotations.pl

```
#!/usr/bin/env perl

use strict;
use warnings;
use Data::Dumper;

# one off script to add ortho group annotations to gff file

my $usage="
Usage:
perl add_annotations.pl  prot.fofn  blast.fofn  uniprot_sprot.names.tsv  gff.fofn  orthofinder/Results_Jul13/Orthogroups.csv

prot.fofn                   fofn of prot augustus predicted proteins (FASTA)
blast.fofn                  fofn of blastp results agains uniprotKB
uniprot_sprot.names.tsv     table of uniprot seq IDs to seq names
gff.fofn                    fofn of augustus gff annotations for groups in orthogroups file
Orthogroups.csv             orthogroup annotations from orthofinder

";

my %orth;       # $orth{sample}{gene} = group
my %prot;       # $prot{sample}{gene} = prot name
my %groups;     # $groups{index} = sample
my %uniprot;    # $uniprot{seqID} = prot name


my $prot = shift or die $usage;
my $blast = shift or die $usage;
my $prot_names = shift or die $usage;
my $gffs = shift or die $usage;
my $orthos = shift or die $usage;


# parse the orthogroups CSV
open my $CSV, '<', $orthos or die;
{
    my $head = <$CSV>;
    my @l=split/\t/,$head;
    for my $i (1..$#l){
        $l[$i]=~s/\s+//g;
        $l[$i]=~s/\.prot//;
        $l[$i]=~s/\.FIXED//;
        $groups{$i}=$l[$i];
    }
}

while(<$CSV>){
    my@l=split/\t/;
    for my $i (keys %groups){
        my@g=split/,/,$l[$i];
        for my $gene (@g){
            $gene =~ s/\s+//g;
            $orth{$groups{$i}}{$gene} = $l[0];
        }
    }
}
close$CSV;


# parse the uniprot name table
open my $UT, '<', $prot_names or die;
while(<$UT>){
    my@l=split/\t/;
    $l[1]=~s/\s+$//;
    $uniprot{$l[0]} = $l[1];
}
close$UT;


# parse the blast hits
open my $BH, '<', $blast or die;
while(my $f = <$BH>){
    $f =~ s/\.uniprotkb\.outfmt6\.gz//;
    $f =~ s/\s+//g;
    open my $TMP, "zcat $f.uniprotkb.outfmt6.gz |" or die;
    while(<$TMP>){
        my@l=split/\s+/;
        ($prot{$f}{$l[0]}) || ($prot{$f}{$l[0]} = $uniprot{$l[1]});
    }
    close$TMP;
}
close$BH;


# parse the protein seqs
open my $PR, '<', $prot or die;
while(my $f = <$PR>){
    $f=~s/\s+//g;
    $f =~ s/\.FIXED//;
    $f =~ s/\.FALC//;
    $f=~s/\.prot\.fasta//;
    open my $TMP, '<', "$f.prot.fasta" or die;
    open my $PO, '>', "$f.annot.prot.fasta" or die;
    while(<$TMP>){
        if(m/^>/){
            my$id=$_;
            $id=~s/>//g;
            $id=~s/\s+//g;
            print $PO ">$id ";
            print $PO (($prot{$f}{$id}) ? ("$prot{$f}{$id};") : ("hypothetical protein;"));
            print $PO (($orth{$f}{$id}) ? ("$orth{$f}{$id}\n") : ("\n"));
        } else {
            print $PO $_;
        }
    }
    close$PO;
    close$TMP;
}
close$PR;


# parse the gffs
open my $GF, '<', $gffs or die;
while(my $f = <$GF>){
    $f=~s/\s+//g;
    $f=~s/.gff$//;
    open my $TMP, '<', "$f.gff" or die;
    open my $TO, '>', "$f.annot.gff" or die;
    while(<$TMP>){
        next if (m/^#/);
        my @l=split/\s+/;
        if($l[2]eq'gene'){
            my$id=$l[8];
            $l[8].=(($orth{$f}{$id})?(";$orth{$f}{$id};"):(";;"));
            $l[8].=(($prot{$f}{$id})?($prot{$f}{$id}):("hypothetical protein"));
        }
        ($_.="\t")for(@l);
        $l[$#l]=~s/\t/\n/;
        print $TO @l;
    }
    close$TO;
    close$TMP;
}
close$GF;
```
