## Additional File 1 for "New genome assemblies reveal patterns of domestication and adaptation across *Brettanomyces* (*Dekkera*) species": codemlOutputScan.pl

```
use strict; 
use warnings;

# one-off script to return loglikelihood ratios above cutoffs

my $fofn = shift or die;

open my $FO, '<', $fofn or die;

while(<$FO>){
    chomp;
    my @lnL = split/\s+/,`grep lnL $_ | awk '{print\$5}'`;
    my $lrt = 2 * ($lnL[1]-$lnL[0]);
    if($lrt > 5.99){
        $_ =~ s/\..*//;
        $_ =~ s/.*\///;
        print "$_\t$lrt\n";
    }
}
```
