## Additional File 1 for "New genome assemblies reveal patterns of domestication and adaptation across *Brettanomyces* (*Dekkera*) species": combineMSAs.pl

```
use strict;
use warnings;

my $usage = "
perl combineMSAs.pl  MSA.fofn  samples.list
";

my $fofn = shift or die $usage;
my $list = shift or die $usage;

# vars
my @samples;
my %align;

# read in list
read_samples();

# concatenate seqs
open my $TMP, '<', $fofn or die;
while(<$TMP>){
    chomp;
    open my $ALI, '<', $_ or die;
    my $smpl;
    my %tmp;
    while(<$ALI>){
        if($_=~/^>(.*)\s/){
            $smpl=$1;
        } else {
            $tmp{$smpl} .= $_;
        }
    }
    close $ALI;
    if(scalar(keys(%tmp))==scalar(@samples)){
        for my $k (keys %tmp){
            $align{$k} .= $tmp{$k};
        }
    }
}
close $TMP;

# dump
for my $k (keys %align){
    print ">$k\n", $align{$k};
}
exit;


sub read_samples {
    open my $TMP, '<', $list or die;
    while(<$TMP>){
        chomp;
        ($_) and push @samples, $_;
    }
    close $TMP;
}
```
