## Additional File 1 for "New genome assemblies reveal patterns of domestication and adaptation across *Brettanomyces* (*Dekkera*) species": HGT-check.pl

```
use strict;
use warnings;
use Data::Dumper;

# one-off script to check blast output for potential horizontal gene transfer

my $usage = "
USAGE:
perl HGT-check.pl  fungi.outfmt6.gz  bact.outfmt6.gz  > candidates.tsv
";


my $fungi = shift or die $usage;
my $bact = shift or die $usage;

(-s $fungi) and (-s $bact) or die $usage;

my %bact;       # $bact{gene}{b} = evalue
                #      {gene}{f} = evalue

open my $BAC, "zcat $bact |" or die "failed to open pipe to $bact";
while(<$BAC>){
    my@l=split/\s+/;
    $bact{$l[0]}{b} = $l[5];
}
close$BAC;

open my $FNG, "zcat $fungi |" or die "failed to open pipe to $fungi";
while(<$FNG>){
    my@l=split/\s+/;
    if(defined($bact{$l[0]})){
        $bact{$l[0]}{f} = $l[5];
    }
}
close$FNG;


for my $gene (sort keys %bact){
    if(!(defined($bact{$gene}{f}))){
        $bact{$gene}{f} = 1;
    }
    if($bact{$gene}{b} < $bact{$gene}{f}){
        print STDOUT "$gene\t$bact{$gene}{b}\t$bact{$gene}{f}\n";
    }
}
```
