## Additional File 1 for "New genome assemblies reveal patterns of domestication and adaptation across *Brettanomyces* (*Dekkera*) species": PipeUtils.pm

```
package PipeUtils;

# Copyright (c) 2017 Michael Roach (Australian Wine Research Institute)
#
# Permission is hereby granted, free of charge, to any person obtaining a copy
# of this software and associated documentation files (the "Software"), to deal
# in the Software without restriction, including without limitation the rights
# to use, copy, modify, merge, publish, distribute, sublicense, and/or sell
# copies of the Software, and to permit persons to whom the Software is
# furnished to do so, subject to the following conditions:
# 
# The above copyright notice and this permission notice shall be included in all
# copies or substantial portions of the Software.
#
# THE SOFTWARE IS PROVIDED "AS IS", WITHOUT WARRANTY OF ANY KIND, EXPRESS OR
# IMPLIED, INCLUDING BUT NOT LIMITED TO THE WARRANTIES OF MERCHANTABILITY,
# FITNESS FOR A PARTICULAR PURPOSE AND NONINFRINGEMENT. IN NO EVENT SHALL THE
# AUTHORS OR COPYRIGHT HOLDERS BE LIABLE FOR ANY CLAIM, DAMAGES OR OTHER
# LIABILITY, WHETHER IN AN ACTION OF CONTRACT, TORT OR OTHERWISE, ARISING FROM,
# OUT OF OR IN CONNECTION WITH THE SOFTWARE OR THE USE OR OTHER DEALINGS IN THE
# SOFTWARE.

use strict;
use warnings;
use Time::Piece;

use Exporter 'import';

our @EXPORT = qw(msg err runcmd qruncmd check_programs check_files);


sub print_message {
    my $t = localtime;
    my $line = "[" . $t->dmy . " " . $t->hms . "] @_\n";
    print STDERR $line;
    print $::LOG $line if ($::LOG);
}

sub msg {
    print_message("INFO: @_");
}

sub err {
    print_message("ERROR: @_\n\nPIPELINE FAILURE\n");
    exit(1);
}

sub runcmd {
    my $job = shift;
    ($job->{silent}) || print_message("RUNNING: $job->{command}");
    if (system("$job->{command}") != 0){
        print_message("ERROR: Failed to run $job->{command}");
        print_message("Check $job->{logfile} for possible errors") if ($job->{logfile});
        err("Exiting due to job failure");
    } else {
        ($job->{silent}) || print_message("FINISHED: $job->{command}");
    }
}

sub qruncmd {
    system(@_) == 0 or err("Failed to run @_\n");
}

sub check_files {
    my $check=1;
    foreach(@_){
        if (!(-s $_)){
            print_message("ERROR: file \"$_\" does not exist or is empty");
            $check=0;
        }
    }
    return $check;
}

sub check_programs {
    my $chk=1;
    my $t = localtime;
    my $line = "[" . $t->dmy . " " . $t->hms . "]";
    foreach my $prog (@_){
        print STDERR "$line CHECKING $prog... ";
        my $notexists = `type $prog 2>&1 1>/dev/null || echo 1`;
        if ($notexists){
            print STDERR "ERROR: missing program $prog\n";
            $chk = 0;
        } else {
            print STDERR "OK\n";
        }
    }
    return $chk;
}
```
