## Additional File 1 for "New genome assemblies reveal patterns of domestication and adaptation across *Brettanomyces* (*Dekkera*) species": map_colors.pl

```
#!/usr/bin/env perl

use strict;
use warnings;

my %col = (
    'AWRI2804_contig2203_scaffold1' => 'set3-12-qual-2',
    'AWRI2804_contig2201_scaffold1' => 'set3-12-qual-1',
    'AWRI2804_contig2057_scaffold1' => 'set3-12-qual-3',
    'AWRI2804_contig1898_scaffold1' => 'set3-12-qual-6',
    'AWRI2804_contig1521_scaffold1' => 'set3-12-qual-5',
    'AWRI2804_contig1179_scaffold1' => 'set3-12-qual-7',
    'AWRI2804_contig795_scaffold1' => 'set3-12-qual-8',
    'AWRI2804_contig787_scaffold1' => 'set3-12-qual-9',
    'AWRI2804_contig762_scaffold1' => 'set3-12-qual-4',
    'AWRI2804_contig2285_scaffold3' => 'set3-12-qual-10',
    'AWRI2804_contig2357_scaffold4' => 'set3-12-qual-11',
    'AWRI2804_contig2276_scaffold4' => 'set3-12-qual-12',
    'tig00000001' => 'set3-5-qual-1',
    'tig00000031' => 'set3-5-qual-2',
    'tig00000034' => 'set3-5-qual-3',
    'tig00000047' => 'set3-5-qual-4',
    'tig00000627' => 'set3-5-qual-5'
);

my @cache;
my $max=0;
while(<>){
    my@l=split/\s+/;
    ($max>$l[$#l])||($max=$l[$#l]);
    ($_.=" ")for(@l);
    push@cache, "@l";
}

for (@cache){
    my@l=split/\s+/;
    $l[$#l]="z=" . int(100 - (100 * ($l[$#l]/$max)));
    $l[$#l].=",color=$col{$l[0]}";
    ($_.=" ")for(@l);
    $l[$#l]=~s/\s+/\n/;
    print @l;
}
```
