## Additional File 1 for "New genome assemblies reveal patterns of domestication and adaptation across *Brettanomyces* (*Dekkera*) species": multiCodeml.pl

```
use strict;
use warnings;
use Data::Dumper;
use threads;
use Thread::Semaphore;
use Thread::Queue;


my $usage = "
perl multiCodeml.pl  MSA.fofn  template.ctl
";

my $fofn = shift or die $usage;
my $template = shift or die $usage;


# vars
my $threads = 16;
my $ctl;


# THREADS
my $available_threads = Thread::Semaphore->new($threads);
my $writing_to_out = Thread::Semaphore->new(1);
my $queue = Thread::Queue->new();


# slurp template
$ctl = `cat $template`;

# queue jobs
queueCodeml();

# spawn workers
for (1..$threads){
    $available_threads->down(1);
    threads->create(\&codemlJob);
}

# wait on jobs
$available_threads->down($threads);
$available_threads->up($threads);
for my $thr (threads->list()){
    $thr->join();
}

# exit
exit(0);


sub queueCodeml {
    open my $TMP, '<', $fofn or die;
    while(<$TMP>){
        chomp;
        (-s $_) or die;
        $queue->enqueue($_);
    }
    close $TMP;
    $queue->end();
}


sub codemlJob {
    while (defined(my $align = $queue->dequeue())) {
        my $og = $align;
        $og =~ s/\.MSA.*//;
        $og =~ s/\.best.nuc.*//;
        $og =~ s/\.cdaln.fa//;
        
        # make the directory to run the prog
        mkdir $og or die;
        # make control file
        open my $CTL, '>', "$og/$og.ctl" or die;
        print $CTL "seqfile = ../$align\noutfile = $og.RESULT\n$ctl";
        close $CTL;
        # run the program
        my $cmd = "cd $og && printf \"\n\" | codeml $og.ctl";
        system($cmd)==0 or die;
        # save output and cleanup
        rename "$og/$og.RESULT", "$og.M8M7";
        `rm -rf $og/`;
    }
    # exit thread
    $available_threads->up(1);

    return;
}
```
