## Additional File 1 for "New genome assemblies reveal patterns of domestication and adaptation across *Brettanomyces* (*Dekkera*) species": nucmer2trans.pl

```
#!/usr/bin/env perl

use strict;
use warnings;
use Getopt::Long;
use Data::Dumper;


# inputs
my $refwind;
my $querywind;
my $coords;


my $usage = "
Usage:
Perl nucmer2trans.pl  -r ref.wind.bed  -q query.wind.bed  -c ref.query.coords

-r      genome windows for reference assembly (bedtools makewindows)
-q      genome windows for query assembly
-c      alignment coords using nucmer, delta-filter (-1), show-coords -TH

";


GetOptions (
    "ref=s" => \$refwind,
    "query=s" => \$querywind,
    "coords=s" => \$coords
) or die $usage;

($refwind)&&($querywind)&&($coords)||die $usage;

# global vars

my %ref;        # @{$ref{contig}} = ((start1,stop1),(start2,stop2), ...)
my %query;      # @{$query{contig}} = ((start1,stop1),(start2,stop2), ...)
my %counts;     # $counts{refwind}{querywind}{len} = net ali len
                # $counts{refwind}{querywind}{+} = net forward ali len
                # $counts{refwind}{querywind}{-} = net reverse ali len
my %out;

my %trn;
my @mtrn;
my @ptrn;

my $alignment_len_cutoff = 100;
my $chain_len = 20000;


## slurp windows
## iterate coords
    ## get window hits
    ## add to window counts
# iterate ref coords
    # output highest match
# iterate out coords
    # merge overlaps
    # trim clashes
    # chain gaps
    # output


# slurp window coords
open my $REF, '<', $refwind or die;
while(<$REF>){
    my@l=split/\s+/;
    # format check?
    if ($l[1] == 0){
        $l[1] = 1;
    }
    @{$ref{$l[0]}[@{$ref{$l[0]}}]} = ($l[1],$l[2]);
}
close $REF;

open my $QRY, '<', $querywind or die;
while(<$QRY>){
    my@l=split/\s+/;
    # format check?
    @{$query{$l[0]}[@{$query{$l[0]}}]} = ($l[1],$l[2]);
}
close $QRY;


# iterate alignment coords
parse_coords();


# iterate counts and call best match
itr_counts();


# merge, trim, chain
chain_output();


# more merging
merge_query();


#print Dumper(\%trn);


exit;


sub parse_coords {
    open my $CRD, '<', $coords or die;
    while(<$CRD>){
        my@l=split/\s+/;
        ($ref{$l[7]}) or die "Ref ctg $l[7] not in $refwind";
        ($query{$l[8]}) or die "Query ctg $l[8] not in $querywind";
        
        # we'll ignore alignments that overlap window edges for the moment,
        # shouldn't be a problem unless the alignments are larger than the 
        # genome windows.
        
        # get the alignment direction
        my $dir = ($l[3]>$l[2])?('+'):('-');
        
        # find the best query window
        my %qwindhits;
        my $nearest=9999999999999999999999999;
        my $key;
        my $len;
        for my $qwind (@{$query{$l[8]}}){
            if (($l[2]>=@$qwind[0])&&($l[3]<=@$qwind[1])){
                if (abs(((@$qwind[0]+@$qwind[1])/2) - $l[2]) < $nearest){
                    $nearest = abs(((@$qwind[0]+@$qwind[1])/2) - $l[2]);
                    $key = "$l[8]\t@$qwind[0]\t@$qwind[1]";
                    $len = abs ((($l[2]+$l[3])/2) - ((@$qwind[0]+@$qwind[1])/2));
                }
            }
        }
        $qwindhits{$key} = $len;
        
        
        # add alignment to ref windows, $l[0]-$l[1]
        for my $rwind (@{$ref{$l[7]}}){
            if (($l[0]>=@$rwind[0])&&($l[1]<=@$rwind[1])){
                for my $qwind (sort {$qwindhits{$b} <=> $qwindhits{$a} } keys %qwindhits){
                    my $r = \%{$counts{$l[7]}{@$rwind[0]}};
                    $$r{$qwind}{E} = @$rwind[1];
                    $$r{$qwind}{len} += $l[4];
                    $$r{$qwind}{$dir} += $l[4];
                    $$r{$qwind}{n} += 1;
                    last;
                }
            }
        }
    }
    return;
}


sub itr_counts {
    # ref contig
    for my $ctg (sort keys %counts){
        # ref window
        for my $rstart (sort keys %{$counts{$ctg}}){
            my $r = \%{$counts{$ctg}{$rstart}};
            Q: for my $qwind (sort { $$r{$b}{len} <=> $$r{$a}{len} } keys %{$r} ){
                if ($$r{$qwind}{len} > $alignment_len_cutoff){
                    my $q = \%{$counts{$ctg}{$rstart}{$qwind}};
                    ($$q{'+'}) || ($$q{'+'}=0);
                    ($$q{'-'}) || ($$q{'-'}=0);
                    my@q=split/\t/,$qwind;
                    $out{$ctg}{$rstart}{E} = $$q{E};
                    @{$out{$ctg}{$rstart}{Q}} = @q;
                    $out{$ctg}{$rstart}{D} = (($$q{'+'}>$$q{'-'})?('+'):('-'));
                    $out{$ctg}{$rstart}{L} = $$r{$qwind}{len};
                    last Q;
                } else {
                    last Q;
                }
            }
        }
    }
    return;
}


sub chain_output {
    for my $ctg (sort keys %out){
        my $cwin;
        W: for my $win (sort { $a <=> $b } keys %{$out{$ctg}}){
            if ($cwin){
                my $c = \%{$out{$ctg}{$cwin}};
                my $w = \%{$out{$ctg}{$win}};
                    # merge conditions
                    if (    
                    (@{$$c{Q}}[0] eq @{$$w{Q}}[0])     &&            # query windows same contig
                    ($$c{D} eq $$w{D}) ){                            # same direction
                        if ($$c{D}eq'+'){
                            if ( ((@{$$w{Q}}[1]<=@{$$c{Q}}[2])&&(@{$$w{Q}}[1]>=@{$$c{Q}}[1])) ||      # left or right overlap
                                 ((@{$$w{Q}}[2]<=@{$$c{Q}}[2])&&(@{$$w{Q}}[2]>=@{$$c{Q}}[1])) ) {
                                #
                                @{$$c{Q}}[2] = @{$$w{Q}}[2];
                                $$c{E} = $$w{E};
                                $$c{L} += $$w{L};
                                next W;
                                #
                            } else {
                                ### I'm not sure if we need any trimming here, trim later ###
                                print_window($ctg,$cwin);
                                $cwin=$win;
                                next W;
                            }
                        } elsif ($$c{D}eq'-'){
                            if ( ((@{$$w{Q}}[1]>=@{$$c{Q}}[2])&&(@{$$w{Q}}[1]<=@{$$c{Q}}[1])) ||      # left or right overlap
                                 ((@{$$w{Q}}[2]<=@{$$c{Q}}[2])&&(@{$$w{Q}}[2]>=@{$$c{Q}}[1])) ) {
                                #
                                @{$$c{Q}}[1] = @{$$w{Q}}[1];
                                $$c{E} = $$w{E};
                                $$c{L} += $$w{L};
                                next W;
                                #
                            } else {
                                ### trim later ###
                                print_window($ctg,$cwin);
                                $cwin=$win;
                                next W;
                            }
                        }
                    } else {
                        ### trim the ref seq windows later ###
                        print_window($ctg,$cwin);
                        $cwin=$win;
                        next W;
                    }
            } else {
                $cwin=$win;
                next W;
            }
        }
        print_window($ctg,$cwin);
    }
    return;
}


sub print_window {
    my $ctg = $_[0];
    my $start = $_[1];
    $start = 0 if ($start == 1);
    my $w = \%{$out{$_[0]}{$_[1]}};
    @{$trn{$ctg.$start}} = ($ctg,$start,$$w{E},@{$$w{Q}}[0],@{$$w{Q}}[1],@{$$w{Q}}[2],$$w{D},$$w{L});
    #print STDOUT "$ctg\t$start\t$$w{E}\t@{$$w{Q}}[0]\t@{$$w{Q}}[1]\t@{$$w{Q}}[2]\t$$w{D}\t$$w{L}\n";
    return;
}


sub merge_query {
    
    my $c;
    for my $l (sort { @{$trn{$a}}[0] cmp @{$trn{$b}}[0] || 
                      @{$trn{$a}}[3] cmp @{$trn{$b}}[3] || 
                      @{$trn{$a}}[4] <=> @{$trn{$b}}[4] } keys %trn) {
        if ($c){
            # same ref and query ctgs
            if (  (@{$trn{$l}}[0] eq @{$trn{$c}}[0]) && 
                  (@{$trn{$l}}[3] eq @{$trn{$c}}[3])    ){
                # merge conditions
                if ( # ref ovl or close enough
                        ((@{$trn{$l}}[1]<=@{$trn{$c}}[2]) && (@{$trn{$l}}[1]>=@{$trn{$c}}[1])) ||
                        ((@{$trn{$l}}[1] - $chain_len<=@{$trn{$c}}[2]) && (@{$trn{$l}}[1] - $chain_len>=@{$trn{$c}}[1])) ){
                    if ( # query ovl or close enough
                          ((@{$trn{$l}}[4]<=@{$trn{$c}}[5]) && (@{$trn{$l}}[4]>=@{$trn{$c}}[4])) ||
                          ((@{$trn{$l}}[4] - $chain_len<=@{$trn{$c}}[5]) && (@{$trn{$l}}[4] - $chain_len>=@{$trn{$c}}[4])) ){
                        # check engulfed
                        if ((@{$trn{$l}}[4]>=@{$trn{$c}}[4]) && (@{$trn{$l}}[5]<=@{$trn{$c}}[5])){
                            delete $trn{$l};
                            next;
                        } elsif (@{$trn{$l}}[6] eq @{$trn{$c}}[6]) {
                            @{$trn{$c}}[2] = @{$trn{$l}}[2];
                            @{$trn{$c}}[5] = @{$trn{$l}}[5];
                            @{$trn{$c}}[7] += @{$trn{$l}}[7];
                            delete $trn{$l};
                            next;
                        } else {
                            (print STDOUT "$_\t") for (@{$trn{$c}});
                            print STDOUT "\n";
                            $c = $l;
                            next;
                        }
                    } else {
                        (print STDOUT "$_\t") for (@{$trn{$c}});
                        print STDOUT "\n";
                        $c = $l;
                        next;
                    }
                } else {
                    (print STDOUT "$_\t") for (@{$trn{$c}});
                    print STDOUT "\n";
                    $c = $l;
                    next;
                }
            } else {
                (print STDOUT "$_\t") for (@{$trn{$c}});
                print STDOUT "\n";
                $c = $l;
                next;
            }
        } else {
            $c = $l;
            next;
        }
    }
}
```
