## Additional File 1 for "New genome assemblies reveal patterns of domestication and adaptation across *Brettanomyces* (*Dekkera*) species": OG-species-tree.pl

```
use strict;
use warnings;
use Data::Dumper;

# one-off script to pull sequences for an OG, for specific species

my $usage = "
USAGE:
perl OG-species-tree.pl  seqsDir/  OGs.tsv  species.list  OGs.list

seqsDir         directory of sequences used by orthofinder
OGs.tsv         orthofinder output file
species.list    list of species to consider
OGs.list        list of OGs for which to pull seqs

outputs:
<OG>.faa        MSA of seqs for OG for species
";

my $inDir = shift or die $usage;
my $ogTSV = shift or die $usage;
my $spList = shift or die $usage;
my $ogList = shift or die $usage;

#(-d $inDir) and (-s $ogTSV) and (-s $spList) and (-s $ogList) or die $usage;

# global vars
my @species;
my @orthogroups;
my %genes;          # @{$gene{species}{og}} = list
my %spIndex;        # $spIndex{species} = index

# read species
@species = read_list($spList);

# read orthogroups
@orthogroups = read_list($ogList);

# pull the gene IDs for each OG for each species
read_ortho_TSV();

# pull the sequences and print the files
pull_seqs();


sub read_list {
    my $f = $_[0];
    my @push;
    open my $TMP, '<', $f or die;
    while(<$TMP>){
        chomp;
        push @push, $_ if ($_);
    }
    close $TMP;
    return @push;
}

sub read_ortho_TSV {
    open my $TMP, '<', $ogTSV or die;
    
    # read the header into the index hash
    my $header = <$TMP>;
    chomp($header);
    my@h=split/\t/,$header;
    for my $i (1..$#h){
        if ( grep( /$h[$i]/, @species ) ){
            $spIndex{$h[$i]}=$i;
        }
    }
    
    # read the TSV and pull the gene IDs
    while(<$TMP>){
        chomp;
        my@l=split/\t/;
        if ( grep( /$l[0]/, @orthogroups ) ){
            for my $i (keys %spIndex){
                if($l[$spIndex{$i}]){
                    my @g = split/,/,$l[$spIndex{$i}];
                    @{$genes{$i}{$l[0]}} = @g;
                }
            }
        }
    }
}


sub pull_seqs {
    for my $og (@orthogroups){
        open my $TMP, '>', "$og.faa" or die;
        for my $sp (@species){
            my $gens;
            ($gens .= "$_ ") for (@{$genes{$sp}{$og}});
            if($gens){
                my $seqs = `samtools faidx $inDir/$sp.faa $gens | sed 's/>/>$sp./'`;
                print $TMP $seqs;
            }
        }
        close $TMP;
    }
}
```
