## Additional File 1 for "New genome assemblies reveal patterns of domestication and adaptation across *Brettanomyces* (*Dekkera*) species": orient_haplotigs.pl

```
#!/usr/bin/env perl

use strict;
use warnings;
use Getopt::Long;
use FindBin qw($Bin);
use lib "$Bin/../lib";
use PipeUtils;
use Data::Dumper;
use List::Util qw(min max);

# This script will use the placement information from a purge_haplotigs ncbiplace run to create an oriented haplotigs
# FASTA file. This can be used to order the haplotigs to the primary contigs, or, to orient a haploid assembly to 
# a reference genome. Contigs are ordered by their median alignment centroid from a global-filtered nucmer alignment. 
# The median alignment is the alignment that contains the median aligned base, hence alignments have a pesudo weighting.

# REFERENCE     --------------------------------------------------------------------------------------------------
# CONTIG            ----        ------                 ------------------------  ----------------- ---------------
# MEDIAN BASE                                                                ^                                    
# CONTIG'S REFERENCE POINT                                         ^


# pre flight checks
check_programs("samtools");


# global vars
my $haplotigs;
my $primaries;
my $placements;
my $TMP = "tmp_purge_haplotigs/NCBIPLACE";
my $out = "oriented";
my $scaffold;
my $dont_rename;
my $tac_contigs;

my %alignments;     # $alignments{primary}{haplotig}{D} = direction (+/-)
                    #                               {C} = median alignment centroid - relative to primary
my @primaries;      # This is purely to order the haplotig ouput

my $usage = "

USAGE:
orient_haplotigs.pl  -p primary_contigs.fa  -h haplotigs.fa  -n ncbi_placements.tsv  [ -d temp_ncbi_placements_dir -o outprefix]

REQUIRED:
-p      primary contigs FASTA
-h      haplotigs FASTA
-n      ncbi placement file from purge_haplotigs ncbiplace

OPTIONAL:
-d      temp directory produced from purge_haplotigs ncbiplace, DEFAULT = $TMP
-o      output file prefix, DEFAULT = $out
-s      pseudo scaffold the haplotigs together (also produces a bed file annotation of the contigs)
-r      don't rename contigs
-t      tac unaligned contigs on the end
";


# parse and check args
GetOptions (
    "p=s" => \$primaries,
    "h=s" => \$haplotigs,
    "n=s" => \$placements,
    "d=s" => \$TMP,
    "o=s" => \$out,
    "s" => \$scaffold,
    "r" => \$dont_rename,
    "t" => \$tac_contigs
) or die $usage;

($primaries) && ($haplotigs) && ($placements) || err($usage);


# cleanup old output and open output filehandles
((-s "$_") && (unlink $_)) for  ("$out.fa", "$out.bed");
open my $OFH, ">", "$out.fa" or err("Failed to open $out.fa for writing");
($scaffold) && (open my $BED, ">", "$out.bed" or err("Failed to open $out.bed for writing"));


# check FASTAs
check_files($haplotigs, $primaries);
( (-s "$_.fai") || runcmd({ command => "samtools faidx $_" }) ) for ($primaries, $haplotigs);


# get the primary contig order
open my $PFH, "<", "$primaries.fai" or err("Failed to open $primaries.fai for reading");
while(<$PFH>){
    my @l = split(/\s+/,$_);
    push @primaries, $l[0];
}
close $PFH;


# parse the placements file
open my $PL, $placements or err("failed to open $placements for reading");

while(<$PL>){
    next if ($_ =~ /^#/);
    my @l = split(/\s+/,$_);
    $alignments{$l[4]}{$l[2]}{D} = $l[5];
}

close $PL;


# get the centroid for each haplotig
for my $primary (keys(%alignments)){
    next if ($primary eq "na");
    for my $haplotig (keys(%{$alignments{$primary}})){
        my $h = $alignments{$primary}{$haplotig};
        my $htig_file = $haplotig;
        $htig_file =~ s/\|.+//;
        (-s "$TMP/$htig_file.$primary.coords") or err("Missing file $TMP/$htig_file.$primary.coords");
        open my $COO, "$TMP/$htig_file.$primary.coords" or err("Failed to open $TMP/$htig_file.$primary.coords for reading");
        
        # parse the coords file for the haplotig
        my @centroids;
        my @bases;
        my $totalbases;
        while(<$COO>){
            next if ($_ !~ /^\d/);
            my @l = split(/\s+/,$_);
            push @centroids, (($l[2] + $l[3])/2);
            push @bases, $l[5];
            $totalbases += $l[5];
        }
        close $COO;
        
        # grab the median alignment centroid
        my $runningbases;
        for (my $i=0; $i < @centroids; $i++){
            $runningbases += $bases[$i];
            if ($runningbases > (0.5 * $totalbases)){
                $alignments{$primary}{$haplotig}{C} = $centroids[$i];
                last;
            } 
        }
    }
}


# iterate the contigs and print out the oriented haplotigs
for my $primary (@primaries){
    
    # scaffolding options (ignored otherwise)
    my $scaffold_seq;
    my $bed_position=0;
    
    my $count=0;
    next if ($primary eq "na");
    
    # iterate haplotigs
    for my $haplotig (sort { $alignments{$primary}{$a}{C} <=> $alignments{$primary}{$b}{C} } keys %{$alignments{$primary}}){
        
        # new contig name
        my $htig_out = $haplotig;
        if (!($dont_rename)){
            $htig_out =~ s/\|.+//;
            $htig_out = $primary . ".H-" . sprintf("%03d", $count) . " " .$htig_out . " " . $alignments{$primary}{$haplotig}{C};
            $count++;
        }
        
        # grab the sequence, remove whitespace
        my $seq = `samtools faidx $haplotigs \"$haplotig\"`;
        $seq =~ s/>.+\n//;
        ($seq) or err("Failed to slup sequence for $haplotig via samtools faidx");
        $seq =~ s/\s//g;
        if ($alignments{$primary}{$haplotig}{D} eq "-"){
            $seq = scalar reverse $seq;
            $seq =~ tr/ACGTacgtN/TGCAtcgaN/;
        }
        
        # print the seq
        if ($scaffold) {
            if ($scaffold_seq) {
                $scaffold_seq .= ("N" x 100);
                $bed_position += 100;
            }
            print $BED "$primary\t$bed_position\t" . ($bed_position + length($seq)) . "\t$haplotig\t" . length($seq) . "\t$alignments{$primary}{$haplotig}{D}\n";
            $bed_position += length($seq);
            $scaffold_seq .= $seq;
        } else {
            print $OFH ">$htig_out\n";
            print $OFH (print_seq($seq));
        }
    }
    
    # print the scaffold seq
    if (($scaffold) && ($scaffold_seq)){
        print $OFH ">$primary\n";
        print $OFH (print_seq($scaffold_seq));
    }
}

# tack on unaligned seqs
if ($tac_contigs){
    for my $primary (keys(%alignments)){
        if ($primary eq "na"){
            for my $haplotig (keys(%{$alignments{$primary}})){
                print $OFH `samtools faidx $haplotigs \"$haplotig\"`;
            }
        }
    }
}


close $OFH;
close $BED if ($scaffold);
msg("Finished!");
exit(0);


sub print_seq {
    my $seq = shift;
    my $out;
    for (my $i=0; $i < length($seq); $i += 60){
        $out .= substr($seq, $i, (($i+60)>length($seq) ? length($seq) - $i : 60) );
        $out .= "\n";
    }
    return $out;
}
```
