## Additional File 1 for "New genome assemblies reveal patterns of domestication and adaptation across *Brettanomyces* (*Dekkera*) species": pull_and_align_SCOs.pl

```
use strict;
use warnings;
use Data::Dumper;
use threads;
use Thread::Semaphore;
use Thread::Queue;

# one-off script to do transcript alignments for all single-copy orthologs
# assumes csv column headers match file names in PWD
my $usage = "
perl pull_and_align_SCOs.pl  Orthogroups.csv  outDir/
";

my $orthogroups = shift or die $usage;
my $outDir = shift or die $usage;


if (!(-d $outDir)){
    mkdir $outDir;
}


# VARS
my $threads = 16;
my %OG;


# THREADS
my $available_threads = Thread::Semaphore->new($threads);
my $writing_to_out = Thread::Semaphore->new(1);
my $queue = Thread::Queue->new();


# parse orthogroups
parseOrtho();

# enqueue jobs
enqueueJobs();

# spawn workers
for (1..$threads){
    $available_threads->down(1);
    threads->create(\&align_job);
}

# wait on jobs
$available_threads->down($threads);
$available_threads->up($threads);
for my $thr (threads->list()){
    $thr->join();
}

# exit
exit(0);


sub parseOrtho {
    open my $TMP, '<', $orthogroups or die;
    my $h = <$TMP>;
    my @head = split /\s+/,$h;
    
    while(<$TMP>){
        my@l=split/\s+/;
        for my $i (1..$#head){
            $OG{$l[0]}{$head[$i]}=$l[$i];
        }
    }
}


sub enqueueJobs {
    # queue jobs
    for my $og (keys %OG){
        $queue->enqueue($og);
    }
    # finalise the queue
    $queue->end();
}


sub align_job {
    while (defined(my $og = $queue->dequeue())) {
        
        # create the multi sequence prot file for aligning
        my $msf = "$outDir/$og.p.fa";
        (-e $msf) and (unlink $msf);
        for my $s (keys %{$OG{$og}}){
            system("samtools faidx $s.faa $OG{$og}{$s} | sed 's/>.*/>$s/' >> $msf")==0 or die;
        }
        
        # create the multi sequence transcript file for later
        my $tmsf = "$outDir/$og.t.fa";
        (-e $tmsf) and (unlink $tmsf);
        for my $s (keys %{$OG{$og}}){
            system("samtools faidx $s.transcripts.fasta $OG{$og}{$s} | sed 's/>.*/>$s/' >> $tmsf")==0 or die;
        }
        
        # run the alignment
        system("muscle -in $msf -out $outDir/$og.MSA.fa")==0 or die;
        
    }
    
    # exit thread
    $available_threads->up(1);

    return;
}
```
