## Additional File 1 for "New genome assemblies reveal patterns of domestication and adaptation across *Brettanomyces* (*Dekkera*) species": separate_reads.pl

```
#!/usr/bin/env perl

use strict;
use warnings;


my $usage = "
Usage:
separate_reads.pl <fastq.gz> <H1.list.gz> <H2.list.gz>

outputs:
H1.list.fastq.gz
H2.list.fastq.gz
";


my $fastq = shift or die $usage;
my $H1 = shift or die $usage;
my $H2 = shift or die $usage;

my $O1 = $H1;
my $O2 = $H2;

($_=~s/gz/fastq.gz/)for($O1, $O2);

my %H1;
my %H2;

open my $FH1, "zcat $H1 |" or die;
while(<$FH1>){
    chomp;
    $H1{$_}=1;
}
close $FH1;

open my $FH2, "zcat $H2 |" or die;
while(<$FH2>){
    chomp;
    $H2{$_}=1;
}
close $FH2;

open my $FQ, "zcat $fastq |" or die;
open my $FO1, "| gzip - > $O1" or die;
open my $FO2, "| gzip - > $O2" or die;

my $c=0;
while(my $h=<$FQ>){
    next if ($h =~ /^\s+$/);
    my $s = <$FQ>;
    my $j = <$FQ>;
    my $q = <$FQ>;
    (length($s)==length($q)) or die;
    my@l=split/\s+/,$h;
    $l[0] =~ s/^@//;
    if ($H1{$l[0]}){
        print $FO1 "$h$s$j$q";
    } 
    if ($H2{$l[0]}){
        print $FO2 "$h$s$j$q";
    }
    if (!($H1{$l[0]})&&!($H2{$l[0]})) {
        print $FO1 "$h$s$j$q";
        print $FO2 "$h$s$j$q";
        $c++;
    }
}

print STDERR "$c unphased reads\n";
```
