## Additional File 1 for "New genome assemblies reveal patterns of domestication and adaptation across *Brettanomyces* (*Dekkera*) species": Snakemake(anomalus).py

```
#!/usr/bin/env snakemake


### CONFIGURATION ###


# give the genome a name
NAME = "B.anomala"

# genome size
GENSIZE = "13m"

# relative path to fast5 directory
FAST5 = "fast5"

# albacore settings
FLOWCELL = "FLO-MIN106"
KIT = "SQK-LSK108"

# stderr directory
ERR = "stderr"


### PIPELINE ###


rule all:
    input:
        NAME + '.fa',
        "cleanup.done"
        

rule cleanup:
    input:
        "gzip_fastq.done",
        "nanopolish.tar.gz"
    output:
        "cleanup.done"
    shell:
        "touch {output}"


rule albacore_basecall:
    input:
        dir=FAST5
    output:
        "albacore/sequencing_summary.txt",
        dir="albacore"
    threads: 16
    log:
        ERR + "/read_fast5_basecaller.stderr"
    shell:
        "read_fast5_basecaller.py --flowcell {FLOWCELL} --kit {KIT} "
        "--recursive --input {input.dir} --worker_threads {threads} --save_path {output.dir} 2> {log}"


rule merge_fastq_reads:
    input:
        "albacore/sequencing_summary.txt"
    output:
        NAME + ".reads.fastq.gz"
    log:
        ERR + "/concatenate_fastq.stderr"
    shell:
        "(cat albacore/workspace/pass/* | gzip - > {output}) 2> {log}"


rule gzip_fastq:
    input:
        NAME + ".reads.fastq.gz"
    output:
        "gzip_fastq.done"
    shell:
        "for i in $(find albacore/ -name \"*.fastq\"); "
            "do gzip $i; "
        "done; "
        "touch {output}"


rule canu_assembly:
    input:
        reads=NAME + ".reads.fastq.gz"
    output:
        gen="canu/" + NAME + ".contigs.fasta",
        dir="canu"
    log:
        ERR + "/canu_assembly.stderr"
    shell:
        "canu -p {NAME} -d {output.dir} -genomeSize={GENSIZE} -correctedErrorRate=0.1 -nanopore-raw {input.reads} 2> {log}"


rule nanopolish_index:
    input:
        summary="albacore/sequencing_summary.txt",
        reads=NAME + ".reads.fastq.gz",
    output:
        expand(NAME + ".reads.fastq.gz.{suffix}", suffix=["index","index.fai","index.gzi","index.readdb"]),
        done="nanopolish_index.done"
    log:
        ERR + "/nanopolish_index.stderr"
    shell:
        "nanopolish index -d {FAST5} -s {input.summary} {input.reads} 2> {log}; "
        "touch {output.done}"


rule BWA_index:
    input:
        "canu/" + NAME + ".contigs.fasta"
    output:
        expand("canu/" + NAME + ".contigs.fasta{suffix}", suffix=[".amb",".ann",".bwt",".pac",".sa"]),
    log:
        ERR + "/bwa_index.stderr"
    shell:
        "bwa index {input} 2> {log}"


rule BWA_mapping:
    input:
        expand("canu/" + NAME + ".contigs.fasta{suffix}", suffix=[".amb",".ann",".bwt",".pac",".sa"]),
        reads=NAME + ".reads.fastq.gz",
        gen="canu/" + NAME + ".contigs.fasta"
    output:
        NAME + ".reads.s.bam"
    threads: 8
    log:
        ERR + "/bwa_mapping.stderr"
    shell:
        "(bwa mem -x ont2d -t {threads} {input.gen} {input.reads} | samtools sort -o {output} -T {NAME}.reads.tmp -) 2> {log}; "
        "samtools index {output}"


rule nanopolish_polish:
    input:
        "nanopolish_index.done",
        gen="canu/" + NAME + ".contigs.fasta",
        reads=NAME + ".reads.fastq.gz",
        bam=NAME + ".reads.s.bam"
    output:
        done="nanopolish/nanopolish.done",
        dir="nanopolish"
    threads: 4
    log:
        ERR + "/nanopolish_polish.stderr"
    shell:
        "( python /home/mike/nanopolish/scripts/nanopolish_makerange.py {input.gen} "
        "| parallel --results nanopolish -P {threads} nanopolish variants --consensus nanopolish/polished.{{1}}.fa "
        "-w {{1}} -r {input.reads} -b {input.bam} -g {input.gen} -t 4 --min-candidate-frequency 0.1 ) 2> {log}; "
        "touch {output.done}"


rule nanopolish_merge:
    input:
        "nanopolish/nanopolish.done",
        dir="nanopolish"
    output:
        NAME + '.merged.fa'
    log:
        ERR + "/nanopolish_merge.stderr"
    shell:
        "python /home/mike/nanopolish/scripts/nanopolish_merge.py {input.dir}/polished.*.fa > {output}"


rule nanopolish_fasta_fix:
    input:
        NAME + '.merged.fa'
    output:
        NAME + '.fa'
    shell:
        "[[ -s {input}.fai ]] || samtools faidx {input}; "
        "for i in `cat {input}.fai | awk '{{print $1}}'`; do "
        "samtools faidx {input} $i >> {output}; done"


rule archive_nanopolish:
    input:
        NAME + '.merged.fa',
        dir="nanopolish"
    output:
        "nanopolish.tar.gz"
    shell:
        "tar czf {output} {input.dir}; "
        "rm -rf {input.dir}"
        
```
