## Additional File 1 for "New genome assemblies reveal patterns of domestication and adaptation across *Brettanomyces* (*Dekkera*) species": species-ratio-OGs.pl

```
use strict;
use warnings;
use Data::Dumper;

# one-off script to identify the orthogroups specific to one of the selected species in an orthofinder results CSV file

my $usage = "
perl species-ratio-OGs.pl  Orthogroups.GeneCount.csv  outDir  sp1  sp2  sp3 

Orthogroups.csv     Orthofinder genecount file
outDir              ouput directory
sp1,2,3             names of the species in the CSV header
";

# input args
my $file_og = shift or die $usage;
my $out_dir = shift or die $usage;

my @sp = @ARGV;
my %sp;             # 0-indexed index for reading CSV file
my %out;            # $out{species}{OG} = (<INT>)

# cutoffs
my $high_cutoff = 6;    # ignore OGs with > this many entries


# read CSV header
open my $FOG, '<', $file_og or die;

{
    chomp(my $h = <$FOG>);
    my @l = split/\t/,$h;
    my $i=0;
    
    # save the indexes to hash
    SP: for my $s (@l){
        $i++;
        for my $q (@sp){
            if ($s eq $q){
                $sp{$s}=$i-1;
                next SP;
            }
        }
    }
    
    # check for missing input sp
    for my $s (@sp){
        ($sp{$s}) or die "$s not in $file_og header";
    }
}

# create the output directory
(-d $out_dir) or mkdir $out_dir;

# parse the CSV
L: while(<$FOG>){
    chomp;
    my@l=split/\t/;
    
    # save the stdev
    my @all;
    for my $s (@sp){
        push @all, $l[$sp{$s}];
    }
    $out{STD}{$l[0]}=stdev(\@all);
    
    # check each species
    SP: for my $s (@sp){
        next L if ($l[$sp{$s}] > $high_cutoff);
        if ($l[$sp{$s}] > 1){
            
            # get the average of the other species
            my $a;
            Q: for my $q (@sp){
               push @all, $l[$sp{$q}];
               next Q if ($s eq $q);
               next L if ($l[$sp{$q}] > $high_cutoff);
               $a+=$l[$sp{$q}]; 
            }
            $a=$a/(scalar @sp - 1);
            
            # prevent divide by zero error
            $a > 0 or $a = 1;
            
            # save the ratio
            $out{$s}{$l[0]}=$l[$sp{$s}]/$a;
        }
    }
}
close $FOG;

# write the output
for my $s (@sp){
    open my $Tbl, '>', "$out_dir/$s.ratio.tsv" or die;
    for my $k (sort {$out{$s}{$b} <=> $out{$s}{$a} }keys (%{$out{$s}})){
        print $Tbl "$k\t$out{$s}{$k}\n";
    }
    close $Tbl;
}
open my $STD, '>', "$out_dir/highStdv.tsv" or die;
for my $k (sort { $out{STD}{$b} <=> $out{STD}{$a} } keys %{$out{STD}}){
    if ($out{STD}{$k} > 2){
        print $STD "$k\t$out{STD}{$k}\n";
    }
}
close $STD;


sub average {
    my($data) = $_[0];
    if (not @{$data}) {
        die("Empty array");
    }
    my $total = 0;
    for (@{$data}) {
        $total += $_;
    }
    my $average = $total / @{$data};
    return $average;
}

sub stdev {
    my($data) = $_[0];
    if(@{$data} == 1){
        return 0;
    }
    my $average = average($data);
    my $sqtotal = 0;
    for (@{$data}) {
        $sqtotal += ($average-$_) ** 2;
    }
    my $std = ($sqtotal / (@{$data}-1)) ** 0.5;
    return $std;
}
```
