## Additional File 1 for "New genome assemblies reveal patterns of domestication and adaptation across *Brettanomyces* (*Dekkera*) species": workflows.pdf

Colour key:

```
# Comment
command
custom_script
```

### *Brettanomyces anomalus* genome assembly

#### Haploid assembly

```
# Link the nanopore and illumina seq
ln -s ../../datasets/generated/anomala-NP/fast5/
ln -s ../../datasets/imported/anomala-PE/ illumina

# DE NOVO ASSEMBLY WITH CANU, CLEANUP WITH PURGE HAPLOTIGS
# Snakemake script will do albacore base-calling, canu assembly
# and nanopolishing.
snakemake --cores 16 -s Snakemake.py

# Cleanup and purge haplotigs
mkdir PH && cd PH
ln -s ../B.anomala.fa

# Align nanopore reads for Purge Haplotigs
bwa index B.anomala.fa
bwa mem -t 8 B.anomala.fa ../B.anomala.reads.fastq.gz \
    | samtools sort -o aligned.s.bam -T ali.tmp -

# Run Purge Haplotigs
purge_haplotigs readhist aligned.s.bam
      # low cutoff = 10, midpoint = 45, high = 120
purge_haplotigs contigcov -in aligned.s.bam.genecov -l 10 -m 45 -h 120
purge_haplotigs purge -g B.anomala.fa -c coverage_stats.csv \
    -b aligned.s.bam -t 16 -windowmasker

cd ../

# Link in main directory
ln -s PH/curated.fasta B.anomala.purged.fasta

# FURTHER BASECALL POLISHING WITH PILON
mkdir pilon && cd pilon/

# Concatenate contigs primary contigs and haploitgs for polishing.
# Re-separate after
cat ../B.anomala.purged.* > gen.fa
bwa index gen.fa
bowtie2-build gen.fa gen.fa

# Map the PE reads
bwa mem -t 12 gen.fa ../illumina/AWRI-953-TS_S3_L001_R1_001.fastq.gz \
    ../illumina/AWRI-953-TS_S3_L001_R2_001.fastq.gz \
    | samtools sort -o PE.s.bam -T ali.tmp -

# Map the MP reads
bowtie2 --fr -I 2000 -X 4000 -p 12 -x gen.fa \
    -1 ../illumina/AWRI-953-MP2-4kb_S7_L001.alltrimmed.1.repaired.fastq.gz \
    -2 ../illumina/AWRI-953-MP2-4kb_S7_L001.alltrimmed.2.repaired.fastq.gz \
    | samtools sort -o MP2.s.bam -T ali.bam -

bowtie2 --fr -I 6000 -X 10000 -p 16 -x gen.fa \
    -1 ../illumina/AWRI-953-MP6-10kb_S11_L001.alltrimmed.1.repaired.fastq.gz \
    -2 ../illumina/AWRI-953-MP6-10kb_S11_L001.alltrimmed.2.repaired.fastq.gz \
    | samtools sort -o MP6.s.bam -T ali.tmp -
```

```
# Merge
samtools merge SR.s.bam PE.s.bam MP2.s.bam MP6.s.bam

# Run pilon
java -jar /home/mike/pilon-1.22.jar --genome gen.fa --frags SR.s.bam \
    --threads 8
cd ..

# Re-separated P and H contigs
samtools faidx B.anomala.purged.fasta
samtools faidx B.anomala.purged.haplotigs.fasta

pil="_pilon"
for i in `cat B.anomala.purged.fasta.fai | awk '{print $1}'`; do \
    samtools faidx pilon/pilon.fasta $i$pil; done \
    > B.anomala.purge.pilon.fasta

for i in `cat B.anomala.purged.haplotigs.fasta.fai | awk '{print $1}'`; do \
    samtools faidx pilon/pilon.fasta $i$pil; done \
    > B.anomala.purge.haplotigs.pilon.fasta
```

#### Diploid assembly (continued from Haploid assembly)

```
# PHASE GENOME WITH HAPCUT2
# Map PE reads for variant calls and longreads for hapcut phasing
mkdir hapcut2
cd hapcut2
ln -s ../B.anomala.purge.pilon.fasta
bwa index B.anomala.purge.pilon.fasta

# Map short reads to make the variant calls
bwa mem B.anomala.purge.pilon.fasta \
    ../illumina/AWRI-953-TS_S3_L001_R1_001.fastq.gz \
    ../illumina/AWRI-953-TS_S3_L001_R2_001.fastq.gz \
    | samtools sort -o PE.s.bam -T ali.tmp -

# Do the variant calling, filter for HET-SNPs on-the-fly
samtools mpileup -f B.anomala.purge.pilon.fasta PE.s.bam --min-MQ 15 \
    | java -jar /home/mike/VarScan.v2.3.9.jar mpileup2snp \
    --min-coverage 20 --min-var-freq 0.2 --min-freq-for-hom 0.8 \
    --p-value 1e-5 --output-vcf grep -P "^#|HET=1" \
    > B.anom.het-SNPs.vcf

# Map the longreads
bwa mem -t 12 -x ont2d B.anomala.purge.pilon.fasta \
    ../B.anomala.reads.fastq.gz \
    | samtools sort -o ON.s.bam -T ali.tmp -
samtools index ON.s.bam

# Split into individual contigs for running hapcut2 (not necessary but
# easier to fix any errors)
mkdir contigs
for i in `cat B.anomala.purge.pilon.fasta.fai | awk '{print$1}'`; do \
    samtools faidx B.anomala.purge.pilon.fasta $i > contigs/$i.fasta; \
    samtools faidx contigs/$i.fasta; \
    done

mkdir bams
for i in `cat B.anomala.purge.pilon.fasta.fai | awk '{print$1}'`; do \
    samtools view -h ON.s.bam $i | samtools view -bS - > bams/$i.bam; \
    samtools index bams/$i.bam; \
    done
```

```
mkdir vcfs
for i in `cat B.anomala.purge.pilon.fasta.fai | awk '{print$1}'`; do \
    grep -P "^#|$i" B.anom.het-SNPs.vcf > vcfs/$i.vcf; \
    done

# Convert hapcut2's output to VCFs for consensus calling
for i in `cat B.anomala.purge.pilon.fasta.fai | awk '{print$1}'`; do \
     whatshap hapcut2vcf -o vcfs/$i.hap.vcf vcfs/$i.vcf haps/$i.hap; \
     done
for i in `ls vcf | grep hap`; do bgzip $i; tabix $i.gz; done

# Write new consensus seqs for each haplotype, these will only have the
# HET-SNPs in the VCF phased, not the SVs, InDels etc.
for i in `cat B.anomala.purge.pilon.fasta.fai | awk '{print$1}'`; do \
    bcftools consensus -f contigs/$i.fasta -H 1 vcfs/$i.hap.vcf.gz \
        > contigs/$i.H1.fasta; \
    bcftools consensus -f contigs/$i.fasta -H 2 vcfs/$i.hap.vcf.gz \
        > contigs/$i.H2.fasta; \
done

# Concatenate the phased contigs and add labels to the contig names
cat contigs/*.H1.fasta | sed 's/pilon/H1/' > phased.contigs.fasta
cat contigs/*.H2.fasta | sed 's/pilon/H2/' >> phased.contigs.fasta

# READ BINNING FOR REASSEMBLY OF SEPARATED HAPLOTYPES
# Map the long reads again
bwa index phased.contigs.fasta
bwa mem -t 12 -x ont2d phased.contigs.fasta ../B.anomala.reads.fastq.gz \
    | samtools sort -o phased.bam -T ali.tmp -

# Get the separated read IDs for the two haplotypes
for i in `cat phased.contigs.fasta.fai | awk '{print$1}' | grep H1`; do \
    samtools view phased.bam $i | awk '{print$1}'; \
    done | gzip - > H1.reads.gz

for i in `cat phased.contigs.fasta.fai | awk '{print$1}' | grep H2`; do \
    samtools view phased.bam $i | awk '{print$1}'; \
    done | gzip - > H2.reads.gz

# Separate the FASTQ reads by read IDs
perl separate_reads.pl ../B.anomala.reads.fastq.gz H1.reads.gz H2.reads.gz

# REASSEMBLY OF SEPARATED HAPLOTYPES WITH CANU
mkdir reassembly

for i in H1 H2; do
    mkdir reassembly/$i && cd reassembly/$i
    ln -s ../../$i.reads.fastq.gz
    canu -p B.anomala.$i -d $i -genomeSize=14000000 -correctedErrorRate=0.1 \
        -nanopore-raw $i.reads.fastq.gz 2> canu.stderr

    # cleanup the reassembly
    bwa index $i/B.anomala.$i.contigs.fasta
    bwa mem -t 12 -x ont2d $i/B.anomala.$i.contigs.fasta $i.reads.fastq.gz \
        | samtools sort -o $i.bam -T ali.tmp -
    purge_haplotigs readhist $i.bam
    purge_haplotigs contigcov -i $i.bam.genecov -l 40 -m 110 -h 150
    purge_haplotigs purge -g $i/B.anomala.$i.contigs.fasta \
        -c coverage_stats.csv -b $i.bam -t 12 -o $i.curated
    cd ../
done
```

```
# Concatenate for polishing, add haplotype suffixes
samtools faidx H1/H1.curated.fasta
samtools faidx H2/H2.curated.fasta
for i in `cat H1/H1.curated.fasta.fai | awk '$2>=10000{print $1}'`; do \
    samtools faidx H1/H1.curated.fasta $i | sed 's/>\(tig[0-9]*\)/>\1.H1/'; \
    done > H1_H2.fasta
for i in `cat H2/H2.curated.fasta.fai | awk '$2>=10000{print $1}'`; do \
    samtools faidx H2/H2.curated.fasta $i | sed 's/>\(tig[0-9]*\)/>\1.H2/'; \
    done >> H1_H2.fasta

# POLISH WITH NANOPOLISH AND PILON
bwa index H1_H2.fasta
bwa mem -x ont2d -t 12 H1_H2.fasta ../../B.anomala.reads.fastq.gz \
    | samtools sort -o H1_H2.bam -T aln.tmp -

# Link the fast5 directory
ln -s ../../fast5/ .

# Nanopolish
python ~/nanopolish/scripts/nanopolish_makerange.py H1_H2.fasta \
    | parallel --results nanopolish.results -P 4 \
        nanopolish variants --consensus \
        polished{1}.fa -w {1} -r ../../B.anomala.reads.fastq.gz \
        -b H1_H2.bam -g H1_H2.fasta \
        -t 4 --min-candidate-frequency 0.1

# Merge, use samtools to fix FASTA format, cleanup
python ~/nanopolish/scripts/nanopolish_merge.py polished.*.fa > pol.fa
samtools faidx pol.fa
for i in `cat pol.fa.fai | awk '{print $1}'`; do \
    samtools faidx pol.fa $i; \
done > H1_H2.nanopol.fa
rm pol.fa* polished.*.fa

# Pilon
# Map PE reads
bwa index H1_H2.nanopol.fa
bwa mem -t 12 H1_H2.nanopol.fa \
    ../../illumina/AWRI-953-TS_S3_L001_R1_001.fastq.gz \
    ../../illumina/AWRI-953-TS_S3_L001_R2_001.fastq.gz \
    | samtools sort -o H1_H2.nanpol.bam -T aln.tmp -
samtools index H1_H2.nanpol.bam
samtools flagstat H1_H2.nanpol.bam

# Run pilon
java -jar ~/pilon-1.22.jar --genome H1_H2.nanopol.fa \
    --frags H1_H2.nanpol.bam --threads 16

# Re-separate the polished contigs
samtools faidx pilon.fasta
for i in `cat pilon.fasta.fai | grep H1 | awk '{print $1}'`; do \
    samtools faidx pilon.fasta $i; \
    done > B.anomala.H1.fasta

for i in `cat pilon.fasta.fai | grep H2 | awk '{print $1}'`; do \
    samtools faidx pilon.fasta $i; \
    done > B.anomala.H2.fasta
```

#### *Brettanomyces custersianus* genome assembly

```
# Link NP dataset
ln -s ../../datasets/generated/custersianus-NP/fast5/

# Basecalling with albacore
read_fast5_basecaller.py --flowcell FLO-MIN107 --kit SQK-LSK108 \
    --recursive --input fast5/ \
    --worker_threads 24 --save_path ./albacore

# Concatenate the fastq files
cat albacore/workspace/pass/* | gzip - > reads.fastq.gz

# DE NOVO ASSEMBLY WITH CANU
mkdir CANU && cd CANU

# Run Canu
canu -p cus -d cus-minion -genomeSize=13m -correctedErrorRate=0.1 \
    -nanopore-raw ../reads.fastq.gz

# BASECALL POLISHING WITH NANOPOLISH
# Link and index canu assembly for mapping
mkdir BWA && cd BWA/
ln -s ../CANU/cus-minion/cus.contigs.fasta
bwa index cus.contigs.fasta
cd ../

# Index reads for Nanopolish
nanopolish index -d albacore -s albacore/sequencing_summary.txt reads.fastq.gz

# Map reads for Nanopolish
bwa mem -x ont2d -t 16 BWA/cus.contigs.fasta reads.fastq.gz \
    | samtools sort -o reads.s.bam -T reads.tmp -
samtools index reads.s.bam

# Run Nanopolish
python ~/nanopolish/scripts/nanopolish_makerange.py BWA/cus.contigs.fasta \
    | parallel --results nanopolish.results -P 4 \
        nanopolish variants --consensus polished.{1}.fa \
        -w {1} -r reads.fastq.gz -b reads.s.bam -g BWA/cus.contigs.fasta \
        -t 4 --min-candidate-frequency 0.1

# Merge
python ~/nanopolish/scripts/nanopolish_merge.py polished.*.fa \
    > B.custersianus.fasta

# Cleanup
mv polished.*.fa nanopolish.results
gzip nanopolish.results/*.fa

# Fix the fasta line lengths (nanoploish makes each fasta seq 1 single line)
samtools faidx B.custersianus.fasta
for i in `cat B.custersianus.fasta.fai \
    | awk '{print $1}'`; do \
        samtools faidx B.custersianus.fasta $i >> tmp; \
    done
mv tmp B.custersianus.fasta

# Cleanup
rm B.custersianus.fasta.fai
samtools faidx B.custersianus.fasta
```

```
# BASECALL POLISHING WITH PILON
# Link the illumina seq data
ln -s ../../datasets/imported/custersianus-PE/ illumina

# Index for mapping
ln -s -r B.custersianus.fasta BWA/
bwa index BWA/B.custersianus.fasta

mkdir BOWTIE
ln -sr B.custersianus.fasta BOWTIE/
bowtie2-build BOWTIE/B.custersianus.fasta BOWTIE/B.custersianus.fasta

# Map PE shortreads
bwa mem -t 16 BWA/B.custersianus.fasta \
    illumina/AWRI-951-TS_S2_L001_R1_001.short.fastq.gz \
    illumina/AWRI-951-TS_S2_L001_R2_001.short.fastq.gz \
    | samtools sort -o PE.s.bam -T PE.tmp -

# Map MP 2-4kb
bowtie2 --fr -I 2000 -X 4000 -p 16 -x BOWTIE/B.custersianus.fasta \
    -1 illumina/AWRI-951-MP2-4kb_S6_L001.alltrimmed.1.repaired.fastq.gz \
    -2 illumina/AWRI-951-MP2-4kb_S6_L001.alltrimmed.2.repaired.fastq.gz \
    | samtools sort -o MP2-4-reads.s.bam -T MP2-4.tmp -

# Map MP 6-10kb
bowtie2 --fr -I 6000 -X 10000 -p 16 -x BOWTIE/B.custersianus.fasta \
    -1 illumina/AWRI-951-MP6-10kb_S10_L001.alltrimmed.1.repaired.fastq.gz \
    -2 illumina/AWRI-951-MP6-10kb_S10_L001.alltrimmed.2.repaired.fastq.gz \
    | samtools sort -o MP6-10-reads.s.bam -T MP6-10.tmp -

# Merge bam files
samtools merge SR.s.bam PE.s.bam MP2-4-reads.s.bam MP6-10-reads.s.bam

# Run Pilon
java -jar ~/pilon-1.22.jar --genome B.custersianus.fasta \
    --frags SR.s.bam

# Rename
mv pilon.fasta B.custersianus.pilon.fasta

# CLEANUP WITH PURGE HAPLOTIGS
# Link and index assembly for mapping
cd BWA/
ln -s ../B.custersianus.pilon.fasta
bwa index B.custersianus.pilon.fasta
cd ../

# Purge haplotigs
mkdir purge_haplotigs && cd purge_haplotigs/

# Link assembly
ln -s ../B.custersianus.pilon.fasta

# Map longreads
bwa mem -x ont2d -t 16 BWA/B.custersianus.pilon.fasta reads.fastq.gz \
    | samtools sort -o reads.pol.s.bam -T reads.tmp -
samtools index reads.pol.s.bam
```

```
# Run Purge Haplotigs
purge_haplotigs readhist reads.pol.s.bam
      # low cutoff = 10, midpoint = 45, high = 120
purge_haplotigs contigcov -in reads.pol.s.bam.genecov -l 20 -m 110 -h 190
purge_haplotigs purge -g B.custersianus.pilon.fasta -c coverage_stats.csv \
    -b aligned.s.bam -t 16 -windowmasker
```

#### *Brettanomyces naardenensis* genome assembly

```
# Link the dataset
ln -s ../../datasets/generated/naardenensis-NP/fast5/

# ASSEMBLY WITH CANU, POLISH WITH NANOPOLISH
# Run the snakemake pipeline (Canu assembly and Nanopolish)
snakemake -s Snakemake.py --cores 32

# Map the short reads for pilon polishing
ln -s ../../datasets/imported/naardenensis-PE/ illumina

# BASECALL POLISH WITH PILON
# Prep for mapping
mkdir BWA && cd BWA/
ln -s ../B.naardenensis.merged.fa
bwa index B.naardenensis.merged.fa
cd ../
mkdir BOWTIE && cd BOWTIE/
ln -s ../B.naardenensis.merged.fa
bowtie2-build B.naardenensis.merged.fa B.naardenensis.merged.fa
cd ../

# Map the PE reads
bwa mem -t 16 ../BWA/B.naardenensis.merged.fa \
    illumina/AWRI-950-TS_S1_L001_R1_001.fastq.gz \
    illumina/AWRI-950-TS_S1_L001_R2_001.fastq.gz \
    | samtools sort -@ 8 -m 2G -o PE.s.bam -T PE.tmp -

# Map the MP reads 1
bowtie2 --fr -I 2000 -X 4000 -p 16 -x ../BOWTIE/B.naardenensis.merged.fa \
    -1 illumina/AWRI-950-MP2-4kb_S5_L001.alltrimmed.1.repaired.fastq.gz \
    -2 illumina/AWRI-950-MP2-4kb_S5_L001.alltrimmed.2.repaired.fastq.gz \
    | samtools sort -@ 8 -m 1G -o MP2.s.bam -T MP2.tmp -

# Map the MP reads 2
bowtie2 --fr -I 6000 -X 10000 -p 16 -x ../BOWTIE/B.naardenensis.merged.fa \
    -1 illumina/AWRI-950-MP6-10kb_S9_L001.alltrimmed.1.repaired.fastq.gz \
    -2 illumina/AWRI-950-MP6-10kb_S9_L001.alltrimmed.2.repaired.fastq.gz \
    | samtools sort -@ 8 -m 2G -o MP6.s.bam -T MP6.tmp -

# Merge the BAM files
samtools merge SR.s.bam PE.s.bam MP2.s.bam MP6.s.bam
samtools index SR.s.bam

# Run pilon
java -jar ~/pilon-1.22.jar --genome B.naardenensis.merged.fa \
    --frags illumina/SR.s.bam --threads 8

# Rename
mv pilon.fasta B.naardenensis.pilon.fa
```

```
# CLEANUP WITH PURGE HAPLOTIGS
# Index the pilon-polished assembly and re-map the longreads
cd BWA
ln -s ../B.naardenensis.pilon.fa .
bwa index B.naardenensis.pilon.fa
cd ..
bwa mem -t 16 BWA/B.naardenensis.pilon.fa B.naardenensis.reads.fastq.gz \
    | samtools sort -@ 4 -m 2G -o reads.pol.s.bam -T reads.pol.tmp -

# Run Purge Haplotigs
# Read-depth histogram
purge_haplotigs readhist reads.pol.s.bam

# Apply coverage cutoffs
purge_haplotigs contigcov -l 12 -m 12 -h 50 -i reads.pol.s.bam.genecov

# Purge step
purge_haplotigs purge -g B.naardenensis.pilon.fa -c coverage_stats.csv \
    -b reads.pol.s.bam -o B.naardenensis.curated
```

#### *Brettanomyces nanus* genome assembly

```
# Link the nanopore dataset
ln -s ../../datasets/generated/nanus-NP/fast5/

# ASSEMBLY WITH CANU AND BASECALL POLISHING WITH NANOPOLISH
# Run the pipeline
snakemake -s Snakemake.py --cores 16

# CLEANUP WITH PURGE HAPLOTIGS
mkdir purgehap

# Map reads
minimap2 B.nanus.merged.fa B.nanus.reads.fastq.gz -t 12 -a \
    | samtools view -hF 256 - \
    | samtools sort -@ 4 -m 1G -o purgehap/aligned.bam -t tmp.ali

cd purgehap

# Read-depth histogram
purge_haplotigs readhist -g ../B.nanus.merged.fa -b aligned.bam
    # cutoffs are -l 5 -m 30 -h 190
purge_haplotigs contigcov -i aligned.bam.gencov -l 5 -m 30 -h 190
purge_haplotigs purge -g ../B.nanus.merged.fa -c coverage_stats.csv

# Copy cleaned assemblies
cp curated.fasta ../B.nanus.NP.PH.fasta
cp curated.fasta ../B.nanus.NP.PH.haplotigs.fasta

cd ../

# BASECALL POLISHING WITH PILON
# Link illumina dataset
ln -s ../../datasets/imported/nanus-PE/ illumina

mkdir pilon && cd pilon/
```

```
# Combine and index for mapping
cat ../B.nanus.NP.PH.fasta ../B.nanus.NP.PH.haplotigs.fasta \
    > B.nanus.NP.PH.p-h.fasta
bowtie2-build B.nanus.NP.PH.p-h.fasta B.nanus.NP.PH.p-h.fasta
bwa index B.nanus.NP.PH.p-h.fasta

# Map PE reads
bwa mem -t 12 B.nanus.NP.PH.p-h.fasta \
    ../illumina/AWRI-2847-TS_S4_L001_R1_001.short.fastq.gz \
    ../illumina/AWRI-2847-TS_S4_L001_R2_001.short.fastq.gz \
    | samtools sort -@ 4 -m 1G -o PE.bam -t tmp.ali

# Map MP reads 1
bowtie2 --fr -I 2000 -X 4000 -p 12 -x B.nanus.NP.PH.p-h.fasta \
  -1 ../illumina/AWRI-2847-MP2-4kb_S8_L001.alltrimmed.1.repaired.short.fastq.gz \
  -2 ../illumina/AWRI-2847-MP2-4kb_S8_L001.alltrimmed.2.repaired.short.fastq.gz \
  | samtools sort -@ 4 -m 1G -o MP2.bam -T MP2.tmp -

# Map MP reads 2
bowtie2 --fr -I 6000 -X 10000 -p 12 -x B.nanus.NP.PH.p-h.fasta \
  -1 \
  ../illumina/AWRI-2847-MP6-10kb_S12_L001.alltrimmed.1.repaired.short.fastq.gz \
  -2 \
  ../illumina/AWRI-2847-MP6-10kb_S12_L001.alltrimmed.2.repaired.short.fastq.gz \
  | samtools sort -@ 4 -m 1G -o MP6.bam -T MP6.tmp -

# Merge BAM files
samtools merge SR.bam PE.bam MP2.bam MP6.bam
samtools index SR.bam

# Run pilon
java -jar ~/pilon.jar --genome B.nanus.NP.PH.p-h.fasta \
    --frags SR.bam --threads 12

# Rename and remove ‘_pilon’ suffix
cat pilon.fasta | sed 's/_pilon//' > tmp
mv tmp pilon.fasta

# Split back into primary contigs and haplotigs
for i in `grep \> B.nanus.NP.PH.p-h.fasta \
         | sed 's/>//' \
         | grep -v HAPLOTIG \
         | awk '{print $1}'`; do
    samtools faidx pilon.fasta $i;
done > ../B.nanus.fasta

for i in `grep \> B.nanus.NP.PH.p-h.fasta \
        | sed 's/>//' \
        | grep HAPLOTIG \
        | awk '{print $1}'`; do
    samtools faidx pilon.fasta $i;
done > ../B.nanus.haplotigs.fasta
```

#### Manual inspection and miss-assembly correction

```
# Only B. naardenensis and B. custersianus shown as no miss-assemblies were
identified # for the other assemblies.
```

##### *Brettanomyces naardenensis*

```
mkdir naard && cd naard
ln -s ../../B.naardenensis.fasta

# Scale down the reads (using for visualisation)
zcat ../../naardenensis/illumina/AWRI-950-TS_S1_L001_R1_001.fastq.gz \
    | perl -e '
      my$c==0;while(<>){
          my$s=<>;
          my$j=<>;
          my$q=<>;
          if($c==3){
              print "$_$s$j$q";
              $c=0;
          }else{
              $c++;
          }
      }' \
    | gzip - > R1.fq.gz

zcat ../../naardenensis/illumina/AWRI-950-TS_S1_L001_R2_001.fastq.gz \
    | perl -e '
      my$c==0;while(<>){
          my$s=<>;
          my$j=<>;
          my$q=<>;
          if($c==3){
              print "$_$s$j$q";
              $c=0;
          }else{
              $c++;
          }
      }' \
    | gzip - > R2.fq.gz

zcat ../../naardenensis/B.naardenensis.reads.fastq.gz \
    | perl -e '
      my$c==0;while(<>){
          my$s=<>;
          my$j=<>;
          my$q=<>;
          if($c==1){
              print "$_$s$j$q";
              $c=0;
          }else{
              $c++;
          }
      }' \
    | gzip - > np.fa.gz
```

```
# Map the reads
bwa index B.naardenensis.fasta
bwa mem -t 12 B.naardenensis.fasta R1.fq.gz R2.fq.gz \
    | samtools sort -o PE.bam -T ali.tmp -
samtools index PE.bam

bwa mem -x ont2d -t 12 B.naardenensis.fasta np.fa.gz \
    | samtools sort -o np.bam -T ali.tmp -
samtools index np.bam

# Align assembly to AWRI2804
nucmer -b 500 -c 40 -d 0.5 -g 200 -l 12 -t 4 ../../AWRI2804.fasta
B.naardenensis.fasta
delta-filter -1 out.delta > out.1.delta
mummerplot --fat --png --large out.1.delta

# Manually curate
# (eyeball the bam file, dotplot, show-coords output,
# note misassemblies and split contigs)
    # MISSASSEMBLIES
        # tig00000001_pilon:1723775
        # tig00007031_pilon:772570
        # tig00000003_pilon:509005
        # tig00000003_pilon:638002

# Break the contigs at the misassembly positions
samtools faidx B.naardenensis.fasta tig00000001_pilon:0-1723775 > cur.fa
samtools faidx B.naardenensis.fasta tig00000001_pilon:1723776- >> cur.fa
samtools faidx B.naardenensis.fasta tig00007031_pilon:0-772570 >> cur.fa
samtools faidx B.naardenensis.fasta tig00007031_pilon:772570- >> cur.fa
samtools faidx B.naardenensis.fasta tig00000003_pilon:0-509005 >> cur.fa
samtools faidx B.naardenensis.fasta tig00000003_pilon:509005-638002 >> cur.fa
samtools faidx B.naardenensis.fasta tig00000003_pilon:638002- >> cur.fa

# Add the rest of the contigs
for i in `cat B.naardenensis.fasta.fai \
         | grep -v tig00000001_pilon \
         | grep -v tig00000003_pilon \
         | grep -v tig00007031_pilon \
         | awk '{print $1}'`; do \
    samtools faidx B.naardenensis.fasta $i; \
done >> cur.fa

# Rename
mv cur.fa B.naardenensis.FIXED.fasta

# Recheck with MP reads
bbmap.sh threads=8 out=stdout ref=B.naardenensis.FIXED.fasta \
    in=AWRI-950-MP2-4kb_S5_L001.alltrimmed.1.repaired.fastq.gz \
    in2=AWRI-950-MP2-4kb_S5_L001.alltrimmed.2.repaired.fastq.gz \
    | samtools sort -o naa.2k.bam -T tmp.ali
samtools index naa.2k.bam

bbmap.sh threads=8 out=stdout ref=B.naardenensis.FIXED.fasta \
    in=AWRI-950-MP6-10kb_S9_L001.alltrimmed.1.repaired.fastq.gz \
    in2=AWRI-950-MP6-10kb_S9_L001.alltrimmed.2.repaired.fastq.gz \
    | samtools sort -o naa.6k.bam -T tmp.ali
samtools index naa.6k.bam
```

```
# More manual checking
    # no obvious miss-assemblies remaining

# Copy the new assembly
cp B.naardenensis.FIXED.fasta ../../../outputs/assemblies/B.naardenensis.fasta
```

##### *Brettanomyces custersianus*

```
mkdir cust && cd cust
ln -s ../../B.custersianus.fasta
bwa index B.custersianus.fasta

# Down-sample reads
zcat ../../custersianus/illumina/AWRI-951-TS_S2_L001_R1_001.fastq.gz \
    | perl -e '
        my$c==0;while(<>){
            my$s=<>;
            my$j=<>;
            my$q=<>;
            if($c==3){
                print "$_$s$j$q";
                $c=0;
            }else{
                $c++;
            }
        }' \
    | gzip - > R1.fq.gz

zcat ../../custersianus/illumina/AWRI-951-TS_S2_L001_R2_001.fastq.gz \
    | perl -e '
        my$c==0;while(<>){
            my$s=<>;
            my$j=<>;
            my$q=<>;
            if($c==3){
                print "$_$s$j$q";
                $c=0;
            }else{
                $c++;
            }
        }' \
    | gzip - > R2.fq.gz

zcat ../../custersianus/reads.fastq.gz \
    | perl -e '
        my$c==0;while(<>){
            my$s=<>;
            my$j=<>;
            my$q=<>;
            if($c==1){
                print "$_$s$j$q";
                $c=0;
            }else{
                $c++;
            }
        }' \
    | gzip - > np.fa.gz
```

```
# Map reads
bwa mem -t 12 B.custersianus.fasta R1.fq.gz R2.fq.gz \
    | samtools sort -o PE.bam -T tmp.aln -
samtools index PE.bam

bwa mem -x ont2d -t 12 B.custersianus.fasta np.fa.gz \
    | samtools sort -o NP.bam -T tmp.aln -
samtools index NP.bam

# Align assembly to AWRI2804
nucmer -b 500 -c 40 -d 0.5 -g 200 -l 12 -t 4 ../../AWRI2804.fasta
    B.custersianus.fasta
delta-filter -1 out.delta > out.1.delta
mummerplot --fat --png --large out.1.delta

# Manually curate
# (eyeball the bam file, dotplot, show-coords output,
# note misassemblies and split contigs)
    # MISSASSEMBLIES
        # tig00000008_pilon:834510

# Break at misassembly point
samtools faidx B.custersianus.fasta tig00000008_pilon:0-834510 > cur.fa
samtools faidx B.custersianus.fasta tig00000008_pilon:834510- >> cur.fa

# Re-add the rest of the contigs
for i in `cat B.custersianus.fasta.fai \
         | grep -v tig00000008_pilon \
         | awk '{print $1}'`; do \
    samtools faidx B.custersianus.fasta $i; \
done >> cur.fa

# Rename
mv cur.fa B.custersianus.FIXED.fasta

# Re-check with MP reads
bbmap.sh threads=8 out=stdout ref=B.custersianus.FIXED.fasta \
    in=AWRI-951-MP2-4kb_S6_L001.alltrimmed.1.fastq.gz \
    in2=AWRI-951-MP2-4kb_S6_L001.alltrimmed.2.fastq.gz \
    | samtools sort -o cus.2k.bam -T tmp.ali
samtools index cus.2k.bam

bbmap.sh threads=8 out=stdout ref=B.custersianus.FIXED.fasta \
    in=AWRI-951-MP6-10kb_S10_L001.alltrimmed.1.fastq.gz \
    in2=AWRI-951-MP6-10kb_S10_L001.alltrimmed.2.fastq.gz \
    | samtools sort -o cus.6k.bam -T tmp.ali
samtools index cus.6k.bam

# Manually inspect again
    # no obvious miss-assemblies

# Copy the new assembly
cp B.custersianus.FIXED.fasta ../../../outputs/assemblies/B.custersianus.fasta
```

### Gene prediction and annotation

```
# GENE PREDICTION WITH AUGUSTUS
# example for Brettanomyces anomalus (haploid)
$GENOME = 'B.anomalus'

# run Augustus
augustus --species=saccharomyces_cerevisiae_S288C $GENOME.fasta > $GENOME.gff

# return protein seqs (Augustus adds these as comments in the GFF file)
cat genome.gff \
    | perl -e '
        $/ = "end gene";
        while(<>){
            my$gen;
            if(m/start gene (\w+)/){
                $gen=$1;
            }else{
                die;
            }
            s/#//g;
            if(m/protein sequence = \[([\w\s]+)\]/){
                $s=$1;
                $s=~s/\s//g;
                print ">$gen\n$s\n";
            }
        }'\
    > $GENOME.prot.fasta

# Add protein names using BlastP hits to UniProtKB/Swiss-Prot
# Need a table of seqIDs to prot names from the uniprot DB for later
grep \> ~/blastdb/uniprot_sprot.fasta \
    | sed 's/>//' \
    | sed 's/OS=.*//' \
    | sed 's/ /\t/' \
    > uniprot_sprot.names.tsv

# blastp predicted proteins against uniprotKB
cat $GENOME.prot.fasta \
    | parallel -j 16 --recstart '>' --blocksize 10k --pipe \
        blastp -query - -db ~/blastdb/uniprot_sprot.fasta \
        -evalue 0.00001 -num_alignments 1 -outfmt 6 \
    | gzip - > $GENOME.uniprotkb.outfmt6.gz

# ORTHOGROUP ANNOTATION WITH ORTHOFINDER
mkdir orthofinder && cd orthofinder

# link all the prot FASTA files
for i in genome1 genome2 genome3; do
    ln -s ../$i.prot.fasta;
done

cd ../

# run
orthofinder -t 16 -f orthofinder/
```

```
# ADD THE PROTEIN NAMES AND ORTHOGROUPS TO FASTAs and GFFs
# custom script uses FOFNs
ls | grep gff | grep -v fofn > gff.fofn
ls | grep outfmt6 > blast.fofn
ls | grep prot.fasta > prot.fofn

# run the script to add annotations to protein FASTAs and GFFs
perl add_annotations.pl  prot.fofn  blast.fofn  \
    uniprot_sprot.names.tsv  gff.fofn  \
    orthofinder/Results_Oct18/Orthogroups.csv
```

#### Synteny plots of assemblies

```
# Example for B. bruxellensis synteny with B. naardenensis
# Some contigs were manually re-arranged in AWRI2804 for optimise the layout

# Place the naardenensis contigs to the bruxellensis reference
purge_haplotigs ncbiplace -p AWRI2804.fasta -h B.naardenensis.fasta \
    -t 8 -c 0 -o BvNa.place.tsv

# Re-order according to the placements
perl orient_haplotigs.pl -p AWRI2804.fasta -h B.naardenensis.fasta \
    -n BvNa.place.tsv -o B.naardenensis.orient -r -t

# Tidy up
mv tmp_purge_haplotigs/ tmp_purge_haplotigs_BvNa

# Make windows of the reference genome (AWRI2804)
bedtools makewindows -g AWRI2804.fasta.fai -w 20000 -s 10000 > AWRI2804.windows

# Add the reference contigs to a kar file, this will be used for all
# plots that have bruxellensis as the reference
cat AWRI2804.fasta.fai \
    | awk '{print "chr - "$1" "$1" 0 "$2" white"}' > AWRI2804.kar
# Manually changed the colors of the 2804 contigs to set3-12-qual-1,
# set3-12-qual-2, etc.

# naardenensis windows
bedtools makewindows -g B.naardenensis.orient.fa.fai -w 20000 -s 10000 \
    > B.naardenensis.windows

# Align naardenensis to reference
nucmer -b 500 -c 40 -d 0.5 -g 200 -l 12 -t 16 \
    AWRI2804.fasta B.naardenensis.orient.fa -p BvNo
delta-filter -1 BvNo.delta > BvNo.1.delta
show-coords -TH BvNo.1.delta > BvNo.coords

# Pair reference and query genome windows based on alignments, write
# the translation table.
perl nucmer2trans.pl -r AWRI2804.windows -q B.naardenensis.windows \
    -c BvNo.coords > BvNo.trn.tsv

# Kar file for circos
cat AWRI2804.kar > BvNo.kar

# Add naardenensis contigs to new kar file
tac B.naardenensis.orient.fa.fai \
    | awk '{print "chr - "$1" "$1" 0 "$2" vdgrey" }' \
    | sed 's/:/_/' >> BvNo.kar
```

```
# Convert the translation table to a links file for circos
cat BvNo.trn.tsv \
    | awk '{
        if($7=="+"){
            print $1"\t"$2"\t"$3"\t"$4"\t"$5"\t"$6"\t"$8
        }else{
            print $1"\t"$2"\t"$3"\t"$4"\t"$6"\t"$5"\t"$8
        }' \
    | sed 's/:/_/' \
    | ./map_colors.pl > BvNo.links

# Circos conf file from template
cat circos/circos.conf | sed 's/##XX##/BvNo/' > circos/BvNo.conf

# Run circos
~/circos-0.69-6/bin/circos -conf circos/BvNo.conf -file BvNo
```

#### Gene enrichment analysis

```
# This analysis used the OrthoFinder run using representative species
# from Saccharomycetaceae (check supplementary information for complete
# list of species).

# Run InterProScan to generate GO-terms for GO enrichment analysis
# Example for B. bruxellensis
interproscan.sh -cpu 8 -d bruxellensis -dp -goterms -i bruxellensis.prot.faa

# convert IPS5 output to GO annotation anno input for cytoscape/BiNGO
for i in anomalus anomalus_H1 anomalus_H2 \
        bruxellensis custersianus \
        naardenensis nanus; do
    printf "(species=Brettanomyces $i)(type=ORFs)(curator=GO)\n" \
        > $i/$i.GO.anno
    cat $i/*.faa.tsv | grep GO: | sed 's/\t.\+\tGO:/\t/' \
        | sed 's/[|GO:]\+/,/g' | perl -e '
            while(<>){
                my@l=split/\s+/;
                my@o=split/,/,$l[1];
                for my $o(@o){
                    print "$l[0] = $o\n";
                }
            }
        ' | sort -k1,1 -k2,2 | uniq >> $i/$i.GO.anno
done

# Link the orthofinder results for use here.
ln -s ../functional-annotation/orthofinder/Results_Nov15/Orthogroups.csv
ln -s ../functional-annotation/orthofinder/Results_Nov15/Orthogroups.GeneCount.csv

# Use a custom script to get the ratios of genecounts for each OG, for
# each species, ignoring species entries with gene counts of one, and ignoring
# OG entries with a species count > 6 (to filter out the TE garbage). It will
# also calculate the standard deviations and output any OGs > 2 STDEV (for OGs
# that are enriched in more than one species).
perl species-ratio-OGs.pl Orthogroups.GeneCount.csv species-ratio-OGs \
    Brettanomyces_anomala_h1 Brettanomyces_anomala_h2 \
    Brettanomyces_bruxellensis Brettanomyces_custersianus \
    Brettanomyces_naardenensis Brettanomyces_nanus
```

```
# Get the list of OGs for each species with ratios >= 2. Use the SQL database to
# pull the gene names for these OGs. We then feed this into cytoscape with BiNGO
# to search for enriched GO terms.
cd species-ratio-OGs

for i in bruxellensis anomala_h1 anomala_h2 custersianus naardenensis nanus; do
    cat Brettanomyces_$i.ratio.tsv | awk '$2>=2{print $1}' \
    > Brettanomyces_$i.ratio.OGs.tsv
done

for i in anomala_h1 anomala_h2 bruxellensis custersianus naardenensis nanus; do
    cat highStdv.tsv >> Brettanomyces_$i.ratio.OGs.tsv
done

cp Brettanomyces_*.OGs.tsv ../../functional-annotation/DB

# Obtaining gene IDs for enriched OGs
# Example in sqlite3:

sqlite> create table brettGO (orthogroup TEXT not null);
sqlite> .import Brettanomyces_bruxellensis.ratio.OGs.tsv brettGO
sqlite> select gene from annotations a
    inner join bruxGO b on b.og = a.orthogroup
    where a.species like 'Brettanomyces_bruxellensis';

# BiNGO analysis
# Gene IDs from SQL queries above were pasted into BiNGO. 'Overrepresentation'
# and 'Visualization' were checked. Hypergeometric test, Bonferroni FWER
# correction, 0.9 significant level. Overrepresented categories after correction
# to visualize. Whole annotation used as reference set. Interproscan GO .anno
# files used as the organism/annotation set. Output table and network saved as
# TSV and SVG respectively. TSV had some java code to be cleaned up like so:
for i in anomala bruxellensis custersianus naardenensis nanus; do
    cat $i.ratio.tsv | sed 's/javax.swing.*text=//' \
        | sed 's/,verticalAlignment=CENTER.*]//' > tmp
    mv tmp $i.ratio.tsv
done

# Extract all the unique gene IDs with enriched GO-terms
cat anomala.ratio.tsv | sed 's/^.*%\sH/H/' | sed 's/G/g/g' \
    | sed 's/\s/\n/g' | sort | uniq | grep H1_ | sed 's/H2_//' \
    > anomala_h1.enrichedGeneIDs
cat anomala.ratio.tsv | sed 's/^.*%\sH/H/' | sed 's/G/g/g' \
    | sed 's/\s/\n/g' | sort | uniq | grep H2_ | sed 's/H2_//' \
    > anomala_h2.enrichedGeneIDs
cat bruxellensis.ratio.tsv | sed 's/^.*%\sG/G/' | sed 's/G/g/g' \
    | sed 's/\s/\n/g' | sort | uniq > bruxellensis.enrichedGeneIDs
cat custersianus.ratio.tsv | sed 's/^.*%\sG/G/' | sed 's/G/g/g' \
    | sed 's/\s/\n/g' | sort | uniq > custersianus.enrichedGeneIDs
cat naardenensis.ratio.tsv | sed 's/^.*%\sG/G/' | sed 's/G/g/g' \
    | sed 's/\s/\n/g' | sort | uniq > naardenensis.enrichedGeneIDs
cat nanus.ratio.tsv | sed 's/^.*%\sG/G/' | sed 's/G/g/g' \
    | sed 's/\s/\n/g' | sort | uniq > nanus.enrichedGeneIDs

# Copy to SQL directory and pull the KEGG annotations for the GO-enriched genes
cp *enrichedGeneIDs ../../functional-annotation/DB/
```

```
# Get the OG IDs for the enriched genes, cross-check for consistency with the
# original OG-enrichment analysis (Example for anomalus Haplome 1)
sqlite> create table anomH1enrichedGenes (gene TEXT not null);
# etc…

# Import enriched gene IDs
sqlite> .import anomala_h1.enrichedGeneIDs anomH1enrichedGenes
# etc…

# pull the OG IDs for the enriched gene IDs, backup to TSV files,
# check against original OG-enrichments
sqlite> .out anomalaH1.enrichedOGs
sqlite> select distinct orthogroup from annotations a
    inner join anomH1enrichedGenes b where a.gene == b.gene and
    a.species like 'Brettanomyces_anomala_h1';
sqlite> .out stdout
# etc…

# Pull the gene IDs for the OGs with enriched GO-terms
sqlite> create table anomH1enrichedOGs (og TEXT not null);
# etc…

# import
sqlite> .import anomalaH1.enrichedOGs anomH1enrichedOGs
# etc…

# Pull the gene IDs
sqlite> .out anomalaH1.GO-enriched.KEGG
sqlite> select k.Ko, k.An, k.Bn, K.Cn, k.Kn from annotations a
    inner join anomH1enrichedOGs b on b.og = a.orthogroup
    inner join kegg k on k.Ko = a.kegg
        where a.species like 'Brettanomyces_anomala_h1%' and
        a.kegg != '.' and k.An != "Organismal Systems" and
        k.An != "Human Diseases" and k.An != "Brite Hierarchies"
    ORDER by K.An, K.Bn, K.Cn, k.Kn;
sqlite> .out stdout
# etc...

# perform alignments and make gene trees
mkdir genePhylogenies && cd genePhylogenies/

# Check what OGs have each KEGG annotation
sqlite> select * from annotations where kegg == 'K10967' and
    species like "Brett%";

# Check if there are any other genes in each OG without KEGG annotations
# (not always annotated perfectly, or exactly the same).
sqlite> select * from annotations where orthogroup=='OG0000020' and
    species like "Brett%";
# etc...

# Get the gene seqs for each OG
# used a custom script to retrieve the gene seqs, manually made a list of
# the Brettanomyces species and the OGs
# link the orthofinder TSV for convenience
ln -s ../../functional-annotation/orthofinder/Results_Nov15/Orthogroups.csv
```

```
# run the script
perl OG-species-tree.pl ../../functional-annotation/orthofinder/ \
    Orthogroups.csv species.list ortho.list

# perform MSA with muscle
# concatenate OGs 4127 with 1589 as these are to be analysed together
cat OG0001589.faa OG0004127.faa > OG0001589.OG0004127.faa

# rename old files (so they're not processed)
mv OG0001589.faa _0001589.faa
mv OG0004127.faa _0004127.faa

# do the alignments
for i in `ls | grep OG`; do
    muscle -in $i > $i.MSA;
done

# Generate gene trees
      # performed manually in seaview

# Inspect gene trees and alignments, note any broken/truncated models
# here and filter excel table
      # OG0000023: nanus: g2048 - truncated
      # OG0000073: nanus: g2092 - incomplete
      # OG0000107: bruxellensis: g995 g996 - fragmented model.
      # OG0000227: anomala_h2: g4317 g4318 g4319 - fragmented model. PRUNE OG
      # OG0000418: anomala_h1h2: almost all models incomplete. PRUNE OG
      # OG0000496: custersianus: g3867 - truncated. PRUNE OG
      # OG0005135: bruxellensis: g4001 - truncated
            # anomala_h2: g10 g11 - truncated
```

#### Adaptive selection analysis

```
# Link protein and transcript sequences
ln -s ../assembly-gene-annotation/AWRI2804.prot.fasta \
    bruxellensis.proteins.fasta
ln -s ../assembly-gene-annotation/AWRI2804.transcripts.fasta \
    bruxellensis.transcripts.fasta
ln -s ../assembly-gene-annotation/B.anomala.prot.fasta \
    anomala.proteins.fasta
ln -s ../assembly-gene-annotation/B.anomala.transcripts.fasta \
    anomala.transcripts.fasta
ln -s ../assembly-gene-annotation/B.custersianus.prot.fasta \
    custersianus.proteins.fasta
ln -s ../assembly-gene-annotation/B.custersianus.transcripts.fasta \
    custersianus.transcripts.fasta
ln -s ../assembly-gene-annotation/B.naardenensis.prot.fasta \
    naardenensis.proteins.fasta
ln -s ../assembly-gene-annotation/B.naardenensis.transcripts.fasta \
    naardenensis.transcripts.fasta
ln -s ../assembly-gene-annotation/B.nanus.prot.fasta \
    nanus.proteins.fasta
ln -s ../assembly-gene-annotation/B.nanus.transcripts.fasta \
    nanus.transcripts.fasta

# link orthofinder results (Brettanomyces, not Saccharomycetaceae)
ln -s ../assembly-gene-annotation/orthofinder/Results_Mar29/Orthogroups.csv

# use only single-copy orthologues
cat Orthogroups.csv | grep -v , | grep -v -P "\s\s" > Orthogroups.SCOs.csv
```

```
# protein alignments and generate transcript multifastas
perl pull_and_align_SCOs.pl Orthogroups.SCOs.csv MSAs

# create the codon alignments
for i in `ls | grep MSA | grep -v prot | sed 's/\.MSA\.fa//'`; do
    pal2nal.pl $i.MSA.fa $i.t.fa -output fasta > $i.cdaln.fa;
done

# Manually create sample list
nano samples.list
    # bruxellensis\ncustersianus\nanomala...

# Combine MSAs and create a species tree
ls | grep MSA.fa > prot.fofn
perl combineMSAs.pl prot.fofn samples.list > all.prot.MSA.fa
    # created a protein tree in seaview with phyML + defaults

# Need a FOFN of the codon alignments
ls | grep cdaln > align.fofn

# Site based positive selection test (Example for M1a vs M2a models)
# run codeml using custom script
perl multiCodeml.pl align.fofn template.ctl

# tidy up
mkdir M1aM2a/
mv OG*M1aM2a M1aM2a/

# scan output
find M1aM2a/ -name "OG*M1aM2a" > M1aM2a.fofn
perl codemlOutputScan.pl M1aM2a.fofn | sort -k2,2n > M1aM2a.results
for i in `cat M1aM2a.results | awk '{print $1}'`; do
    grep $i ../Orthogroups.SCOs.csv ;
done | awk '{print $2}' > M1aM2a.posSel.brux.genes.lst
```

#### Horizontal Gene Transfer check

```
# blastp proteins for each species to refseq fungi and bact datasets
for i in Brettanomyces_anomala_h1 Brettanomyces_anomala_h2 \
         Brettanomyces_bruxellensis Brettanomyces_custersianus \
         Brettanomyces_naardenensis Brettanomyces_nanus; do

    # blastp against refseq fungal dataset
    cat ../$i.faa \
        | parallel --block 100k --recstart '>' -j 16 --pipe \
          blastp -query - -db /scratch/refseq/fungi \
          -outfmt \"6 qseqid sseqid qlen slen length evalue bitscore\" \
          -evalue 1e-20 -task blastp-fast -num_alignments 1 \
        | awk '$5/$3>=0.5{print}' \
        | gzip - > $i.fungi.gz

    # blastp against refseq bacterial dataset
    cat ../$i.faa \
        | parallel --block 100k --recstart '>' -j 8 --pipe \
          blastp -query - -db /scratch/refseq/bact \
          -outfmt \"6 qseqid sseqid qlen slen length evalue bitscore\" \
          -evalue 1e-20 -task blastp-fast -num_alignments 1 \
        | awk '$5/$3>=0.5{print}' \
        | gzip - > $i.bact.gz

    # retrieve the candidates
    perl HGT-check.pl B.bruxellensis.fungi.gz B.bruxellensis.bact.gz \
        > B.bruxellensis.HGT.tsv
done

# pull the OGs from the SQL database for these candidates
cp *HGT.tsv ../DB/

# example for B. anomalus haplome 1 only
sqlite> create table anomH1HGT (gene TEXT, eval1 TEXT, eval2 TEXT);
sqlite> .import B.anomala.h1.HGT.tsv anomH1HGT
sqlite> select orthogroup from annotations a
    inner join anomH1HGT b where b.gene == a.gene and
    a.species like "Brettanomyces_anomala_h1";
      # copy-pasted to file HGT.list

# sort and retrieve unique OGs
cat HGT.list | sort | uniq > HGT.uniq.list

# manually inspect the trees for these OGs
```
