## Supplementary figures and images for "New genome assemblies reveal patterns of domestication and adaptation across *Brettanomyces* (*Dekkera*) species"

### Figure S1

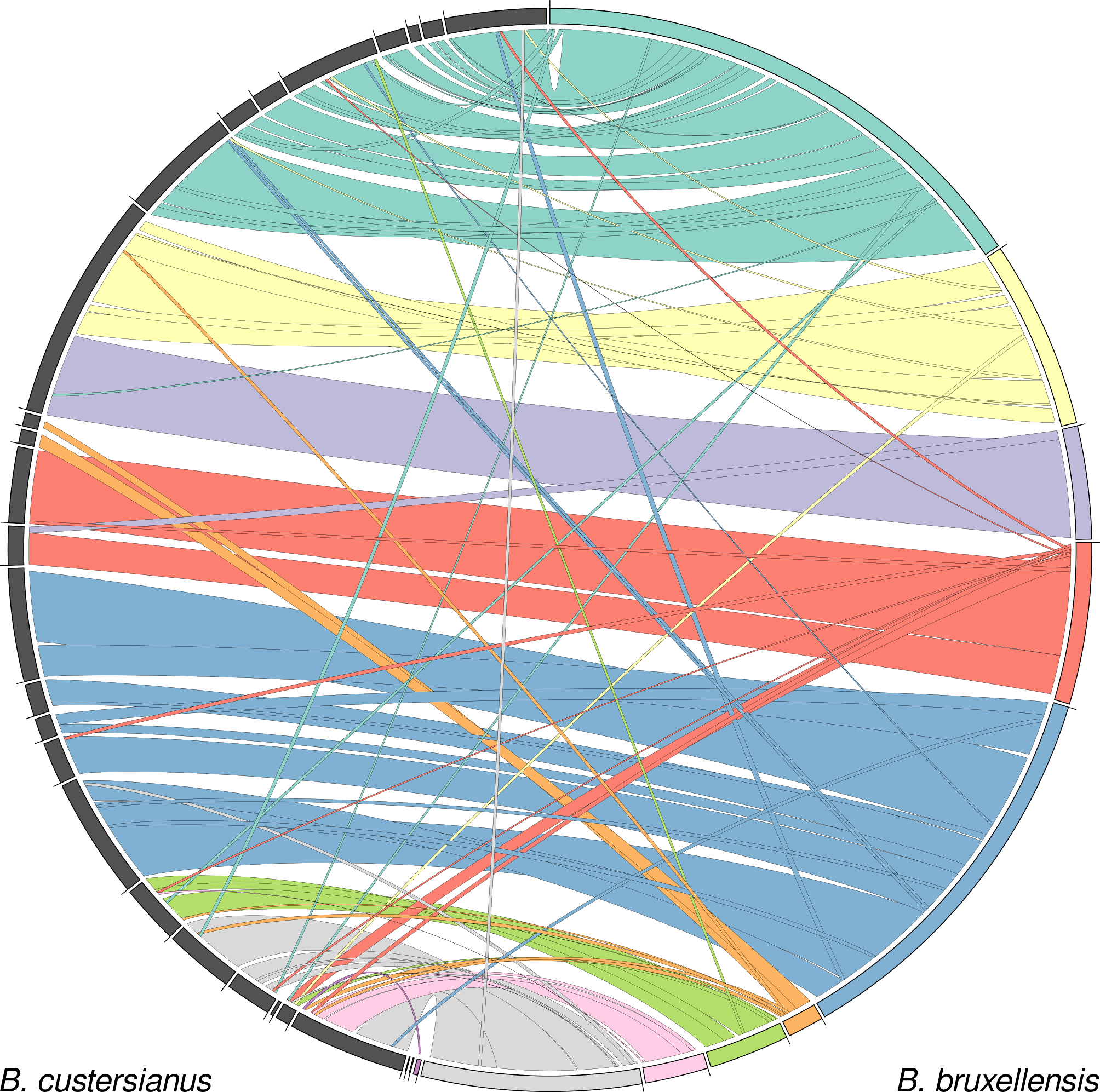

### Figure S2

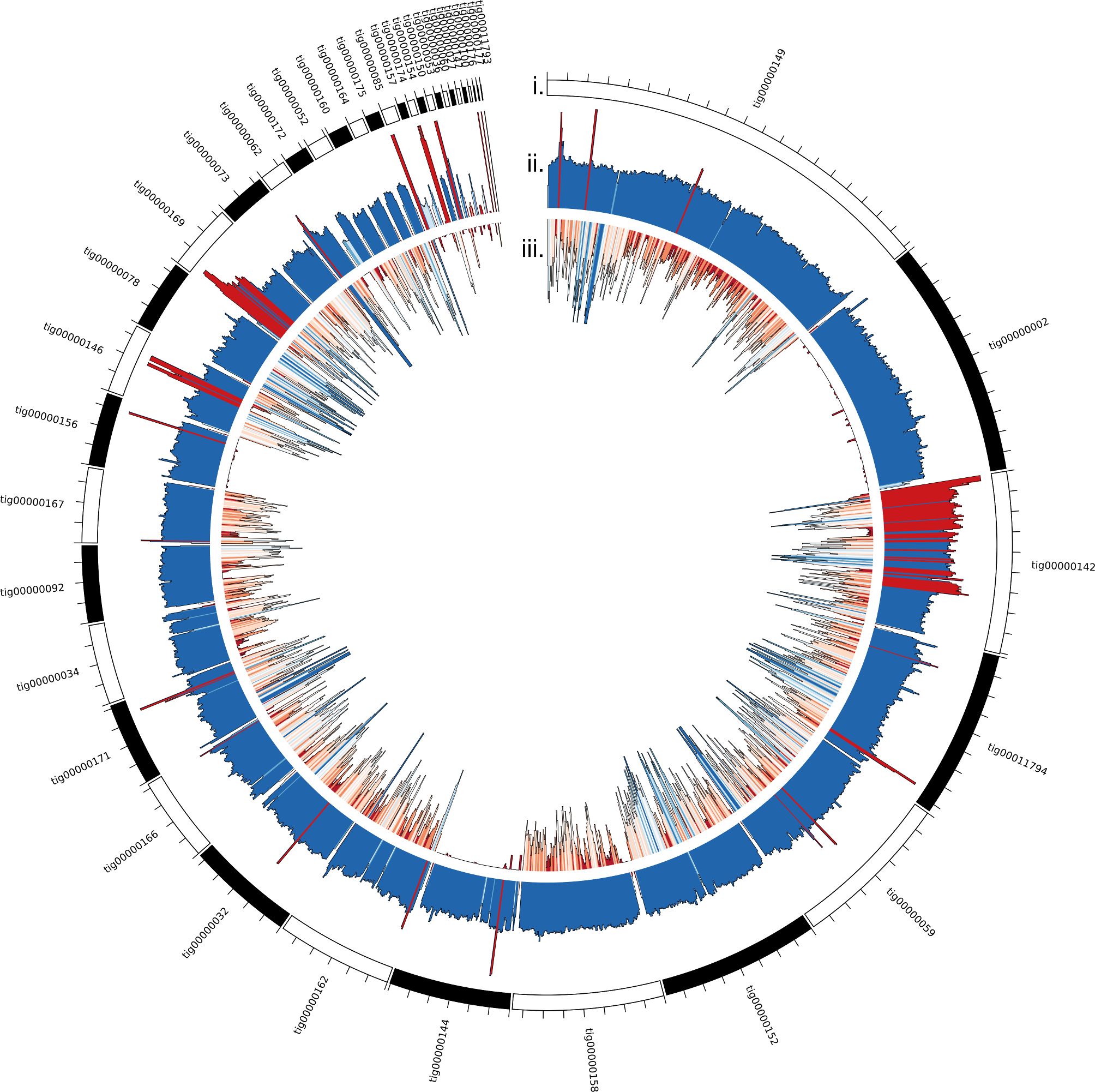
