## Supplementary material for "New genome assemblies reveal patterns of domestication and adaptation across *Brettanomyces* (*Dekkera*) species": Tables S1-4

### Table S1: MinION sequencing metrics for *Brettanomyces* sequencing.

| | **Median read length (bp)** | **Median read quality (QV)** | **Number of reads** | **Read length N50 (bp)** | **Total bases (Gb)** |
|---|---|---|---|---|---|
| ***B. anomalus*** | 4 685 | 8.7 | 296 479 | 7 859 | 1.65 |
| ***B. custersianus*** | 6 695 | 7.9 | 180 237 | 10 588 | 1.35 |
| ***B. naardenensis*** | 4 249 | 8.9 | 83 581 | 9 336 | 0.497 |
| ***B. nanus*** | 14 912 | 10.8 | 28 586 | 30 870 | 0.508 |

### Table S2: Predicted genes and gene density for the *Brettanomyces* genomes

| | **Number of genes** | **Total genic sequence (bp)** | **Total genic sequence (% of genome)** |
|---|---|---|---|
| ***B. anomalus*** | 5 735 | 8 571 724 | 62.2 |
| ***B. bruxellensis*** | 5 293 | 8 469 775 | 64.2 |
| ***B. custersianus*** | 5 255 | 8 094 938 | 75.4 |
| ***B. naardenensis*** | 5 334 | 8 393 268 | 75.2 |
| ***B. nanus*** | 5 083 | 7 960 011 | 78.1 |
| ***S. cerevisiae* (S288C)** | 6 445 | 9 008 924 | 74.1 |

Table S3: KEGG-annotated genes under site selection across *Brettanomyces*

| KEGG ID | CATEGORY | SUB-CATEGORY | PATHWAY | GENE |
|---|---|---|---|---|
| **K07827** | Cellular Processes | Cell growth and death | Apoptosis | GTPase KRas |
| **K07827** | Cellular Processes | Cell growth and death | Apoptosis - fly | GTPase KRas |
| **K08334** | Cellular Processes | Cell growth and death | Apoptosis - multiple species | beclin |
| **K02541** | Cellular Processes | Cell growth and death | Cell cycle | DNA replication licensing factor MCM3 |
| **K02210** | Cellular Processes | Cell growth and death | Cell cycle | DNA replication licensing factor MCM7 |
| **K06067** | Cellular Processes | Cell growth and death | Cell cycle | histone deacetylase 1/2 |
| **K02603** | Cellular Processes | Cell growth and death | Cell cycle | origin recognition complex subunit 1 |
| **K02604** | Cellular Processes | Cell growth and death | Cell cycle | origin recognition complex subunit 2 |
| **K08866** | Cellular Processes | Cell growth and death | Cell cycle | serine/threonine-protein kinase TTK/MPS1 |
| **K02541** | Cellular Processes | Cell growth and death | Cell cycle - yeast | DNA replication licensing factor MCM3 |
| **K02210** | Cellular Processes | Cell growth and death | Cell cycle - yeast | DNA replication licensing factor MCM7 |
| **K10259** | Cellular Processes | Cell growth and death | Cell cycle - yeast | F-box and WD-40 domain protein MET30 |
| **K02220** | Cellular Processes | Cell growth and death | Cell cycle - yeast | G2/mitotic-specific cyclin 1/2 |
| **K06676** | Cellular Processes | Cell growth and death | Cell cycle - yeast | condensin complex subunit 2 |
| **K06678** | Cellular Processes | Cell growth and death | Cell cycle - yeast | condensin complex subunit 3 |
| **K06666** | Cellular Processes | Cell growth and death | Cell cycle - yeast | general transcriptional corepressor TUP1 |
| **K02603** | Cellular Processes | Cell growth and death | Cell cycle - yeast | origin recognition complex subunit 1 |
| **K02604** | Cellular Processes | Cell growth and death | Cell cycle - yeast | origin recognition complex subunit 2 |
| **K08866** | Cellular Processes | Cell growth and death | Cell cycle - yeast | serine/threonine-protein kinase TTK/MPS1 |
| **K07827** | Cellular Processes | Cell growth and death | Cellular senescence | GTPase KRas |
| **K04441** | Cellular Processes | Cell growth and death | Cellular senescence | p38 MAP kinase |
| **K06269** | Cellular Processes | Cell growth and death | Cellular senescence | serine/threonine-protein phosphatase PP1 catalytic subunit |
| **K05863** | Cellular Processes | Cell growth and death | Cellular senescence | solute carrier family 25 (mitochondrial adenine nucleotide translocator), member 4/5/6/31 |
| **K08337** | Cellular Processes | Cell growth and death | Ferroptosis | ubiquitin-like modifier-activating enzyme ATG7 |
| **K02541** | Cellular Processes | Cell growth and death | Meiosis - yeast | DNA replication licensing factor MCM3 |
| **K02210** | Cellular Processes | Cell growth and death | Meiosis - yeast | DNA replication licensing factor MCM7 |
| **K02603** | Cellular Processes | Cell growth and death | Meiosis - yeast | origin recognition complex subunit 1 |
| **K02604** | Cellular Processes | Cell growth and death | Meiosis - yeast | origin recognition complex subunit 2 |
| **K06269** | Cellular Processes | Cell growth and death | Meiosis - yeast | serine/threonine-protein phosphatase PP1 catalytic subunit |
| **K11251** | Cellular Processes | Cell growth and death | Necroptosis | histone H2A |

| | | | | |
|---|---|---|---|---|
| **K05863** | Cellular Processes | Cell growth and death | Necroptosis | solute carrier family 25 (mitochondrial adenine nucleotide translocator), member 4/5/6/31 |
| **K06269** | Cellular Processes | Cell growth and death | Oocyte meiosis | serine/threonine-protein phosphatase PP1 catalytic subunit |
| **K07827** | Cellular Processes | Cell motility | Regulation of actin cytoskeleton | GTPase KRas |
| **K06269** | Cellular Processes | Cell motility | Regulation of actin cytoskeleton | serine/threonine-protein phosphatase PP1 catalytic subunit |
| **K03097** | Cellular Processes | Cellular community - eukaryotes | Adherens junction | casein kinase II subunit alpha |
| **K06269** | Cellular Processes | Cellular community - eukaryotes | Focal adhesion | serine/threonine-protein phosphatase PP1 catalytic subunit |
| **K07827** | Cellular Processes | Cellular community - eukaryotes | Gap junction | GTPase KRas |
| **K07827** | Cellular Processes | Cellular community - eukaryotes | Signaling pathways regulating pluripotency of stem cells | GTPase KRas |
| **K04441** | Cellular Processes | Cellular community - eukaryotes | Signaling pathways regulating pluripotency of stem cells | p38 MAP kinase |
| **K10591** | Cellular Processes | Cellular community - eukaryotes | Tight junction | E3 ubiquitin-protein ligase NEDD4 |
| **K09489** | Cellular Processes | Cellular community - eukaryotes | Tight junction | heat shock 70kDa protein 4 |
| **K07827** | Cellular Processes | Transport and catabolism | Autophagy - animal | GTPase KRas |
| **K08334** | Cellular Processes | Transport and catabolism | Autophagy - animal | beclin |
| **K08337** | Cellular Processes | Transport and catabolism | Autophagy - animal | ubiquitin-like modifier-activating enzyme ATG7 |
| **K08334** | Cellular Processes | Transport and catabolism | Autophagy - other | beclin |
| **K08337** | Cellular Processes | Transport and catabolism | Autophagy - other | ubiquitin-like modifier-activating enzyme ATG7 |
| **K07827** | Cellular Processes | Transport and catabolism | Autophagy - yeast | GTPase KRas |
| **K06902** | Cellular Processes | Transport and catabolism | Autophagy - yeast | MFS transporter, UMF1 family |
| **K08334** | Cellular Processes | Transport and catabolism | Autophagy - yeast | beclin |
| **K08337** | Cellular Processes | Transport and catabolism | Autophagy - yeast | ubiquitin-like modifier-activating enzyme ATG7 |
| **K12493** | Cellular Processes | Transport and catabolism | Endocytosis | ADP-ribosylation factor GTPase-activating protein 2/3 |
| **K10591** | Cellular Processes | Transport and catabolism | Endocytosis | E3 ubiquitin-protein ligase NEDD4 |
| **K18442** | Cellular Processes | Transport and catabolism | Endocytosis | brefeldin A-inhibited guanine nucleotide-exchange protein |
| **K12481** | Cellular Processes | Transport and catabolism | Endocytosis | rabenosyn-5 |
| **K18467** | Cellular Processes | Transport and catabolism | Endocytosis | vacuolar protein sorting-associated protein 29 |
| **K01052** | Cellular Processes | Transport and catabolism | Lysosome | lysosomal acid lipase/cholesteryl ester hydrolase |
| **K07827** | Cellular Processes | Transport and catabolism | Mitophagy - animal | GTPase KRas |
| **K08334** | Cellular Processes | Transport and catabolism | Mitophagy - animal | beclin |
| **K03097** | Cellular Processes | Transport and catabolism | Mitophagy - animal | casein kinase II subunit alpha |
| **K08955** | Cellular Processes | Transport and catabolism | Mitophagy - yeast | ATP-dependent metalloprotease |
| **K03097** | Cellular Processes | Transport and catabolism | Mitophagy - yeast | casein kinase II subunit alpha |

| | | | | |
|---|---|---|---|---|
| **K04441** | Cellular Processes | Transport and catabolism | Mitophagy - yeast | p38 MAP kinase |
| **K11644** | Cellular Processes | Transport and catabolism | Mitophagy - yeast | paired amphipathic helix protein Sin3a |
| **K19029** | Environmental Information Processing | Signal transduction | AMPK signaling pathway | 6-phosphofructo-2-kinase / fructose-2,6-biphosphatase 2 |
| **K07827** | Environmental Information Processing | Signal transduction | Apelin signaling pathway | GTPase KRas |
| **K08334** | Environmental Information Processing | Signal transduction | Apelin signaling pathway | beclin |
| **K05863** | Environmental Information Processing | Signal transduction | Calcium signaling pathway | solute carrier family 25 (mitochondrial adenine nucleotide translocator), member 4/5/6/31 |
| **K07827** | Environmental Information Processing | Signal transduction | ErbB signaling pathway | GTPase KRas |
| **K07827** | Environmental Information Processing | Signal transduction | FoxO signaling pathway | GTPase KRas |
| **K04441** | Environmental Information Processing | Signal transduction | FoxO signaling pathway | p38 MAP kinase |
| **K03259** | Environmental Information Processing | Signal transduction | HIF-1 signaling pathway | translation initiation factor 4E |
| **K06269** | Environmental Information Processing | Signal transduction | Hippo signaling pathway | serine/threonine-protein phosphatase PP1 catalytic subunit |
| **K02218** | Environmental Information Processing | Signal transduction | Hippo signaling pathway - multiple species | casein kinase 1 |
| **K07827** | Environmental Information Processing | Signal transduction | MAPK signaling pathway | GTPase KRas |
| **K04441** | Environmental Information Processing | Signal transduction | MAPK signaling pathway | p38 MAP kinase |
| **K07827** | Environmental Information Processing | Signal transduction | MAPK signaling pathway - fly | GTPase KRas |
| **K04441** | Environmental Information Processing | Signal transduction | MAPK signaling pathway - fly | p38 MAP kinase |
| **K04508** | Environmental Information Processing | Signal transduction | MAPK signaling pathway - fly | transducin (beta)-like 1 |
| **K10591** | Environmental Information Processing | Signal transduction | MAPK signaling pathway - yeast | E3 ubiquitin-protein ligase NEDD4 |
| **K02220** | Environmental Information Processing | Signal transduction | MAPK signaling pathway - yeast | G2/mitotic-specific cyclin 1/2 |
| **K02218** | Environmental Information Processing | Signal transduction | MAPK signaling pathway - yeast | casein kinase 1 |
| **K06666** | Environmental Information Processing | Signal transduction | MAPK signaling pathway - yeast | general transcriptional corepressor TUP1 |
| **K04441** | Environmental Information Processing | Signal transduction | MAPK signaling pathway - yeast | p38 MAP kinase |
| **K19704** | Environmental Information Processing | Signal transduction | MAPK signaling pathway - yeast | protein phosphatase PTC1 |
| **K19833** | Environmental Information Processing | Signal transduction | MAPK signaling pathway - yeast | serine/threonine-protein kinase CLA4 |
| **K03097** | Environmental Information Processing | Signal transduction | NF-kappa B signaling pathway | casein kinase II subunit alpha |
| **K06067** | Environmental Information Processing | Signal transduction | Notch signaling pathway | histone deacetylase 1/2 |
| **K07827** | Environmental Information Processing | Signal transduction | PI3K-Akt signaling pathway | GTPase KRas |
| **K03259** | Environmental Information Processing | Signal transduction | PI3K-Akt signaling pathway | translation initiation factor 4E |
| **K19801** | Environmental Information Processing | Signal transduction | Phosphatidylinositol signaling system | phosphatidylinositol 4-kinase B |
| **K07827** | Environmental Information Processing | Signal transduction | Phospholipase D signaling pathway | GTPase KRas |
| **K07827** | Environmental Information Processing | Signal transduction | Rap1 signaling pathway | GTPase KRas |
| **K04441** | Environmental Information Processing | Signal transduction | Rap1 signaling pathway | p38 MAP kinase |
| **K07827** | Environmental Information Processing | Signal transduction | Ras signaling pathway | GTPase KRas |

| | | | | |
|---|---|---|---|---|
| **K07827** | Environmental Information Processing | Signal transduction | Sphingolipid signaling pathway | GTPase KRas |
| **K04441** | Environmental Information Processing | Signal transduction | Sphingolipid signaling pathway | p38 MAP kinase |
| **K12351** | Environmental Information Processing | Signal transduction | Sphingolipid signaling pathway | sphingomyelin phosphodiesterase 2 |
| **K04441** | Environmental Information Processing | Signal transduction | TNF signaling pathway | p38 MAP kinase |
| **K07827** | Environmental Information Processing | Signal transduction | VEGF signaling pathway | GTPase KRas |
| **K04441** | Environmental Information Processing | Signal transduction | VEGF signaling pathway | p38 MAP kinase |
| **K03097** | Environmental Information Processing | Signal transduction | Wnt signaling pathway | casein kinase II subunit alpha |
| **K04508** | Environmental Information Processing | Signal transduction | Wnt signaling pathway | transducin (beta)-like 1 |
| **K06269** | Environmental Information Processing | Signal transduction | cAMP signaling pathway | serine/threonine-protein phosphatase PP1 catalytic subunit |
| **K06269** | Environmental Information Processing | Signal transduction | cGMP-PKG signaling pathway | serine/threonine-protein phosphatase PP1 catalytic subunit |
| **K05863** | Environmental Information Processing | Signal transduction | cGMP-PKG signaling pathway | solute carrier family 25 (mitochondrial adenine nucleotide translocator), member 4/5/6/31 |
| **K20404** | Environmental Information Processing | Signal transduction | mTOR signaling pathway | DEP domain-containing protein 5 |
| **K07827** | Environmental Information Processing | Signal transduction | mTOR signaling pathway | GTPase KRas |
| **K03259** | Environmental Information Processing | Signal transduction | mTOR signaling pathway | translation initiation factor 4E |
| **K02737** | Genetic Information Processing | Folding, sorting and degradation | Proteasome | 20S proteasome subunit beta 5 |
| **K03028** | Genetic Information Processing | Folding, sorting and degradation | Proteasome | 26S proteasome regulatory subunit N1 |
| **K03029** | Genetic Information Processing | Folding, sorting and degradation | Proteasome | 26S proteasome regulatory subunit N10 |
| **K03039** | Genetic Information Processing | Folding, sorting and degradation | Proteasome | 26S proteasome regulatory subunit N9 |
| **K09486** | Genetic Information Processing | Folding, sorting and degradation | Protein processing in endoplasmic reticulum | hypoxia up-regulated 1 |
| **K14003** | Genetic Information Processing | Folding, sorting and degradation | Protein processing in endoplasmic reticulum | prolactin regulatory element-binding protein |
| **K12581** | Genetic Information Processing | Folding, sorting and degradation | RNA degradation | CCR4-NOT transcription complex subunit 7/8 |
| **K12606** | Genetic Information Processing | Folding, sorting and degradation | RNA degradation | CCR4-NOT transcription complex subunit 9 |
| **K04077** | Genetic Information Processing | Folding, sorting and degradation | RNA degradation | chaperonin GroEL |
| **K12592** | Genetic Information Processing | Folding, sorting and degradation | RNA degradation | exosome complex protein LRP1 |
| **K13126** | Genetic Information Processing | Folding, sorting and degradation | RNA degradation | polyadenylate-binding protein |
| **K10591** | Genetic Information Processing | Folding, sorting and degradation | Ubiquitin mediated proteolysis | E3 ubiquitin-protein ligase NEDD4 |
| **K10259** | Genetic Information Processing | Folding, sorting and degradation | Ubiquitin mediated proteolysis | F-box and WD-40 domain protein MET30 |
| **K03648** | Genetic Information Processing | Replication and repair | Base excision repair | uracil-DNA glycosylase |
| **K02541** | Genetic Information Processing | Replication and repair | DNA replication | DNA replication licensing factor MCM3 |
| **K02210** | Genetic Information Processing | Replication and repair | DNA replication | DNA replication licensing factor MCM7 |
| **K08735** | Genetic Information Processing | Replication and repair | Mismatch repair | DNA mismatch repair protein MSH2 |
| **K08737** | Genetic Information Processing | Replication and repair | Mismatch repair | DNA mismatch repair protein MSH6 |
| **K03142** | Genetic Information Processing | Replication and repair | Nucleotide excision repair | transcription initiation factor TFIIH subunit 2 |

| | | | | |
|---|---|---|---|---|
| **K03143** | Genetic Information Processing | Replication and repair | Nucleotide excision repair | transcription initiation factor TFIIH subunit 3 |
| **K03142** | Genetic Information Processing | Transcription | Basal transcription factors | transcription initiation factor TFIIH subunit 2 |
| **K03143** | Genetic Information Processing | Transcription | Basal transcription factors | transcription initiation factor TFIIH subunit 3 |
| **K03006** | Genetic Information Processing | Transcription | RNA polymerase | DNA-directed RNA polymerase II subunit RPB1 |
| **K03011** | Genetic Information Processing | Transcription | RNA polymerase | DNA-directed RNA polymerase II subunit RPB3 |
| **K12881** | Genetic Information Processing | Transcription | Spliceosome | THO complex subunit 4 |
| **K12817** | Genetic Information Processing | Transcription | Spliceosome | pre-mRNA-splicing factor 18 |
| **K12837** | Genetic Information Processing | Transcription | Spliceosome | splicing factor U2AF 65 kDa subunit |
| **K01870** | Genetic Information Processing | Translation | Aminoacyl-tRNA biosynthesis | isoleucyl-tRNA synthetase |
| **K01869** | Genetic Information Processing | Translation | Aminoacyl-tRNA biosynthesis | leucyl-tRNA synthetase |
| **K12881** | Genetic Information Processing | Translation | RNA transport | THO complex subunit 4 |
| **K13126** | Genetic Information Processing | Translation | RNA transport | polyadenylate-binding protein |
| **K03242** | Genetic Information Processing | Translation | RNA transport | translation initiation factor 2 subunit 3 |
| **K03245** | Genetic Information Processing | Translation | RNA transport | translation initiation factor 3 subunit J |
| **K03259** | Genetic Information Processing | Translation | RNA transport | translation initiation factor 4E |
| **K03260** | Genetic Information Processing | Translation | RNA transport | translation initiation factor 4G |
| **K03243** | Genetic Information Processing | Translation | RNA transport | translation initiation factor 5B |
| **K02885** | Genetic Information Processing | Translation | Ribosome | large subunit ribosomal protein L19e |
| **K02891** | Genetic Information Processing | Translation | Ribosome | large subunit ribosomal protein L22e |
| **K02892** | Genetic Information Processing | Translation | Ribosome | large subunit ribosomal protein L23 |
| **K02893** | Genetic Information Processing | Translation | Ribosome | large subunit ribosomal protein L23Ae |
| **K02921** | Genetic Information Processing | Translation | Ribosome | large subunit ribosomal protein L37Ae |
| **K02930** | Genetic Information Processing | Translation | Ribosome | large subunit ribosomal protein L4e |
| **K02937** | Genetic Information Processing | Translation | Ribosome | large subunit ribosomal protein L7e |
| **K02955** | Genetic Information Processing | Translation | Ribosome | small subunit ribosomal protein S14e |
| **K02958** | Genetic Information Processing | Translation | Ribosome | small subunit ribosomal protein S15e |
| **K02962** | Genetic Information Processing | Translation | Ribosome | small subunit ribosomal protein S17e |
| **K07178** | Genetic Information Processing | Translation | Ribosome biogenesis in eukaryotes | RIO kinase 1 |
| **K11883** | Genetic Information Processing | Translation | Ribosome biogenesis in eukaryotes | RNA-binding protein NOB1 |
| **K03097** | Genetic Information Processing | Translation | Ribosome biogenesis in eukaryotes | casein kinase II subunit alpha |
| **K14538** | Genetic Information Processing | Translation | Ribosome biogenesis in eukaryotes | nuclear GTP-binding protein |
| **K14565** | Genetic Information Processing | Translation | Ribosome biogenesis in eukaryotes | nucleolar protein 58 |
| **K03685** | Genetic Information Processing | Translation | Ribosome biogenesis in eukaryotes | ribonuclease III |
| **K14536** | Genetic Information Processing | Translation | Ribosome biogenesis in eukaryotes | ribosome assembly protein 1 |

| | | | | |
|---|---|---|---|---|
| **K12881** | Genetic Information Processing | Translation | mRNA surveillance pathway | THO complex subunit 4 |
| **K14402** | Genetic Information Processing | Translation | mRNA surveillance pathway | cleavage and polyadenylation specificity factor subunit 2 |
| **K14416** | Genetic Information Processing | Translation | mRNA surveillance pathway | elongation factor 1 alpha-like protein |
| **K00565** | Genetic Information Processing | Translation | mRNA surveillance pathway | mRNA (guanine-N7-)-methyltransferase |
| **K13126** | Genetic Information Processing | Translation | mRNA surveillance pathway | polyadenylate-binding protein |
| **K14396** | Genetic Information Processing | Translation | mRNA surveillance pathway | polyadenylate-binding protein 2 |
| **K14400** | Genetic Information Processing | Translation | mRNA surveillance pathway | pre-mRNA cleavage complex 2 protein Pcf11 |
| **K06269** | Genetic Information Processing | Translation | mRNA surveillance pathway | serine/threonine-protein phosphatase PP1 catalytic subunit |
| **K00931** | Metabolism | Amino acid metabolism | Arginine and proline metabolism | glutamate 5-kinase |
| **K00818** | Metabolism | Amino acid metabolism | Arginine biosynthesis | acetylornithine aminotransferase |
| **K01738** | Metabolism | Amino acid metabolism | Cysteine and methionine metabolism | cysteine synthase |
| **K01079** | Metabolism | Amino acid metabolism | Glycine, serine and threonine metabolism | phosphoserine phosphatase |
| **K00143** | Metabolism | Amino acid metabolism | Lysine biosynthesis | L-2-aminoadipate reductase |
| **K00931** | Metabolism | Biosynthesis of other secondary metabolites | Carbapenem biosynthesis | glutamate 5-kinase |
| **K01183** | Metabolism | Carbohydrate metabolism | Amino sugar and nucleotide sugar metabolism | chitinase |
| **K00966** | Metabolism | Carbohydrate metabolism | Amino sugar and nucleotide sugar metabolism | mannose-1-phosphate guanylyltransferase |
| **K00469** | Metabolism | Carbohydrate metabolism | Ascorbate and aldarate metabolism | inositol oxygenase |
| **K01958** | Metabolism | Carbohydrate metabolism | Citrate cycle (TCA cycle) | pyruvate carboxylase |
| **K19029** | Metabolism | Carbohydrate metabolism | Fructose and mannose metabolism | 6-phosphofructo-2-kinase / fructose-2,6-biphosphatase 2 |
| **K00966** | Metabolism | Carbohydrate metabolism | Fructose and mannose metabolism | mannose-1-phosphate guanylyltransferase |
| **K01803** | Metabolism | Carbohydrate metabolism | Fructose and mannose metabolism | triosephosphate isomerase (TIM) |
| **K00873** | Metabolism | Carbohydrate metabolism | Glycolysis / Gluconeogenesis | pyruvate kinase |
| **K01803** | Metabolism | Carbohydrate metabolism | Glycolysis / Gluconeogenesis | triosephosphate isomerase (TIM) |
| **K00469** | Metabolism | Carbohydrate metabolism | Inositol phosphate metabolism | inositol oxygenase |
| **K19801** | Metabolism | Carbohydrate metabolism | Inositol phosphate metabolism | phosphatidylinositol 4-kinase B |
| **K01803** | Metabolism | Carbohydrate metabolism | Inositol phosphate metabolism | triosephosphate isomerase (TIM) |
| **K00948** | Metabolism | Carbohydrate metabolism | Pentose phosphate pathway | ribose-phosphate pyrophosphokinase |
| **K00616** | Metabolism | Carbohydrate metabolism | Pentose phosphate pathway | transaldolase |
| **K01958** | Metabolism | Carbohydrate metabolism | Pyruvate metabolism | pyruvate carboxylase |
| **K00873** | Metabolism | Carbohydrate metabolism | Pyruvate metabolism | pyruvate kinase |
| **K01803** | Metabolism | Energy metabolism | Carbon fixation in photosynthetic organisms | triosephosphate isomerase (TIM) |
| **K01958** | Metabolism | Energy metabolism | Carbon fixation pathways in prokaryotes | pyruvate carboxylase |
| **K01079** | Metabolism | Energy metabolism | Methane metabolism | phosphoserine phosphatase |

| | | | | |
|---|---|---|---|---|
| **K03953** | Metabolism | Energy metabolism | Oxidative phosphorylation | NADH dehydrogenase (ubiquinone) 1 alpha subcomplex subunit 9 |
| **K03935** | Metabolism | Energy metabolism | Oxidative phosphorylation | NADH dehydrogenase (ubiquinone) Fe-S protein 2 |
| **K01738** | Metabolism | Energy metabolism | Sulfur metabolism | cysteine synthase |
| **K05290** | Metabolism | Glycan biosynthesis and metabolism | Glycosylphosphatidylinositol (GPI)-anchor biosynthesis | phosphatidylinositol glycan, class K |
| **K10967** | Metabolism | Glycan biosynthesis and metabolism | Other types of O-glycan biosynthesis | alpha 1,2-mannosyltransferase |
| **K06123** | Metabolism | Lipid metabolism | Ether lipid metabolism | 1-acylglycerone phosphate reductase |
| **K00667** | Metabolism | Lipid metabolism | Fatty acid biosynthesis | fatty acid synthase subunit alpha, fungi type |
| **K18097** | Metabolism | Lipid metabolism | Glycerolipid metabolism | glycerol 2-dehydrogenase (NADP+) |
| **K06123** | Metabolism | Lipid metabolism | Glycerophospholipid metabolism | 1-acylglycerone phosphate reductase |
| **K00967** | Metabolism | Lipid metabolism | Glycerophospholipid metabolism | ethanolamine-phosphate cytidylyltransferase |
| **K04713** | Metabolism | Lipid metabolism | Sphingolipid metabolism | sphinganine C4-monooxygenase |
| **K12351** | Metabolism | Lipid metabolism | Sphingolipid metabolism | sphingomyelin phosphodiesterase 2 |
| **K00222** | Metabolism | Lipid metabolism | Steroid biosynthesis | Delta14-sterol reductase |
| **K01052** | Metabolism | Lipid metabolism | Steroid biosynthesis | lysosomal acid lipase/cholesteryl ester hydrolase |
| **K01012** | Metabolism | Metabolism of cofactors and vitamins | Biotin metabolism | biotin synthase |
| **K00560** | Metabolism | Metabolism of cofactors and vitamins | One carbon pool by folate | thymidylate synthase |
| **K01598** | Metabolism | Metabolism of cofactors and vitamins | Pantothenate and CoA biosynthesis | phosphopantothenoylcysteine decarboxylase |
| **K09680** | Metabolism | Metabolism of cofactors and vitamins | Pantothenate and CoA biosynthesis | type II pantothenate kinase |
| **K01719** | Metabolism | Metabolism of cofactors and vitamins | Porphyrin and chlorophyll metabolism | uroporphyrinogen-III synthase |
| **K00949** | Metabolism | Metabolism of cofactors and vitamins | Thiamine metabolism | thiamine pyrophosphokinase |
| **K01469** | Metabolism | Metabolism of other amino acids | Glutathione metabolism | 5-oxoprolinase (ATP-hydrolysing) |
| **K00967** | Metabolism | Metabolism of other amino acids | Phosphonate and phosphinate metabolism | ethanolamine-phosphate cytidylyltransferase |
| **K03006** | Metabolism | Nucleotide metabolism | Purine metabolism | DNA-directed RNA polymerase II subunit RPB1 |
| **K03011** | Metabolism | Nucleotide metabolism | Purine metabolism | DNA-directed RNA polymerase II subunit RPB3 |
| **K00873** | Metabolism | Nucleotide metabolism | Purine metabolism | pyruvate kinase |
| **K00948** | Metabolism | Nucleotide metabolism | Purine metabolism | ribose-phosphate pyrophosphokinase |
| **K03006** | Metabolism | Nucleotide metabolism | Pyrimidine metabolism | DNA-directed RNA polymerase II subunit RPB1 |
| **K03011** | Metabolism | Nucleotide metabolism | Pyrimidine metabolism | DNA-directed RNA polymerase II subunit RPB3 |
| **K00560** | Metabolism | Nucleotide metabolism | Pyrimidine metabolism | thymidylate synthase |

Table S4: Saccharomycetaceae species used with *Brettanomyces* species in OrthoFinder

| **Species** | **Genome accession** |
|---|---|
| *Agaricus bisporus* | GCF_000300575.1 |
| *Babjeviella inositovora* | GCF_001661335.1 |
| *Candida albicans* | GCF_000182965.3 |
| *Candida boidinii* | GCA_001599335.1 |
| *Candida glabrata* | GCF_000002545.3 |
| *Candida tenuis* | GCF_000002545.3 |
| *Citeromyces matritensis* | GCA_003243085.1 |
| *Debaryomyces hansenii* | GCF_000006445.2 |
| *Eremothecium gossypii* | GCF_000091025.4 |
| *Hansenisapora osmophila* | GCA_001747045.1 |
| *Kazachstania africana* | GCF_000304475.1 |
| *Kluveromyces lactis* | GCF_000002515.2 |
| *Komagataella phaffii* | GCF_000027005.1 |
| *Lachancea thermotolerans* | GCF_000142805.1 |
| *Lodderomyces elongisporus* | GCF_000149685.1 |
| *Nadsonia fulvescens* | GCA_001661315.1 |
| *Nakaseomyces bscillisporus* | GCA_001046975.1 |
| *Naumovozyma dairenensis* | GCF_000227115.2 |
| *Ogataea methanolica* | GCA_001600755.1 |
| *Ogataea parapolymorpha* | GCF_000187245.1 |
| *Ogataea polymorpha* | GCF_001664045.1 |
| *Pachysolen tannophilus* | GCA_001661245.1 |
| *Pichia membranifaciens* | GCF_001661235.1 |
| *Saccharomyces bayanus* | GCA_000167035.1 |
| *Saccharomyces boulardii* | GCA_001413975.1 |
| *Saccharomyces cerevisiae* | GCF_000146045.2 |
| *Saccharomyces eubayanus* | GCF_001298625.1 |
| *Saccharomyces mikatae* | GCA_000166975.1 |
| *Saccharomyces paradoxus* | GCA_002079145.1 |
| *Saccharomyces uvarum* | GCA_002242645.1 |
| *Spathaspora xylofermentans* | GCA_002105455.1 |
| *Torulaspora delbrueckii* | GCF_000243375.1 |
| *Wickerhamiella sorbophila* | GCF_002251995.1 |
| *Wickerhamomyces anomalus* | GCF_001661255.1 |
| *Wickerhamomyces ciferrii* | GCF_000313485.1 |
| *Zygosaccharomyces parabailii* | GCA_001984395.2 |
